# Mice navigate complex landscapes by learning a sparse vocabulary of paths

**DOI:** 10.64898/2026.10.10.757726

**Authors:** Tzuhsuan Ma, Alison E Comrie, Vojko Pjanovic, Lili Karashchuk, Emily J Dennis, Ann M Hermundstad, Jakob Voigts

**Affiliations:** HHMI Janelia Research Campus, Ashburn, VA, USA

## Abstract

Effective navigation is critical for survival, but how animals contend with the scale, complexity, and self-similarity of natural landscapes is unknown. Here, we investigate spontaneous exploratory mouse behavior, both outdoors and in dark self-similar landscapes made from tens to hundreds of tiles of varying heights. Within 2 hours of exploration, mice detected small covert changes to the landscape and jumped between towers to invisible landing spots, revealing detailed spatial knowledge. In this process, mice converged on a sparse set of memorized paths. These routes depended on prior experience and increased the compressibility of sensory information over time. Our results demonstrate that mice rapidly learn sparse internal representations of their surroundings and suggest that animals use structured behavioral sequences to reduce the complexity of natural navigation.

## Introduction

In nature, animals navigate complex terrain where sensory landmarks do not clearly reveal their location (*1*), and they rapidly adapt to changes in their surroundings (*2*). While lab experiments in simple environments have shown that rodents can memorize mazes (*3–5*) and exploit short-cuts (*6*), little is known about navigation in naturalistic terrain (*1*, *7–9*) that lacks globally visible landmarks, clearly defined routes, or discrete decision points (*10*). Further, because natural landscapes are large (*11*, *12*) and self-similar (*13*, *14*), local sensory information is not sufficient to localize oneself in space. Thus navigation requires integration of self-motion (*15*) with ambiguous sensory information (*16*, *17*). Beyond this strategy, humans handle such complexity by learning hierarchical representations of space (*18*, *19*)—sacrificing accuracy in favor of a connectivity structure that can be supplemented with local sensory information (*20*, *21*)—and by re-using memorized paths (*22*, *23*). This ability to simplify complex tasks by bundling sequences of sensory cues and motor actions into mental objects is thought to be a fundamental building block of higher cognition (*24–26*). In nature, many species navigate via stable routes (*8*, *9*, *27–29*), but it is unknown to what extent these routes arise in response to local environmental features or are generated from internal memorized sequences.

We thus sought to test whether and how mice learn to organize their action repertoire into structured extended sequences, by studying how they navigate complex naturalistic scenarios. We quantified how they spontaneously navigated without reinforced goals, in large self-similar artificial landscapes in darkness, and in a 9×9 meter grassy outdoor enclosure (Fig.S1A,S2). Indoors, we recorded wild-derived mice (N=67, PWK * C57BL/6J F1 hybrids, aged 3-10 months, see tables 1–3) continuously in the dark for multiple 8-hour sessions on consecutive days (Fig. 1A). Outdoors, we recorded wild-derived mice continuously for 3-4 days with thermal cameras (N=6, Fig. 2A, see Methods), enabling us to identify behavioral strategies that generalized to more naturalistic environments. To create complex and self-similar terrain indoors, we constructed two landscapes of 1.2 or 1.8 m diameter from 52 or approximately 151 hexagonal tiles of 17 cm diameter (*30*). Tile heights varied in steps of 2.5 cm, yielding a variety of shapes, from small steps that can be easily traversed, to 40 cm tall towers that cannot be easily jumped (Fig.1A, Fig.S1B&C). Landscapes were illuminated with infrared LEDs and otherwise dark, limiting sensory information to local tactile and auditory cues about the shape of the landscape, and olfactory cues left by each mouse. The combination of darkness, repeated local shapes, and a lack of global orienting cues made local sensory information ambiguous in many locations (Fig. 1A,C, Fig.S1E), posing a complex navigation challenge.

**Fig. 1.**
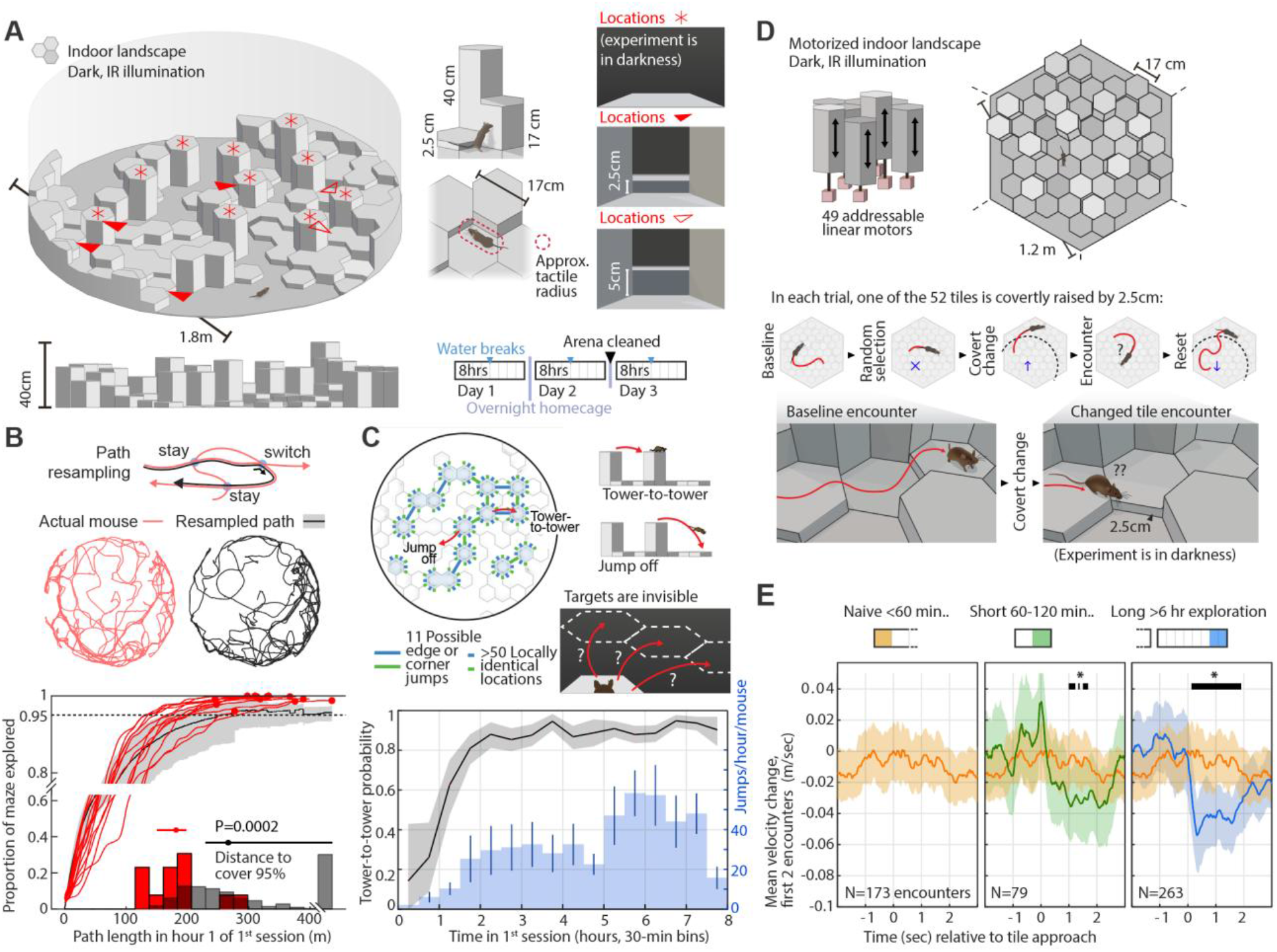
Mice quickly learn complex landscapes. (**A**) Illustration of the self-similar artificial indoor landscape. Many locations share the same local shape. Mice were filmed in darkness by an overhead camera, and their nose and tail base were tracked using SLEAP (*37*). After 4 hours, water was made available (see Methods). Mice were housed in their home cages between sessions. (**B**) Mice covered the entire arena within the first hour (distance needed to cover 95% of arena, N=13 mice, red: real vs. black: physically possible resampled paths, P= 0.0002). (**C**) Points from which mice jumped are not easily identifiable via local cues; e.g., the tops of hexagonal towers have 6 identical sides. Mice reliably jumped between towers from memory in the dark after ∼2 hours. (**D**) In a motorized variant of the landscape, we exposed mice to small covert changes of individual tile heights, again in darkness. (**E**) Mice were able to detect these changes after 2 hrs of exploration (Plots show 95% CI of mean slowdown relative to previous encounters of same tiles, bars show P <0.05 against <60 min condition, N=35 mice with 0-120min, and N=10 with >6 hrs of exploration).

**Fig. 2.**
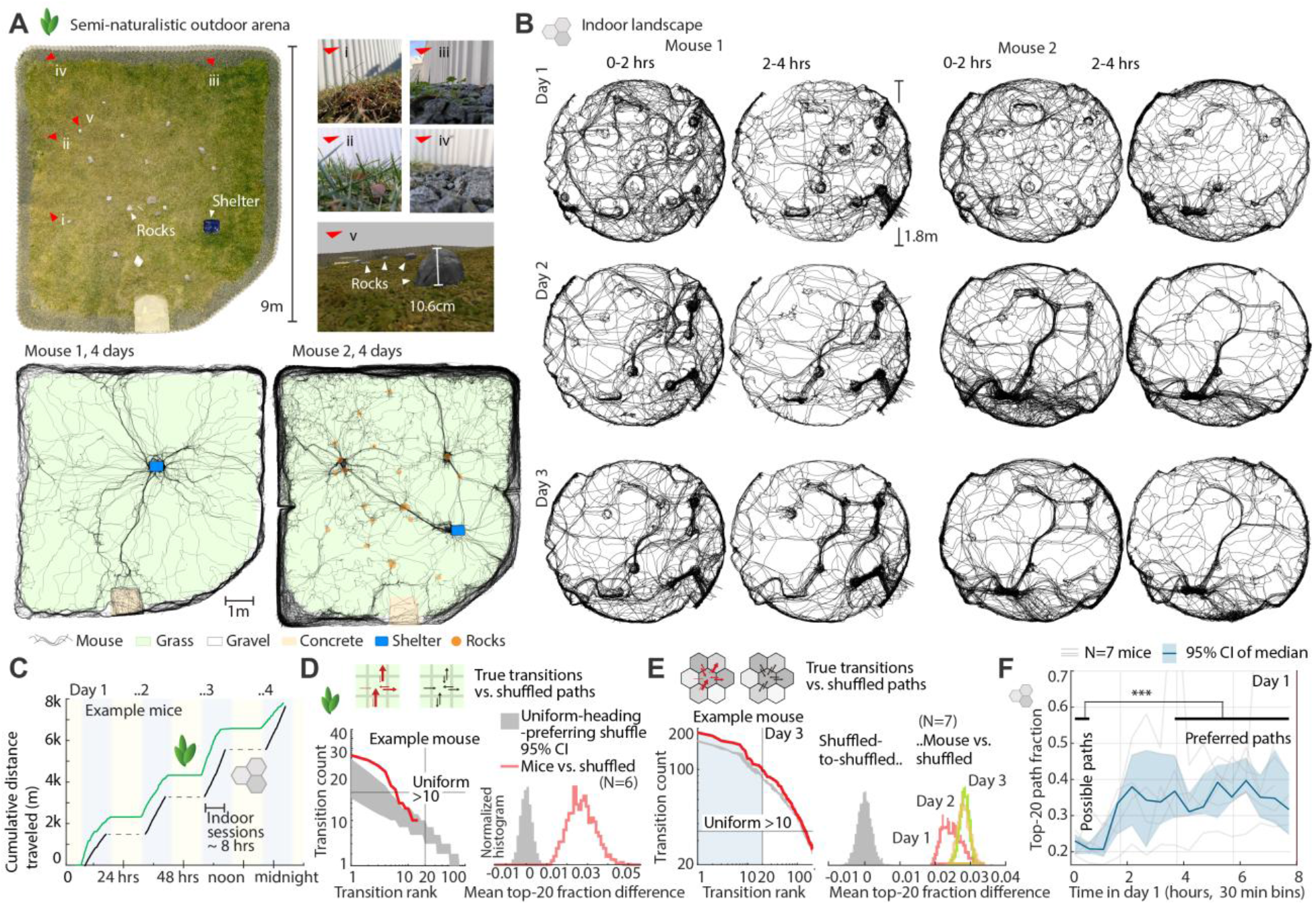
Mice converge on a sparse subset of preferred paths. (**A**) Large outdoor enclosure with a shelter box, short grass, and rocks (N=6 mice). Right: example paths of two mice showing that many routes can be taken, but few are taken repeatedly. (**B**) Example trajectories from two mice in the indoor landscape showing initial uniform coverage (see Fig. 1B) and subsequent convergence on sparse paths. (**C**) Distance run by one example mouse outdoors over 4 nights (green), and by another example mouse in the large indoor landscape over 4 sessions (black), aligned to nights for visual comparison. (**D**) Left: Frequency of specific grid transitions on one example day outside. Grey: 95% CI from shuffling paths with uniform direction preference (see Methods). The horizontal line shows the count at which all transitions that are taken >10 times are equally likely. Right: Summary statistics (P < 0.0002 vs. shuffle, N=6 mice). (**E**) Same as D but for indoor landscape. Right: Summary stats (P<0.0005 vs. shuffle, N=7 mice). (**F**) Sparsity of paths quantified as proportion of the 20 most frequent tile transitions of all transitions, showing sparsification over time (P<0.001, first 30 min vs. hrs 4-8, N=7 mice).

**Table 1.** Large indoor landscape animals (N=21):

| Mouse | Sex | Age (months) | Large indoor landscape experiment | N days |
| --- | --- | --- | --- | --- |
| 3 (not reverse-cycled) | F | 7 | (Config A) + local edit on day 4 | 4 |
| 4 (not reverse-cycled) | F | 7 | “ | 4 |
| 5 | M | 6 | “ | 4 |
| 6 | M | 6 | “ + global rotation control | 4 |
| 7 | M | 6 | Hysteresis (Config B, then A) | 5 |
| 8 (not reverse-cycled) | M | 7 | “ | 5 |
| 9 | F | 4 | “ | 5 |
| 10 | F | 4 | “ but (B, then A, then slightly modified A, then whisker trimming controls) | 22 |
| 11 | F | 5 | Hysteresis (Config B, then A) | 10 |
| 12 | M | 6 | Initial exploration only (Config A) | 1 |
| 13 | M | 6 | “ | 1 |
| 14 | M | 6 | “ | 1 |
| 15 | M | 6 | “ | 1 |
| 16 | F | 5 | “ | 3 |
| 17 | M | 4 | “ | 1 |
| 18 | M | 4 | “ | 3 |
| 19 | M | 7 | “ | 3 |
| 20 | F | 5 | “ | 3 |
| 22 | F | 8 | Split landscape, then full (Config A) | 4 |
| 23 | F | 7 | “ | 4 |
| 25 | F | 9 | “ | 4 |

**Table 2.** Reconfigurable landscape animals: (N=46)

| Mouse | Sex | Age (months) | Reconfigurable indoor landscape experiment | N days |
| --- | --- | --- | --- | --- |
| Piccolo | F | 4 | Global rotation of shape + covert changes after >3 days | 4 |
| Ktan | F | 4 | " | 5 |
| Malo | F | 5 | " | 5 |
| Klein | M | 5 | Covert changes after >6 hours | 1 |
| Petit | F | 5 | " | 2 |
| Pequena | F | 5 | " | 1 |
| Xiao | M | 4 | " | 3 |
| Chaparro | M | 5 | " | 2 |
| Malenkiy | M | 7 | " | 2 |
| Chiisai | F | 5 | " | 5 |
| Little | F | 7 | Covert changes start immediately, ~1-2 hr sessions | 1 |
| Tiny | F | 7 | " | 1 |
| Bb | F | 7 | " | 1 |
| Smol | F | 7 | " | 1 |
| Lütt | M | 9 | " | 1 |
| Kurz | M | 9 | " | 1 |
| Minute | F | 8 | " | 1 |
| Compact | F | 8 | " | 1 |
| Not big | F | 8 | " | 1 |
| Pouco | M | 8 | " | 1 |
| Kuchak | M | 8 | " | 1 |
| Liten | M | 8 | " | 1 |
| Maliit | M | 8 | " | 1 |
| Gamay | M | 8 | " | 1 |
| Kecil | M | 8 | " | 1 |
| Cilik | M | 8 | " | 1 |
| Sedikit | M | 8 | " | 1 |
| Iti | M | 8 | " | 1 |
| Saeutik | M | 8 | " | 1 |
| Femto | F | 9 | " | 1 |
| Micro | F | 9 | " | 1 |
| Nano | F | 9 | " | 1 |
| Munchkin | F | 8 | " | 1 |
| Funsize | F | 8 | " | 1 |
| Maz21a | F | 10 | Covert changes start immediately, ~2 hr sessions | 1 |
| Maz21b | F | 10 | " | 1 |
| Mini40 | M | 5 | " | 1 |
| Mini41 | M | 4 | " | 1 |
| Mini42 | M | 4 | " | 1 |
| Mini43 | M | 4 | " | 1 |
| Mini44 | F | 9 | " | 1 |
| Mini45 | F | 9 | " | 1 |
| Mini46 | F | 9 | " | 1 |
| Mini47 | M | 9 | " | 1 |
| Mini48 | F | 10 | " | 1 |
| Mini49 | F | 10 | " | 1 |

**Table 3.**
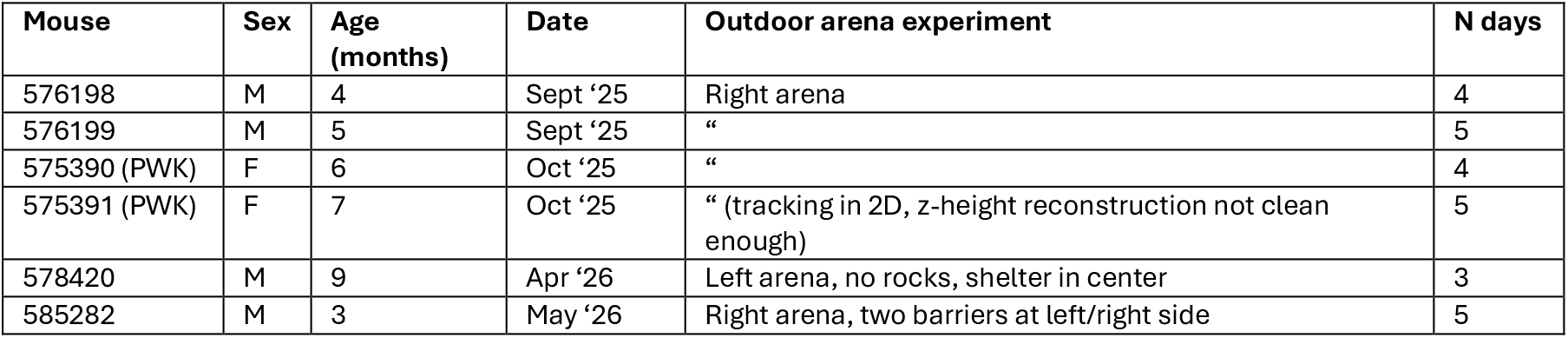
Outdoor arena animals: (N=6)

### Mice survey new complex environments with structured exploratory behavior

Naïve mice quickly explored the entire large complex landscape in darkness (Fig. 1B). Mice covered all tiles of the maze at least once in the first hour (after traveling [150 198] m real vs. [222 436] m for re-sampled paths, distance to 95% coverage in 7.2 cm grid, 95% CI via bootstrap, Fig.1B, Fig.S1H&I, P= 0.00024 signed-rank over mice), showing that they avoid repeatedly exploring the same regions, similar to behavior in binary mazes (*4*). Alongside this structured exploratory behavior, mice initially repeated a pattern of slow outwards exploration and faster backtracking along the same path (Fig.S1F&G, Video1, first hour: 0.175 vs 0.200 m/sec excluding data <0.1m/sec, P = 0.0002 paired, N=13 mice), as anecdotally described in water shrews (*31*). When backtracking, mice typically did not use short-cuts (Fig. S1F), suggesting the use of a memory-based path reversal or of olfactory or thermal cues. These exploration patterns that rapidly covered the full landscape differ from behavior in illuminated featureless boxes where rodents pick home locations (*32*), stick close to walls (*33*), and increasingly explore the landscape after habituation, modulated by anxiety (*34–36*).

After ∼20 min, mice also transitioned from preferring recessed areas near walls or surrounded by high towers (*34*) to spending more time on top of high places (Fig. S3A-C). Outdoors, mice started to jump on rocks (∼10 cm height) after ∼2-4 hours, indicating a similar preference for elevated locations (Fig. S3D-G, Video 2). These behaviors suggest that mice are intrinsically driven, in the absence of explicit reinforcement or threats, to systematically and fully explore new complex environments.

### Mice rapidly learn to jump across gaps and to detect covert landscape changes

We next tested the precision of the spatial knowledge that mice acquired about the indoor landscape. Within the first 2 hours of exploration in darkness, mice began to jump between nearby towers (Fig. 1C top) that were separated by 14.5 cm (edges) or 9 cm (corners) and thus precluded tactile perception via whiskers (*38*, *39*). The probability of a jump between two towers increased from 23% in the first hour to 88% in hours 2-4 (Fig. 1C bottom, P<0.001 via bootstrap). These jumps present a challenging localization problem: of the >50 locally identical jumping-off points, only 11 permit jumps to another tower (separated by one tile gap), suggesting that mice rapidly learned to discern specific routes to these jumping-off points.

Mice also concatenated jumps without slowing down in-between (Fig.S4 A,B), and they continued jumping after vibrissae trimming (Fig. S4 I,J). This suggests that they jumped in darkness without first verifying the presence of the target landing spot with vision or touch. To further test this hypothesis, we covertly removed one of the towers before day 4 of the experiment (Fig. S4C,D). As a result, mice jumped into air and fell (Fig. S4D,E, Video 3). Mice also jumped across gaps without having recently visited landing spots (after 1 hour, 20% of tower-to-tower jumps occur after >30m paths from the landing spot, Fig. S5D-G), and from locations that lack distinct landscape shapes (for straight paths to towers, 53% of paths are ambiguous after 2 tile-steps, 28% after 3 tile-steps, Fig. S5A-C), suggesting that jumps across specific gaps are driven by a more extended global form of memory and not by local path integration or learned local sensory features alone.

To further test the accuracy of landscape learning, and to control for possible olfactory or other allocentric cues, we let mice freely explore a smaller motorized landscape of 52 tiles (Video 4). We then covertly changed the height of individual randomly-selected tiles (Fig.1D). Whereas naïve mice that had explored the arena for <60 minutes did not react by slowing down and exploring the manipulated tile upon encountering it (Fig. 1E, N=35 mice, 173 encounters, CI= [-0.023, +0.010] m/sec speed change at 1 second relative to previous encounters of the same tiles), more experienced mice did; after only 2 hours of exploration in darkness, mice slowed down at manipulated tiles (N=35 mice, 79 encounters with 2 hrs, N=10 mice, 263 encounters >6 hrs, P< 0.004, compared to naïve mice, Video 5, Fig.S6). These results show that mice, whose only prior experience was in a 15 x 30 cm home cage, quickly acquired and maintained precise spatial knowledge of complex landscapes that was specific down to individual jump locations and tile heights.

### Mice navigate complex environments using a sparse vocabulary of paths

After initial exploration both indoors and outdoors, the paths mice took became increasingly sparse (Fig. 2A,B,D-F, Fig. S7, S8). This sparse ‘vocabulary’ of repeated paths contrasted the more uniformly distributed paths taken early in exploration (as in Figs. 1B, 2B, S7). Mice chose a sparse subset of paths over similarly feasible alternative paths (Fig.2E, Video 6, P<0.0001 comparing distribution of directed tile transitions vs. paths shuffled with a preference for uniform heading distribution, see Methods, Fig.S8), and paths became sparser over time (Fig.2F, P<0.001 comparing first 30 min vs. end of day 1). This held true when excluding jumps (Fig.S8H). Mice also used sparse paths in the outdoor arena (Fig. 2A,D, Fig.S8A-D, 5/6 mice P=0.005, one mouse P=0.095, same method), consistent with previous reports of habitual paths in the wild (*8*, *9*, *27–29*) and thus not specific to our indoor landscape. We reasoned that in complex environments, sparse path vocabularies arise through learning and could support efficient navigation.

### Mice navigate along learned directional paths

We next tested to what extent the sparse path vocabulary (Fig. 2) results from mice learning specific path sequences in the landscape, or whether the paths could be explained by generic preferences for specific types of locomotion (e.g. jumping) or egocentric landscape features (e.g. lower grass, valleys, walls, etc.). Towards this, we first asked how the paths taken by different mice navigating the same landscape compared to each other (as in Fig. 2B). To compare the structure of mouse paths, we analyzed their structure independently of running speed or motor variations: we interpolated paths to a constant speed (4 and 10 cm/sample, indoors and outdoors) and then identified similar path segments: For every point on a path, we defined a ‘path bundle’ as the times and lengths of earlier or later segments that approached it within 5 cm (indoors) or 50 cm (outdoors). Comparing shared bundle segments then measures the similarity of sets of paths, e.g. across mice or across time segments (Fig. S9, Methods). Individual mice in the same landscape converged on overlapping path patterns but also exhibited individual variability and variability over time (Fig. 2B, Fig. S7, S10). This leaves open to what degree the path vocabularies are driven by local landscape features alone or learned extended sequences fit to the environment structure. Additionally, mouse path structure was robust to rotating the shape of the motorized landscape while maintaining all other allocentric cues (Fig. S12), showing that path decisions were not driven by room cues in our experiment.

If the path vocabulary were solely a function of the landscape itself, we first reasoned that it should stabilize as mice learn the constant landscape. We tested this hypothesis by comparing the path vocabulary across all 30-minute bins using the path bundle method. We found an autocorrelation in the structure of the paths (Fig. 3A), where paths taken close in time were more similar to one another. Further, paths across the break between days 2 and 3, where we cleaned the maze, were more similar to each other than to paths within the same day (D2/D3 across-day lags 2-4 h vs. D2CD3 within-day lags 6-8 h, P = 0.0156, signed rank, N=7 mice, Fig. 3B). Thus, path vocabulary changed over experience rather than stabilizing, and this path memory was maintained across overnight home-cage rest periods, and was not primarily driven by olfactory cues (*40–42*). Outdoor paths showed a similar pattern, with autocorrelations on the order of ∼400 meters (Fig. 3ECF, 300m vs. 500m bin P=0.031 paired sign rank, N=6 mice).

**Fig. 3.**
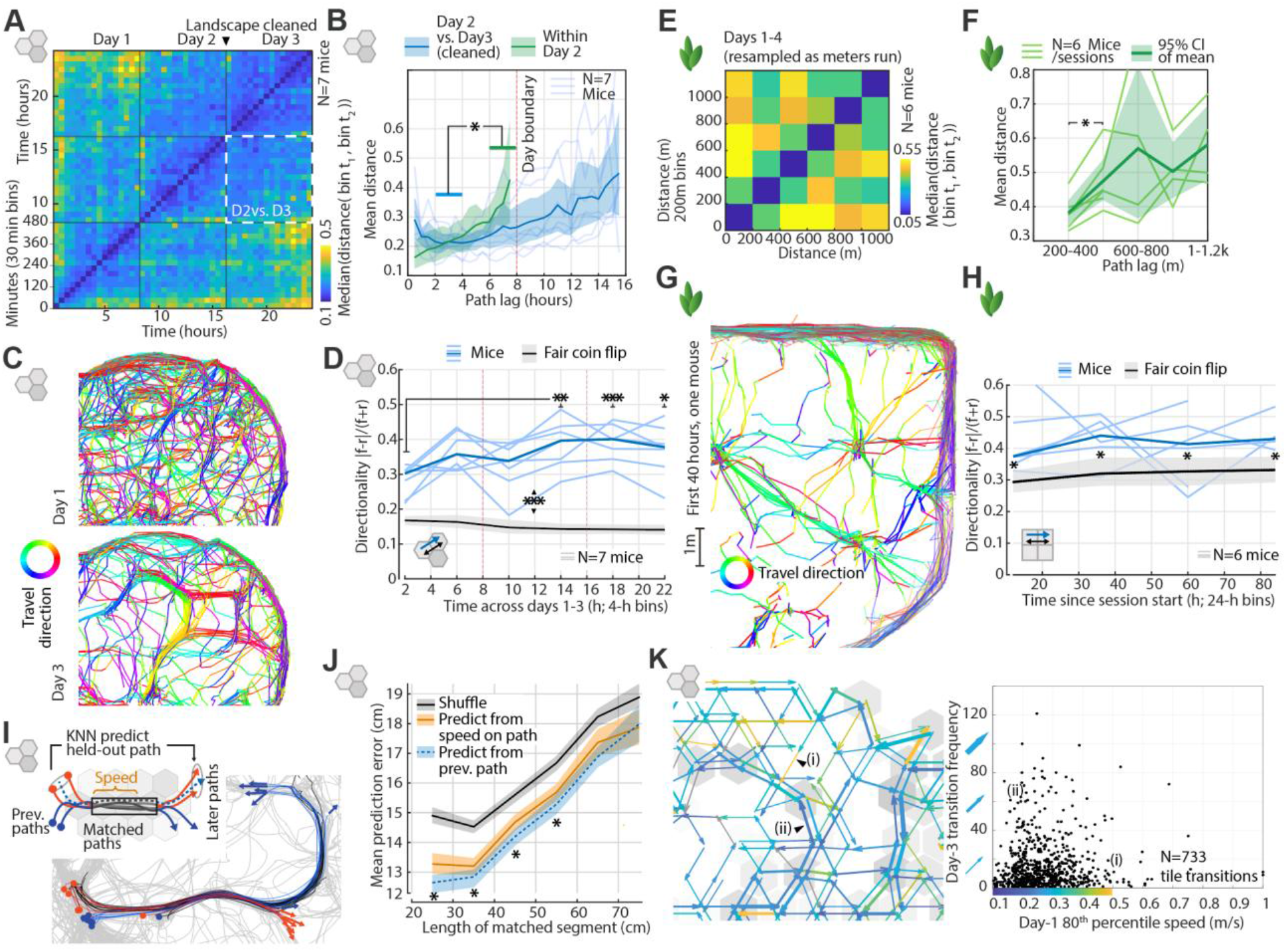
Sparse paths are learned and change over time. (**A**) Distance between path bundles, (see Methods, Fig. S9) between all 30-minute bins in the indoor landscape, summarized across 7 mice. (**B**) Path structure autocorrelation was retained across the Day 2-Day 3 gap where we cleaned the maze. (**C**) Tile transitions are more directed than fair coin flips and become more directed over time. First 2 hours of days 1 and 3 of one mouse. (**D**) Tile transition directionality over time and against fair coin flips (pooled directionality: 0.34 mice vs. [0.0786 0.0852] fair coin 95% CI, P<0.0001, N: 289 tile pairs with >10 traversals). (**E**) Same as A but for outdoor enclosure (exploration of periphery is excluded, bins are defined over distance due to gaps in exploration). (**F**) Same as B for outdoor data (300m vs. 500m bin P=0.031 paired sign rank, N=6 mice). (**G**) Same as C but for outdoor data. Note that directions at arena walls are mixed but sparse paths are primarily unidirectional. (**H**) Same as D for outdoor data, stars indicate p <0.05 against fair coin flips. (**I**) The ends of paths are predictable from their beginnings across shared intermediate segments. (**J**) summary statistics (N=1543 path sets with >6 paths, stars indicate P<0.005 via bootstrap). (**K**) Path preference on day 3 is not explained by how easy the paths are to navigate, quantified via 80^th^ percentile speed on day 1. Left: day 3 transition counts by arrow width, colored by day 1 speed. Right: Tile transition frequency versus speed for one example mouse: N=733 unique directed tile transitions, Spearman’s r=0.05 p=0.215 (Fig.S8L,M).

Second, if the path vocabulary were explained by preference for local landscape features (e.g. next to a tower, shorter grass), then these preferences should apply to the same locations regardless of travel direction. We found that indoor paths became direction-specific over time (Fig. 3C,D), and outdoor paths were also directional (Fig.3G,H) showing that both indoors and outdoors, mice learned directional paths, and local landscape features alone did not determine the path vocabulary. The preferred set of paths was not biased toward those that were easy to traverse (Fig.3K, Fig. S8L&M, speed on day 1 does not predict transition frequency on day 3, spearman r range [0.008, 0.136] across mice, 2/7 mice individually p<0.05). Thus, the sparse path vocabulary did not only reflect the landscape itself and local sensorimotor features within it; rather, pathing decisions are driven by a more extended evolving internal decision process.

To test if location-specific navigation paths were guided by more extended sequence memory in addition to sensory cues, we identified times when mice repeated paths but diverged before and after a matched set of intermediate segments (for 20-60cm, N= 1543 sets of paths with >6 matched segments, Fig. 3I). The paths after the matched segments could be predicted by the paths before the matched segments, and this prediction was better than a prediction from running speed during the matched segment (N=34416 predictions in 1543 matched path sets in 7 mice, days 2&3, median errors from prev. path: 13.77 cm, from speed: 14.24 cm, P<0.0001 signed-rank, P=0.0312 across per-mouse medians, Fig.S11). This suggests that during matched path segments with similar sensory cues, mice maintain a memory state that informs future path choices, possibly analogous to splitter cells in trained short-term memory tasks (*43–45*).

Together, these results show that the sparse path vocabulary is not solely a function of exposure to this type of artificial landscape, nor preferences for local landscape features, but rather reflects extended internal knowledge of sequences that are fit to the landscape and change with experience.

### The learned paths allow for a compressed map and make sensory data more predictable

The sparseness of the path vocabulary, and its dependence on an internal memory, suggests that it could simplify the computational complexity of navigation, similar to strategies in humans (*18–21*). We first tested whether the sparse paths make it possible to navigate using maps that omit infrequently visited areas. We simulated agents that use a map to maximize location accuracy while compensating for different degrees of path-integration error (Fig. 4A, Bayesian ideal observer, see methods and supplementary materials). Each map was composed of sensory cues that encoded specific tile height transitions; by iteratively removing the least informative cues via greedy search, we generated optimal maps of different sizes. Optimized maps yielded lower localization errors than random maps, and maps optimized for mouse paths were more compact than those optimized for random paths or rotated mouse paths (Fig. 4B-D, Fig. S13). This suggests that mouse paths are matched to the landscape in a way that could allow mice to navigate effectively with partial knowledge of the landscape.

**Fig. 4.**
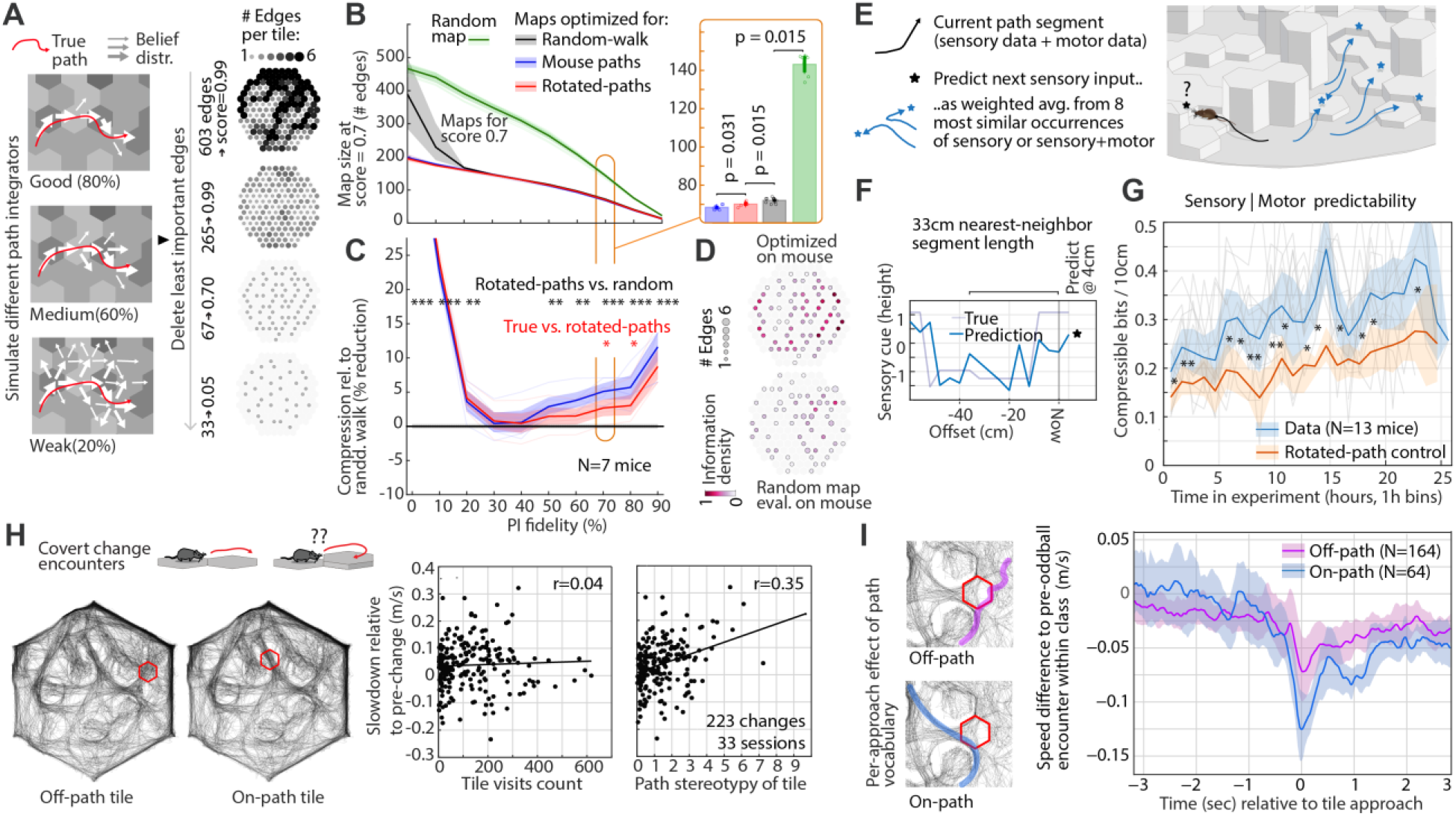
Paths allow for a compressed map and make sensory data more predictable. (**A**) Based on the mouse path, we simulated accurate or inaccurate path-integrators that diverge from the true path and re-localize using sensory cues and a map. For each path-integration (PI) level, we generated optimized sparse maps of varying sizes. (**B**) Maps for mouse trajectories were more compressible than for random, or rotated trajectories. Right: statistics for a 70% integrator (N=7 mice). (**C**) Mouse paths allow for smaller maps for strong path integrators. Stars indicate significance levels. (**D**) Two example maps from a 70% integrator, mouse-optimized maps are more informative (see Methods). (**E**) Computing sensory compressibility via forecasting: cues are predicted by averaging 8 nearest neighbor (‘NNs’) sensory & motor segments in 1-h time bins. (**F**) Example prediction. (**G**) Over time, sensory cues become more predictable from motor actions (P=12/13 mice with rising slope, P=0.002). (**H**) Per-tile slowdown on changed tiles (Fig.1D&E) is correlated with whether the tile lies on a preferred path (mean number of repeated path segments through the tile before the change vs. visits to tile). (**I**) For individual changed tiles, approaches on preferred paths (fraction of all earlier crossings of that tile that entered from the same neighboring tile) correlates with stronger slow-down (speed rel. to earlier crossings from the same entry tile, Off-path: commonness <0.2, N = 164; on-path: >0.4, N = 64).

Beyond decreasing the memory demands for encoding the landscape, the sparse path vocabulary may act as an implicit model of the landscape, supporting efficient knowledge building by increasing sensory predictability. Internal models that can predict sensory inputs (*46–50*) make those inputs compressible by allowing an agent to preferentially encode inputs that are unexpected, or by expressing complex sensory data as parameters of a simpler model. We reasoned that if specific path sequences become sufficiently uniquely associated to individual locations in the landscape, then sensory cues along these paths could become predictable from the motor sequences that form the paths. To test this hypothesis, we predicted upcoming sensory cues for each 4 cm step along a mouse path by first finding the 8 nearest-neighbor path segments (‘NNs’, ∼33cm length, chosen within a 1-hour time bin, Fig. 4E) with the closest matching sensory profile (height of tiles at the mouse head and sides). We then predicted the next sensory input as a weighted average of these NNs (Fig. 4F). When we added motor data (speed & turn speed sequences) to the NN path lookup, the sensory prediction improved. We quantified this improvement in bits, capturing the extent to which knowledge of the path’s motor sequence allows compression of sensory data (see Methods). Compressibility increased over the first 3 days of exploration (Fig. 4G, P=0.002, slope P<0.05 in 12/13 mice), showing that mice could predict upcoming landscape features from their sequence of actions. Compressibility was reduced when we rotated the mouse paths relative to the landscape (real vs. rotated paths P<0.0001 per-bin signrank, 14/23 time bins significant, N=13 mice, rotations >90 deg), showing that this compressibility results from a matching of the path vocabulary to the landscape, and not from movement patterns alone.

To test whether sparse paths play a role in how mice learn complex landscapes, we returned to the slowing-down responses exhibited by mice upon encountering covert tile height changes (Fig.1 D&E). Mice reacted to changed tiles that were on stereotyped paths more than to changes of frequently visited tiles (Fig. 4H; stereotypy effect > visit-count effect, P = 0.008 with baseline crossing speed as a covariate; P < 0.001 with slowdown expressed as a fraction of baseline speed), and they exhibited more reliable reactions when visiting the same tiles from a common path (Fig.4I, deceleration from the second before contact to 0.2 to 0.3 s after it: on-path exceeds off-path by 0.048 m/s, CI = [0.011, 0.082], session-level bootstrap P = 0.008). This suggests that the model of the landscape learned by a mouse may be related to its path vocabulary. Together, these findings point to the sparse path vocabulary as a potential strategy for increasing sensory predictability and reducing memory load to enable navigation in complex environments.

## Discussion

Here, we observed that in the absence of an explicit task, mice quickly learned to navigate complex, large, and locally ambiguous environments including self-similar artificial landscapes (Fig. 1) and a semi-naturalistic grassy outdoor enclosure (Fig. 2). This entailed a brief period of structured exploration, followed by an extended period during which mice learned and refined a sparse set of extended paths that were driven by internally generated choices and matched to the environment. Within a few hours, mice were able to jump across specific memorized gaps in the dark (Fig.1C) and detect small changes in complex landscapes (Fig.1D,E, Fig.4H,I), demonstrating the ability to quickly acquire precise spatial knowledge and to resolve sensory ambiguities within the landscapes. This study shows how self-structured naturalistic behavior, including in nature, can be used to study cognition (*51*), by applying analyses that do not require trial repetitions, and by using targeted lab experiments to test hypotheses that arise from more naturalistic data.

We find that the path sequences the mice take are learned rather than resulting from preferences to specific local landscape features (*8*, *9*, *27–29*, *33*, *52*), suggesting that mice maintain spatial knowledge in a compressed yet adaptable path vocabulary that reflects the structure of the environment and enables effective navigation. Paths were matched to the landscape such that they made sensory cues more predictable from sequences of motor actions (Fig. 4E-G), and under simplified assumptions, enabled effective navigation with partial landscape knowledge (Fig.4A-D). In this strategy, a subset of locations in the environment could be used as landmarks, between which mice navigate by relying on path integration while comparing sensory cues to an internal model (*53*, *16*).

Learned task-specific representations have been observed across a wide array of tasks in humans and other mammals (*22*, *54*). In navigation, recent work has shown how discrete paths allow for more efficient planning, faster inference, and rapid recalibration of movement around obstacles (*55*); in this view, the path vocabulary could be supported by a compact set of ‘anchor points’ that are adapted over time and used to compose extended paths. Re-casting planning in terms of learned extended sequences (*56*, *55*, *57*) could provide a natural substrate for successor representations (*49*, *50*), and could shed light on how ‘cognitive maps’ can arise from solving planning problems (*58*, *59*).

The brain encodes repeated sub-routes in mazes and progress towards navigational goals along paths in hippocampus (*43–45*) and cortex (*60*, *61*). Our behavioral results suggest that location in environments, even in the absence of explicit task structure, could be represented in terms of path identity and sensorimotor sequences, in addition to allocentric map coordinates, similar to how humans reason about space in terms of warped hierarchical maps (*19–21*).

Our results point towards applying a Bayesian sensorimotor learning perspective to navigation (*62–64*, *55*): instead of an explicit map, mice can learn a sparse set of environment-specific sequences; these sequences could enable mice to maintain and update hypotheses about their environment through the comparison between sensory cues and path-specific expectations, and to rely on path integration otherwise. In this view, mice interact with the world not by directly reacting to sensory inputs, but rather by building a sparse internal model of the world. Building on decades of research on the neural basis of navigation (*65–67*), these results pave the way for future work to address the neural mechanisms behind fast, structured learning of world models in naturalistic behavior.

## Supporting information

Video 1. Initial exploration backtracking behavior

Video 2. Mice outdoors climb on rock

Video 3. Jumps

Video 4. Exploration behavior in reconfigurable landscape

Video 5. Detection of covert changes

Video 6. Exploration in large indoor landscape

Sparsemap documentation

## Acknowledgements

We thank Roian Egnor, Sandro Romani, Laura Lewis, and Mark Nicholas for discussions and comments on the manuscript. Kristin Branson, Brett Mensch, Christopher Rodgers, the class and TAs of 2015 of the Transylvanian Experimental Neuroscience Summer School organized by Raul Muresan, Adam Kampff, and Florin Albeanu, as well as Juan Ignacio Sanguinetti, Federico Carnevale, Petr Znamenskiy, Goncalo Lopes for discussions. Jon Newman for discussions and work on earlier versions of the indoor landscape. Angela Jacobs and Josue Perdomo for mouse handling and experimental support. Peter Polidoro, Sam Jager, Alex Sohn, Jeff Talbot, and Jon Arnold for design and fabrication of the motorized landscape, Joanne Sheppard for mouse handling support and work on experiment control software. Roian Egnor, Jessica Teran, Gillian Harris, Daniel Savson, Jim Cox, Bill Garth, Ray Cho, the Janelia IACUC and facilities teams for work on the outdoor arena. This research was funded by the Howard Hughes Medical Institute at the Janelia Research Campus as part of the Mechanistic Cognitive Neuroscience program.

## Contributions

Conceptualization: JV, AH, TM, AC Lab data acquisition: JV, VP, AC Covert change task data acquisition: AC Outdoor data acquisition: JV, EJD Outdoor data processing and methods repository: JV, EJD, LK, AC Data curation: JV, TM, VP, AC Data analysis: JV, AH, AC, TM Simulation & modeling: TM, AH Supplementary text: TM, AH Pathbundle & sparsemap repositories: TM Writing: JV with input from all authors.

## Data and code availability

Code used in this paper is available on github, at https://github.com/HermundstadLab/sparsemap for the map compressibility, https://github.com/HermundstadLab/pathbundle for the path-bundle distance, and https://github.com/lambdaloop/mouse-outdoor-tracking for outdoor mouse tracking methods. Analysis code and datasets will be made available on Figshare with the peer reviewed paper.

## AI use

Claude (Anthropic) was used in development of parts of the code, all code was manually reviewed.

## Methods

### Animals

73 mice, either F1 crosses between PWK/PhJ and C57BL/6J or C57BL/6J mice (Jackson labs), or PWK/PhJ, aged 3 to 10 months, males and females were used in this study. N of animals per experiment was: Large indoor landscape N=21 (92 sessions total), reconfigurable indoor landscape N=46 (66 sessions), outdoor arena N=6 (26 days). See Tables 1–3 for a detailed breakdown of the individual animals. Animals were group or singly housed. Group housed animals were housed in rat-sized cages (∼25 x 45 cm), single-housed mice in regular mouse cages (∼15 × 30 cm). All procedures are approved by the Janelia Institutional Animal Care and Use Committee and in compliance with the standards set forth by the Association for Assessment and Accreditation of Laboratory Animal Care.

### Vibrissa trimming

For experiments controlling for the use of vibrissa (Fig.S4I&J), we trimmed the vibrissae of a mouse to ∼2mm length under brief isoflurane anesthesia.

### Behavioral arena and data acquisition

The arena was constructed from ∼151 hexagonal prisms with flat top surfaces made from dense Styrofoam, measuring 17cm along the long axis, cut to varying heights, painted with a water-resistant acrylic primer. Tiles were cut at the periphery to fit into a circular outer wall of 180 cm diameter (Fig.1, Fig.S1B&C). Interior tiles were grouped in clusters of 7 tiles which could easily be exchanged, rearranged, and rotated to change the layout of the arena, for cleaning, or to otherwise control for olfactory cues. The landscape was reconfigured depending on the experiment, with most of the experiments conducted in the default configuration ‘A’, and some experiments in ‘B’ (See Fig.S1 for details).

Mice explored the arena in their dark period (some exceptions are indicated in Table 1), for four 8-hour sessions (more in some experiments, see Table 1) on consecutive days, interrupted after 4 hours for a brief period in which water was made available in a small dish for at least 10 minutes. Mice were housed in their home cage between sessions. If indicated by the experiment (see Fig.1), or between animals, the arena was disassembled and the tiles were cleaned by first vacuuming and wiping them, spraying them with VitalOxide (Vital Solutions, Florida), wiping them clean, and then fully submerging them in water, with a final rinse in the same water bath for the entire maze, distributing any possible local olfactory cues.

Behavior was tracked with an overhead camera either at 1195 x 1000, or 2448×2048 pixels and 25 or 40Hz in Bonsai (*68*). All experiments were conducted under infrared light (850/940 nm) with no visible light (light levels at ∼10^−4^ lux) depriving mice of visual cues (*cS*, *70*). Animal position was tracked using SLEAP (*37*) using the nose, ears, base of head, and base of tail as keypoints.

### Reconfigurable arena

For a subset of control experiments, we used a smaller motorized arena made from 55 full and 18 half hexagonal prisms of the same individual dimensions as in the main arena, but made from ABS plastic, with smooth sides and a textured upper surface. The outside width of this arena was 120 cm. The height of 49 of the tiles could be changed via linear actuators, making it possible to create arbitrary patterns. Experiments in this arena were conducted with a white noise background of 73 db(C) (C weighting, extech sl510 meter) in total darkness (light levels at ∼10^−5^ lux, impossible for a fully dark-adapted human to identify shapes in the arena). The arena was cleaned in between animals and as described in the results section by vacuuming and wiping the top and side surfaces of the prisms with VitalOxide as for the main arena.

### Novelty detection experiments

To test whether mice learned a sufficiently detailed representation of their environment to detect small changes, we investigated their reaction to small covert changes (spatial/shape oddballs) in the landscape in the reconfigurable arena (Fig.1D,E). Mice were either tested naively with no prior exposure to the arena, or after previous experience. The arena was completely dark and we applied a constant 73 dbc (C weighting) white noise background. The covert changes were executed by an automatic algorithm: First, either immediately or after a 6 hour delay (see Fig.S6), a 10 minute timer was started. After this timer elapsed, we first selected a random tile from a subset of 25 out of the 49 movable tiles, selected to avoid modifying the height of tall towers. The program then waited until mouse was at least 80 cm away from the chosen tile to exclude the use of auditory cues, and raised the tile by 25 mm. The program then checked that the mouse intersected the tile (13cm distance mouse center of mass to tile center) and again waited until the mouse was 80 cm far from the tile before returning it to its base position. This resulted in mice occasionally being able to re-visit the oddball tile multiple times before it was reset to the base position. The timer was then reset to 10 minutes. For analysis, we examine the speed of the mice, aligned in time to the point at which they intersect the oddball tile compared to the speed profile for the same tile in the preceding encounters. In the main analysis the first 1-2 oddball intersections (if available) are used. For Fig. 4H, one slowdown value per change was computed as the median control profile minus the mean of the first 1-3 post-change encounters (if as many are available), averaged 0.1–0.4 s after entry. Two per-tile predictors were computed from the path up to the change: visit count (number of tile encounters) and path stereotypy, the mean number of other path segments that bundled with the path for ≥20 cm at each sample within the zone, using the path-bundle method (Fig. S9). We report Pearson correlations with 95% CIs from a two-level bootstrap (sessions > encounters). Because tiles on stereotyped paths are crossed faster, we also fit the model with each tile’s baseline crossing speed as a covariate and with slowdown expressed as a fraction of that speed. For Fig. 4I we analyzed single crossings as in Fig.1. Each approach’s commonness was the fraction of all earlier crossings of that tile that entered from the same neighboring tile. Speed was expressed relative to the mean profile of earlier crossings from the same entry tile (≥2 required), and slowdown as the mean of this difference over the second before entry minus its mean 0.2–0.3 s after entry, so that differences in arrival speed between routes cancel. Off-path and on-path crossings were those with commonness <0.2 and >0.4; the group difference and its CI were obtained by bootstrapping over sessions, and the dependence on commonness was tested by regressing slowdown on commonness with time in session and prior visit count as covariates. Data were pooled across 38 sessions from 28 mice (16 sessions after >6 h of exposure, 22 in naive mice), using crossings after 30 min of exposure.

### Outdoor arena and data acquisition

Outdoor experiments were conducted in two approximately 9×9 meter arenas (‘left’ and ‘right’, see Table 3) formed by a base of gravel and soil, with grass and other plants native to northern Virginia. Plants were trimmed to approximately 15cm height with a string trimmer. Mice were provided with a home box (50 x 45 cm footprint, 30 cm height) made from plastic with an entry tunnel, and with bedding material, to provide shelter as needed. Arena walls were made from smooth corrugated steel sheets set into gravel, making tunneling out impossible, and a suspended net protected mice from birds. The floor of the box contained a scale together with a motion-triggered camera that was used to ensure that mice did not lose more than 30% of their initial weight. Animals were slowly acclimatized to outdoor temperatures before each experiment, and no experiments were conducted outside of a ±5 degree band of their prior acclimatization range or outside of 4-33 deg C. Experiments typically lasted 4 days. Occasionally, experimenters entered the arena during the day to check on mice or perform necessary maintenance. This was only done after verifying that no mouse was currently exploring the arena. Mice were filmed using 9-12 thermal cameras (FLIR triton, 640×480 pixels, 30Hz) mounted around the arena periphery using Bonsai (*68*), and mice were tracked with a simple motion detector via continuous background subtraction (*τ* = 30*sec*) and simple thresholding. Tracking data was reconstructed to a 3d trace using anipose (*71*) with custom methods for initial camera calibration and robust 3d triangulation (https://github.com/lambdaloop/mouse-outdoor-tracking).

Erroneous tracks due to insects, plants moving in the wind, experimenters etc., were removed based on position heuristics: 3D points with high reprojection error (>40 px, at camera resolution 640×480 px), points outside the arena bounds or at implausible heights, and samples with physically impossible frame-to-frame speeds (together with ±2 neighboring frames) were discarded. Isolated short track segments were additionally removed: any tracked segment whose endpoints did not pass within 1 m of a longer (≥1 min) track segment occurring within ±1 hour was discarded. Remaining gaps in tracking of less than 1 m and up to 20 s were linearly interpolated, and tracks were median-filtered (0.6 s window).

For path similarity analyses on the outdoor data, periods in which the mouse was inactive (average speed below 1 cm/s in a ∼30 min smoothing window, e.g. while in the shelter or a burrow) were excised, and tracks were additionally filtered by removing backtracking within a 2.5 cm radius: the path was advanced only once the track moved more than 2.5 cm from its last point, discarding dwelling and local back-and-forth motion, to make paths where mice slowly foraged along a trajectory more comparable. The resulting paths were resampled to equidistant 10 cm steps, and samples along the arena perimeter or within 60 cm of the home shelter were excluded.

### Statistics

Unless stated otherwise, shaded bands are 95% confidence via bootstrap resampling, either of the mean or median, as stated. For statistics across mice, 2-level bootstraps are used, sampling across mice first, then across e.g. time points. For probability data (e.g. direction of tile transitions), we assumed binomial distributions. Unless explicitly stated, non-bootstrapped P values were computed via Wilcoxon signed-rank tests for paired and Mann-Whitney U tests for non-paired observations.

### Path preprocessing

Tracking time series were preprocessed by first aligning each recording to a common reference coordinate system to eliminate any camera or landscape misalignment. This processing step also corrected for sessions where we rotated the entire landscape to test whether room-centric cues drove the mice’s path selection (Fig.S12). Paths were then filtered by excluding minor tracking artifacts in which tail position was outside the landscape perimeter or unrealistically moved >200 px across consecutive frames. This included manually reviewing and filtering out groups of frames containing the mouse cage being placed in the arena, experimenter hands, and water dish. Parts of the recording where mice were briefly invisible behind tall towers were interpolated over. Position coordinates were gaussian-smoothed.

For analyses of the path structure, equalized trajectories were computed by re-interpolating the tracks to a sampling of 4 cm (indoors) and ∼10 cm (outdoors) cm per sample and labeling spatially confined movement periods when the maximum pairwise distance among eight equalized steps was < 30 cm, yielding a smoothed, speed-equalized trajectory.

For analysis of running distance, distance was computed after significant path smoothing to exclude local search behavior and only quantify long-distance travel. In the outdoor data, x/y time series were smoothed with a running median in a 20 second window, and then cumulative distances were calculated from points every 20 seconds. For data from the indoor landscapes, x/y time series were low-pass filtered at 0.1 Hz with a 3^rd^ order butterworth filter to exclude local exploration movement and to collapse periods of stationarity into single points despite tracking jitter and mouse movement.

### Shuffled paths

For some control conditions, e.g. computing a null hypothesis for how quickly mice explore the maze initially (Fig. 1B), or to validate our pathbundle distance index (Fig.S9D), we computed shuffled substitute paths that retain all properties of the true mouse path except for its extended temporal structure.

For generating such Markov shuffled paths, from indoor-landscape data, we sample from the true mouse path, but after every 2 m, the path can jump to any adjacent point in the path within 1.5 cm if at least 4 candidate paths exist, resulting in a time-shuffled path that lacks the temporal structure of the original path but on average retains the same occupancy distribution, and is physically realistic, i.e. the resulting paths could have been chosen by the mouse. Computing a large set of these shuffled paths can then be used to compute null hypothesis confidence intervals.

For testing whether path choices are sparse (Fig. 2), particularly in outdoor data where the amount of available data is not sufficient to use the evolution of the sparsity over time relative to day 1, we computed a similar set of shuffled paths, but with an additional selection on which path to jump from that favors angularly uniform exploration: After each 1 m since the last shuffle, candidate path segments within 15 cm were selected based on their angle at which they exit a 30 cm radius by binning into 32 angular bins. The shuffled path then jumps to the path with a direction which has so far been least taken by the surrogate. To prevent local cycles, whenever the maximum bin count of this usage histogram exceeded 5, we instead jump to a random point on the recorded trajectory. Because random resampling from the set of initial paths re-samples clustered paths proportionally, this procedure could only bias the estimate towards more clustered paths, making it a stronger rather than weaker null hypothesis for tests of clustering.

### Estimating egocentric shapes around the mice

The shape around the mice was estimated by manually annotating the camera image with the height of each hexagonal prism, correcting for perspective distortions. A circular area of ∼20 cm around the base of the mouse’s head (enough to cover about one of the tiles) was rotated to align with the mouse’s heading, and the height under the mouse’s body was subtracted from this height map to derive an estimate of the egocentric area around the mouse.

For analysis of mice climbing on rocks in the outdoor arena (Fig. S3), we defined a rock approach as the mouse getting within 30cm of a manually annotated rock position. We estimated the ground height by computing the 25th percentile of the mouse’s z-height in a circle of 30-50 cm around the rock center. This method avoids having to align tracking data to CAD models of the arena and captures session-to-session variation. We then analyzed mouse behavior in 3D around the rocks.

### Quantification of early exploration behavior

Locomotion was categorized as either outward exploration or backtracking depending on whether a preceding path segment (excluding the currently ongoing traversal) was found within ∼3.8 cm of the current position. Preceding path segments were only considered within the last 8 m of path run, so that locomotion was labeled as backtracking only where the mouse re-traced a recently laid path. For analyses of inward/outward running speed (Fig. S1F,G,J), backtracking was detected on the raw tracking data with a ∼7.6 cm radius, considering preceding path segments within a 10-minute window preceding each sample (matching the 10-minute bins of the time-course analysis; the first 10 minutes of each session were excluded so that all lookback windows were complete), and only periods of locomotion (speed 0.1–2 m/s) were included.

### Jumps

Body position was low-pass filtered at 3 Hz (3rd-order Butterworth, applied bidirectionally for zero phase lag), and speed was computed as the frame-to-frame displacement of the filtered position. Jumps were defined as times when 1) this smoothed speed exceeded 0.65 m/s and, 2) path curvature stayed under 1.5 rad/sec (∼86 deg/sec) for the duration of the high-speed segment and 3) the path between the start and end of the putative jump contained a height range of ≥ 7.5 cm (3 tile height steps). Jumps were categorized as successful if they spanned a gap between two towers (min. height difference ≥ 7.5 cm, min distance one tile), and as unsuccessful if they started from a tower but ended on a lower spot. Jumps along the arena perimeter, or motion segments classified as jumps under 20 cm were excluded from this analysis.

### Quantification of loop length and path lengths

For measuring the length of the paths mice took from eventual landing spots to the locations from where they would later jump off (Fig. S5D-G), we looked backwards in time from each jump until the path approached the landing spot by 20 cm. The resulting path was then filtered with a running median (window size 6 sec) and gaussian smoothing (σ=1.1 sec) to retain broad locomotion and discard local motion. The length of this path was then analyzed. For quantifying the number of turns, we counted the number of times the mouse deviated by over 60 degrees from its previous heading along this path and every time such a deviation occurred, the current heading was set as next reference heading.

### Quantification of sensory ambiguity

The hexagonal maze structure is self-similar, with the shape of local patches repeating throughout the arena. To estimate the amount of memory required to uniquely localize oneself in this arena (Fig. S5A-C, also Fig.S1E), we used a Bayesian ideal observer method in a simplified discrete version of the arena: The arena was represented by an array of the heights of the tiles *H*, and simple paths of the mouse consisting of a series of moves in the six possible directions [[1 0][1 1][0 1][−1 0][−1 − 1][0 − 1]] between adjacent tiles were used. We then computed the egocentric sensory sequence of height steps encountered along this path 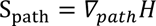 as well as equivalent sequences for translated and rotated versions of the path leading to all possible locations in the maze. By evaluating the difference between the pairs of sequences, we computed a sensory likelihood for all locations in the maze, at all headings. The error for the true location is by definition zero, since the true reference path and the location path are identical. For our analysis, we then counted how many spurious locations with no error exist for paths of increasing length. This model simplifies sensory input to only the local height steps directly under the mouse and does not account for accurate sensory information at the boundary of two or three tiles but is sufficient to estimate the degree of symmetry of the maze for typical paths taken by the mice. For illustrating the local self-similarity of the mazes (Fig.S1E), we used the same method and computed the posterior spatial hypothesis distribution that results from visiting three adjacent tiles, showing the location information that could be inferred by a mouse sensing the relative height of the intersection of these 3 tiles.

### Path bundle method and distance measure

It is challenging to measure behavioral similarity for our datasets, because they are continuous trajectories without any predefined trial structure. Given that there could be infinitely many ways to label these trajectories, it is unclear how to formulate a sensible distance measure. Ideally, we would first run a joint segmentation and clustering algorithm to convert the raw data into distributions of a finite behavioral repertoire before applying an appropriate distance measure on these distributions. Unfortunately, this approach requires extensive experimentation to select a suitable algorithm, and it is computationally costly to work with our extensive datasets (7 mice, 24 hours for indoor arena). To overcome these challenges, we first define a “path bundle” as a set of spatially overlapping paths that are repeated across different times. This allows us to quantify the structure of repeated paths, and to compare two paths, from the same mouse at different times, or across mice (See Fig. S9 for schematics and examples). To construct the path bundle, we first interpolate the path to a constant speed, so that every step is a fixed distance (see section on path preprocessing), yielding a path P of length N points. We then compute for every point xref in P the set of other earlier or later points in the path that approach xref by some threshold (unless stated otherwise: 5 cm indoors, 50 cm outdoors). For each of these points *i* and associated path, the entry *pb*^P^*_ref_*_,*i*_ in the path bundle is then assigned the length for which the overlapping path segment of P stays within some threshold distance (15 cm for lab experiments, and 150 cm for outdoor data) of *P* starting at xref (the length is counted forwards and backwards along P). The diagonal entry is set to the length of the longest other overlapping path segment. Points along the overlapping path segment that fold back over itself within 1.5*step-distance are not counted towards this length. Over all points *ref* = 0. *N* of P, this forms the complete path bundle *pb*^P^ of size N x N (e.g. Fig.S9A,C).

Path bundles span a continuum, and do not yield discrete clusters. Nevertheless, we can use the path bundle to quantify the distance between two intervals of behavior. By concatenating two paths A and B into AB so that the boundary between A and B is at point ℎ, and computing their joint path bundle *pb*^AB^, we can now compute for each point along AB whether long segments in its path bundle originate equally from both paths, or are exclusive to one of the paths. To discard short overlapping segments, elements of *pb* that are under a threshold length can be removed (2 steps for the indoor data, none for outdoor data), then compute 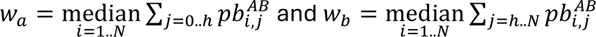 which sums up the length of paths in the bundle originating in A or B. Normalizing the difference via *d*_PB_(*A*, *B*) = |*w_a_* − *w_b_*|^2^⁄(|*w_a_*|^2^ + |*w_b_*|^2^) now gives us a distance metric between any two paths that quantifies the structure of extended path repetitions. To quantify the evolution of a path P over time, we can now use *d*_PB_(*P_si_*, *P_sj_*) for e.g. all 30-minute, or 200m segments *si*, *sj* of a path, computing a distance matrix M. Due to subsampling, this measure is not strictly symmetric, and we use (*M* + *M^T^*)/2.

Note that *d*_PB_ is fast to compute; no prior clustering is needed, because *d*_PB_ can be directly computed with unlabeled paths. Moreover, since a clustering algorithm is not required, computing *d*_PB_ only requires no further hyperparameters. However, this is not a pure measure of only the long repeated paths that we observe, and should be interpreted as a relative measure, e.g. via changes over time, or across conditions, and interpreted via statistical tests rather than by absolute numbers, as one would use measures of similarity like a KS divergence or cosine distance. Fig S9 shows examples from real and synthetic data, illustrating some key properties: In the presence of overlapping paths, the distance results in 0 for identical paths, and increases as more of the path length is occupied by aligned paths. Overlapping paths in opposing directions do not register, leaving the metric at 1. Paths that cross each other also do not register because the resulting short path bundles do not meet the threshold length. Finally, random walks can result in short but sufficient overlapping path bundles, for which the distance between two paths approximates the similarity of their spatial occupancy.

Due to the continuous nature of the path bundle, this measure is potentially prone to double counting. While this does not affect our key conclusions, we can mitigate this in future extensions of this work by formulating a revised path bundle distance between two intervals, *I_i_* and *I_j_*, as follows:

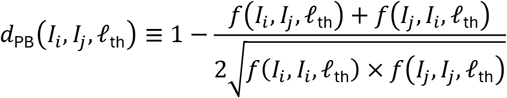

with the auxiliary function defined as

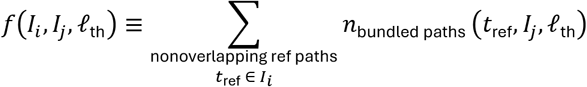

where *I_i_* and *I_j_* are two temporal intervals *i* and *j* of a continuous trajectory, *t*_ref_ is the starting time of a reference path, and *n*_bundled paths_(*t*_ref_, *I_j_*; l_th_) counts the number of overlapping paths, within interval *I_j_*, to a reference path at *t*_ref_, whose length exceeds a manually selected threshold l_th_. The nonoverlapping condition ensures that the starting time of two nearby reference paths are well separated by at least l_th_—i.e., *t*_ref_ + l ∉ {*t*_ref_} ∀0 < l < l_th_. In this study, we focus on measuring the similarity of path shape and ignore speed information. To this end, all raw trajectories are preprocessed to progress an equal distance (a quarter-tile width, ∼ 4.25cm) from one time point to the next. And l_th_ ∈ {1,2,3, …} is thereby specified in this discrete distance space. Two paths **x** = (*x*_0_, *x*_1_, …) and **x**′ = (*x*′_0_, *x*′_1_, …) overlap when their position sequences are “point to point matched”—i.e., *d*(*x_t_*, *x*′*_t_*) < *d*_th_ ∀0 ⩽ *t* < l_matched_, where the distance threshold is chosen to be a half-tile width, ∼8.5cm. It’s worth noting that this revised path bundle distance *d*_PB_ can be clearly interpreted: When all paths in intervals *I_i_* and *I_j_* can be well clustered, i.e., each path appears at most in a single bundle, *d*_PB_ simply converges to a cosine distance: 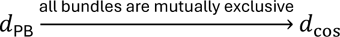. See https://github.com/HermundstadLab/pathbundle for code.

### Path prediction analysis

To test whether path decisions were driven by a memory, rather than being derived from sensory cues with some stable rule, we identified repeated path segments that were preceded and followed by diverging paths and tested whether the following paths could be predicted by the preceding ones (Fig. 3I, Fig. S11). For days 2 and 3 of mice in landscape ‘A’ (7 mice, 14 sessions), we ran path bundles from starting positions in a 20cm grid covering the inner 140cm of the landscape to exclude paths that follow the periphery. For each of these starting positions, pathbundles for all paths approaching the starting position by 20cm were computed, and the path with the bundle with most paths over 18 cm, excluding paths with stationary periods was selected for analysis. For this path, a path bundle with an initial distance threshold of 6 cm was computed, and paths were stopped when they diverged from the seed path by 10 cm, resulting in sets of paths with closely matched paths. This analysis was done using raw tracking data which captures running speed differences, in contrast to the path structure analysis for which we used speed-equalized trajectories. From this pathbundle, we then selected paths with a mean speed of >0.06 m/sec and lengths of > 4 sec, and if over 6 paths remained, this set was used for prediction analysis, yielding sets of predictions from ∼1500 bundles. For each of these bundles, we extended the paths by 0.8 sec forwards and backwards from the points where they diverged from the seed path, and discarded a margin of 0.4 sec, leaving the beginnings and ends of paths that, by construction via the pathbundle, had by now diverged from the seed path (Fig. S11 A-C for examples and schematic). For each of the resulting pre/post path pairs, we then predicted the post path from the pre path via K-nearest-neighbor prediction, using the average Euclidean distances to look up matches: For each post path, that path was removed from the set, and the 2 closest pre-paths were found, and their corresponding post-paths averaged to predict the held out path segment. As a control, the same procedure was applied to shuffled paths. To further test whether path choices could be informed by movement speed along the path, we also computed the same prediction but using the speed along the matched path up to the matching point (to avoid trivially predicting speed in the post segment from immediately preceding speed).

### Analysis of room-centric cues

To test if paths were driven by room-centric cues (AC noise, position of the door, etc.), we ran 3 mice in an experiment where we rotated the shape of the arena using the reconfigurable arena motors (Fig.S12). Mice were run for 3-4 sessions of 8 hours on consecutive days, with a 5 minute water break after 4 hrs daily. 60 degree rotations to the landscape were applied before day 3 (n=3 mice) and again before day 4 (n=2 mice). Mouse centroid position was tracked online using bonsai and an overhead camera in IR lighting. To transform mouse position paths from the room reference frame to the landscape reference frame, an affine transformation (cv2.estimateAffine2d) was fit to map the 6 maze corner points and the maze center point from each day of all mice to one reference day’s points. This transformation was applied to the tracked mouse position across all days to produce de-rotated coordinates in the landscape reference frame.

### Map compressibility analysis

To estimate whether the sparse path vocabulary allows mice to theoretically use a compressed map to navigate the arena (Fig. 4), we enumerate Bayesian ideal observers with incomplete maps of various sparsity who self-localizes using a noisy path integrator and ambiguous sensory measurements. The performance of ideal observers represents the upper limit of what information is available to mouse.

#### Self-localization task under uncertainty

While there is no explicit task for our freely moving mice, we hypothesize that mice are well equipped to self-localize after initial exploration. To test this, we model an agent as ideal observer who traverses an existing mouse trajectory with a noisy path integrator (PI) and infers its self-location from ambiguous observations against incomplete map of the arena. The agent maintains a belief distribution of self-location, which starts sharp at the ground-truth location and diffuse over time due to imperfect path integration. When an observation (relative height between two neighboring tiles) is made at a transition between tiles, the belief distribution is sharpened by ruling out impossible locations that are known to the agent. The task performance, i.e., localization score *S*(*κ*, **x,** *M*), evaluated as the mean of belief probability of ground-truth location over entire timeline, thereby depends on 1) PI fidelity *κ*, 2) the agent’s observations along its trajectory **x**, and 3) the agent’s knowledge—i.e., a sparse map *M* that encodes a subset of all 930 edges of 155 hexagonal tiles.

#### Map optimization

To test whether the observed sparse sets of paths allow the mice to efficiently self-localize, we asked how many edges (or minimal map size *m*_min_) are required to encode in a map to reach a desirable localization performance *S*_target_. Equivalently, this map optimization problem can be stated as maximizing score given a fixed map size, PI fidelity, and trajectory:

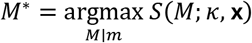

This optimization is nontrivial due to a vast solution space to search: 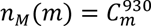 which is astronomical for even a moderate map size, e.g., *m* ≈ 50. To overcome this, we used a deletion-based greedy optimization scheme in which one starts from a full map *m*_full_ = 930 and deletes the least important edge one at a time until reaching the target map size. Note that we do not use an addition-based scheme because it is not compatible with an annealing-like scheduler that allows a major speed up in task evaluations during optimization (see Supplementary Materials for details). This greedy optimization is still costly, but much more feasible and effective than say a sampling based exhaustive search.

#### Numerical experiments

The statistical claims in this study are either to compare random maps *M*_rand_ with optimized maps *M*^∗^, or to compare mouse trajectories **x**_mouse_ with shuffle controls **x**_sh_ or random walks **x**_rw_. To make these a fair comparison, we convert mouse trajectories into tile walks in which a consecutive step does not occupy the same tile. For **x**_mouse_, we use 7 Day-3 (8 hours) trajectories from 7 mice who experience the same fixed arena for 3 days. For **x**_sh_, we use the same mouse trajectories but rotated by 60°, 120°, 180°, 240°, 300° with respect to the center of the arena which amount to 35 trajectories. These rotations break the correlation between motor behavior and sensory cues. For **x**_rw_, we generate 10 random tile walk with the same average length equivalent to 8-hour-long trajectory; each tile walk step uniformly samples the next tile from up to 6 neighboring tiles. The same map optimization is applied to find efficient maps 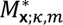 for the three types of trajectories given a fixed PI fidelity and map size. The effectiveness of the optimization algorithm is accessed by comparing optimized maps with random maps *M*_rand;_ *_m_* for which we simply shuffle the edge index *e* ∈ [0, 929] and truncated at selected map size.

#### Information density

To investigate what makes an optimized map more efficient when leveraging mouse behavior, we ask whether such a map encodes more informative edges. Specifically, we devise an information-theoretic measure, i.e., information density *ID*(*s*_encoded_), that quantifies the information gain, immediately after an observation, that can attribute to an encoded tile *s*_encoded_:

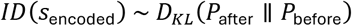

where *P*_before_ and *P*_after_ are belief distributions localized near *s*_encoded_. The amount of information gain is defined as KL-divergence between the two distributions. Lastly, we make this a density measure such that 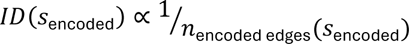. To unpack the above equation, we further defined four auxiliary functions: locality, ambiguity, identifiability, and fractional information gain which are discussed in Supplementary Materials in details. Intuitively, *ID* has a simple interpretation when performance is high and belief distribution is well localized:

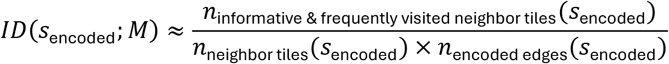

where computing *ID* comes down to simply counting how many unambiguous and frequently observed tiles around an encoded tiles vs counting how many encoded edges it takes to achieve such identifiability. It’s worth highlighting that the information density we define here depends only on a single observation at each step, and it is thus insensitive to the exact sequence prior to the observation. A more general higher order information density that depends on *k*-step history is beyond the scope of this paper. For full details regarding how and why we made certain modeling and algorithmic choices, see the Sparsemap Documentation in Supplementary Materials. Code is available at https://github.com/HermundstadLab/sparsemap.

### Sensory compressibility analysis

To quantify whether the paths chosen by the mice make the sensory information more predictable or compressible (Fig.4), we tested how much the sensory information in each upcoming sample in the equalized path (each sample is ∼4 cm) can be predicted from the previous sensory data, or sensory and motor data. To compute egocentric sensory data and motor descriptors, data from maze A was computed. Body position was registered to a map of maze tile heights using the same method as for the egocentric shape preference analysis, and at each sample we extracted a 3-element vector of height at the mouse’s head, and on the left and right side with a ∼4cm offset. Motor output was quantified as speed and path curvature. Each channel was z-scored across the whole recording. Compressibility was quantified via nearest-neighbor prediction (*72*), a continuous-valued estimate of statistical compressibility, in 1-hour chunks: Within each 1 hour segment, the data excluding the current point by ±96 cm was used as a library to predict future data. Each signal was delay-embedded (4 samples, with lags {0,3,6,9}) and forecast one step ahead by averaging the next steps of the closest matches in this library of ∼33 cm long trajectories: the predicted next state was the distance-weighted mean (weights ∝ exp(−distance to neighbor/ distance to nearest neighbor)) of its 8 nearest neighbors in the library, excluding neighbors within 24 samples / 96 cm of the current sample. Predictability was quantified via mean square error (*MSE*) as *C* = 1 − *MSE_pred_*/*MSE_surrogate_*, where *MSE_surrogate_* was obtained by applying the forecaster to phase-randomized surrogates of the same 1 hour chunk (amplitude-preserving, phase-scrambled with a shared random phase for all sensory and motor dimensions; median of 10 surrogates (*73*)). *C* therefore captures recurrent motif structure beyond the signal’s linear autocorrelation. Because of z-scoring, MSE measures normalized variance and predicting the mean yields MSE ≈ variance ≈ 1, while a perfect predictor gives MSE → 0. All predictability values *C* were converted to a Gaussian-equivalent predictive-information rate, −0.5 · *d* ·*log*₂(1 − *C*) bits per predicted step (*d* = 3 number of predicted dimensions), and reported per distance travelled (bits cm⁻¹). To keep bins comparable, each 1 hour bin estimate used a fixed sample N of 4000 query and 8000 library vectors, and bins with fewer than 2000 valid embedding points (mouse is moving enough in this bin) were excluded. To check the predictability of the sensory data given known motor data, we computed the conditional predictability as a partial variance reduction, computed as the error in predicting the next sensory sample *S_t_*_+1_from the sensory past *S_past_*, the motor past *M_past_*, and the current motor action *M_t_*_+1_, over the error of just predicting next sensory sample *S_t_*_+1_ from the sensory past *S_past_* as *C*_*S*|*M*_ = 1 – 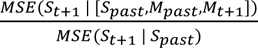. We used a large-lag motor time shift shuffle as control for the added dimensionality of the joint predictor. *C_S_*_|*M*_was converted to bits per cm by assuming normal distributed data, *I*_*S*|*M*_ = 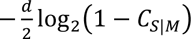 bits, giving an estimate of the conditional mutual information between the upcoming sensory sample and the motor state given the sensory past. This assumes that the forecast residuals are Gaussian and is approximate. Summing over the three sensory dimensions further assumes independent residuals. We therefore interpret these values as a relative measure across time and conditions rather than as absolute information rates.

## Supplementary Figures

**Fig. S1.**
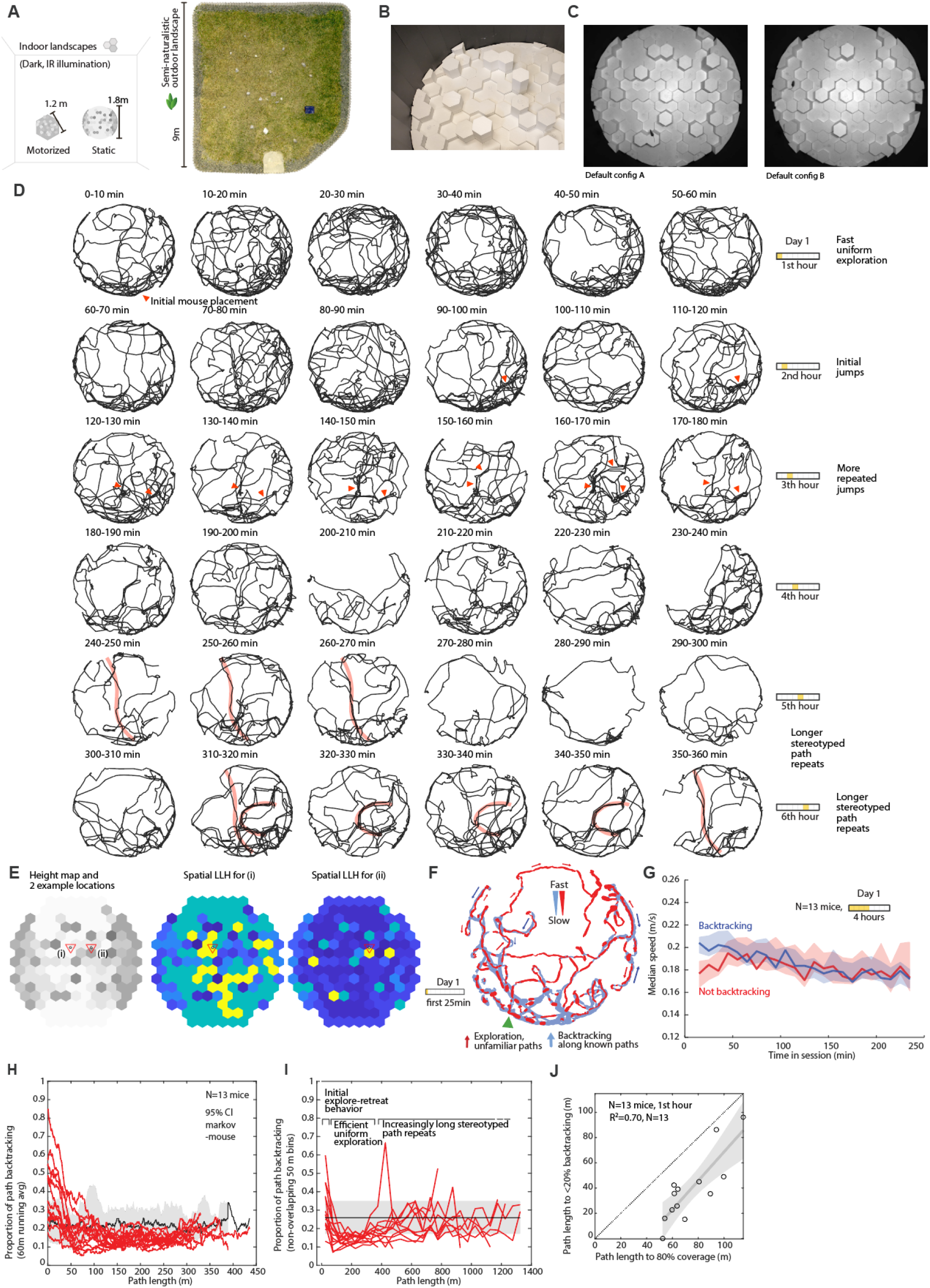
Dynamics of exploration behavior in large complex arenas. (**A**) To-scale size comparison of all three landscapes used in the study. (**B**) Photo of the large indoor landscape with room lights on. All experiments were conducted in darkness under IR illumination (See Methods). (**C**) Top-down views from the camera that was used for the behavior tracking under IR light, showing the default indoor landscape configurations ‘A’ and ‘B’. (**D**) The first 6 hours raw tracking data from one animal, plotted in 10 minute bins. (**E**) Examples of local sensory ambiguity resulting from the self-similarity of the maze. Two example locations are shown, with the likelihood resulting from an ideal Bayesian observer matching the three surrounding tile heights to the complete map of the arena (See methods). (**F**) Example raw data from the first 25 minutes of mouse exploration, same as in Video 1. Paths are classified as backtracking or exploration, and running speed is indicated as line width. (**G**) Summary plot of 13 mice (all mice with initial exploration in landscape ‘A’), showing that over the first ∼50 meters (∼30 minutes), mice explore outwards slowly and backtrack more quickly. (**H**) proportion of backtracking as function of distance run, running average of last 60 m. (**I**) same as H but in 50 m bins. The distinct phases of exploration are visible in this plot: Explore-retreat behavior leads to a lot of overlap with previous paths, then a phase of efficient uniform exploration avoids backtracking (N=14 mice, bins from 75-325 m P<0.0005 signed rank vs. shuffled null), and then mice start repeating stereotyped paths. (**J**) Distance at which backtracking ended is correlated with distance at which they cover 80% of the arena, indicating a per-mouse variation in exploratory behavior (N=13, r=0.83, slope p-value P=0.0004).

**Fig. S2.**
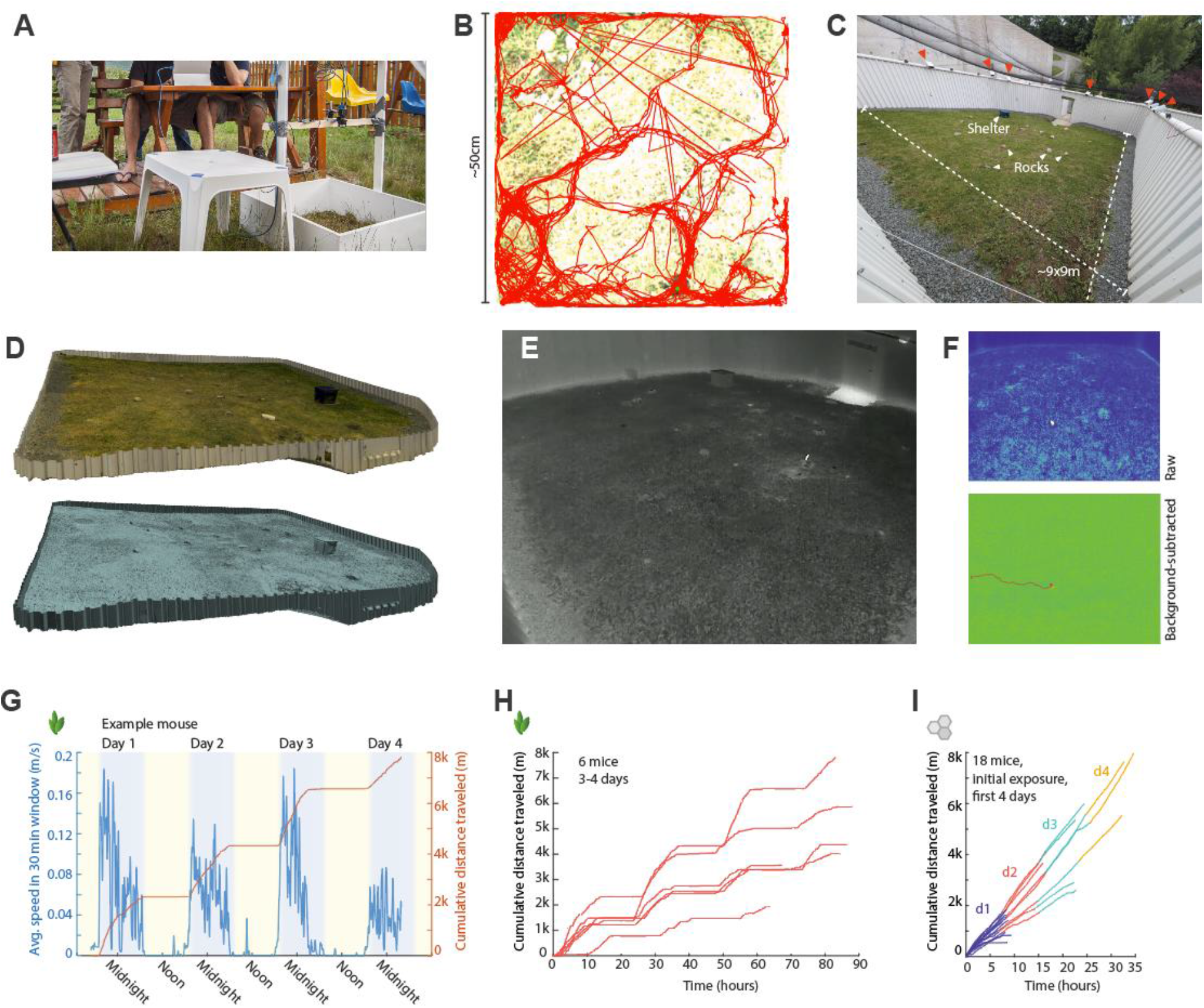
Mice explore outdoor enclosures. (**A**) Teaching assistants and students at the 2015 Transylvanian Experimental Neuroscience Summer School filming and tracking mice in a small outdoor enclosure (bottom right). (**B**) Tracks of a mouse from one of the recording sessions, showing early evidence of clustered paths. (**C**) One of the large outdoor enclosures at Janelia in 2025. Rocks, the shelter box, and some of the thermal cameras are indicated. (**D**) 3D-scan of the outdoor arena. (**E**) Example frame from one thermal camera showing a mouse rearing. Also see video 2. (**F**) Example raw and background-subtracted frames and mouse track. The resulting 2D tracks were then combined into 3D position estimates using anipose (*71*), see Methods for details. (**G**) Averaged smoothed speed and cumulative distance run by a mouse in the outdoor arena. Distance was computed after significant smoothing to exclude local search behavior and only quantify long-distance travel: x/y time series were smoothed with a running median in a 20 second window, and then cumulative distances were calculated from points every 20 seconds. (**H**) Same for all 6 mice. (**I**) Cumulative distance run for 18 mice in the large indoor landscape, for 8 hours daily, for up to 4 days, showing comparable distances. Distances were also computed after significant smoothing: x/y time series were low pass filtered at 0.1 Hz with a 3^rd^ order butterworth filter to exclude local exploration movement and to collapse periods of stationarity into single points despite tracking jitter and mouse movement.

**Fig. S3.**
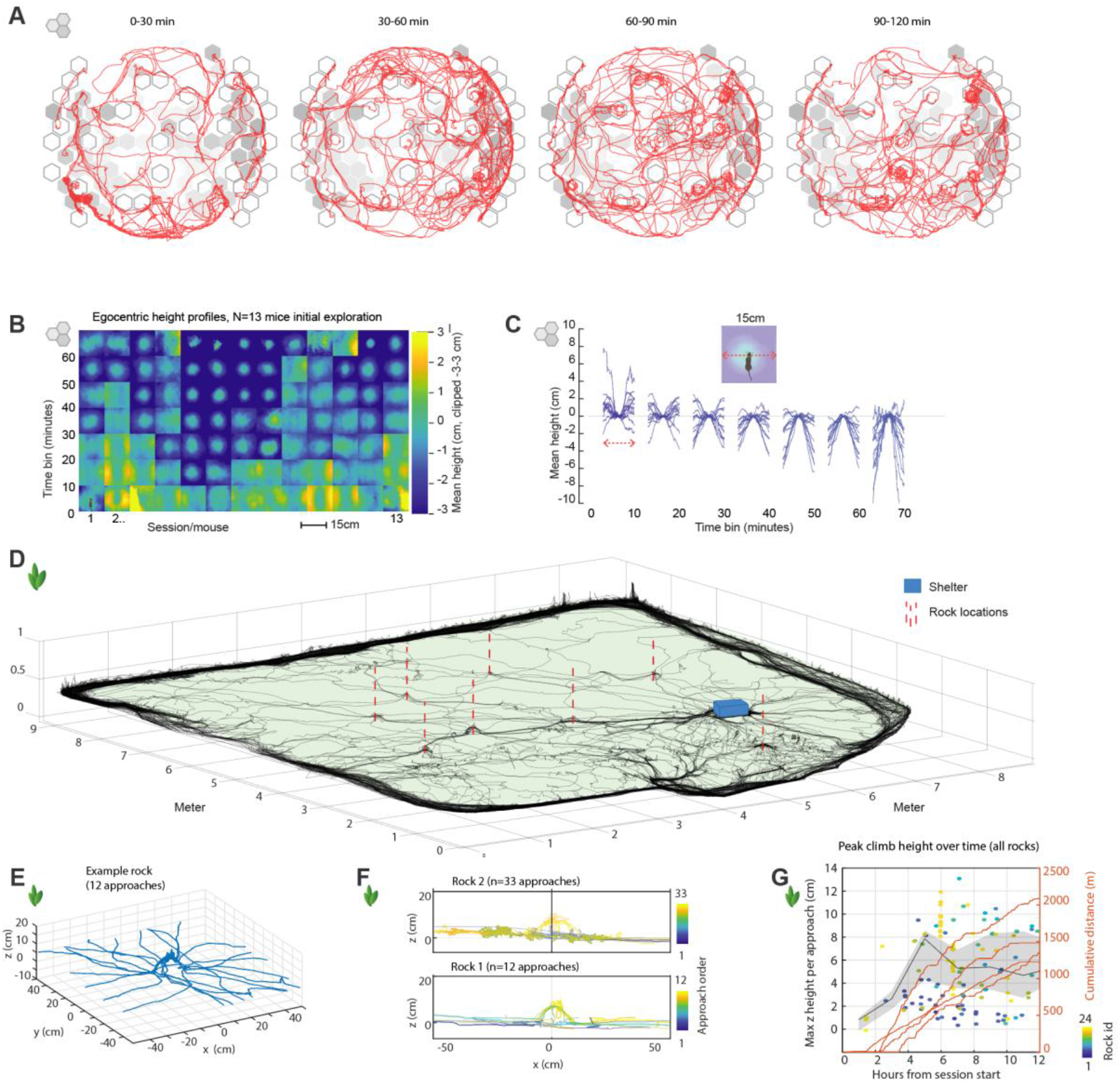
Mice quickly explore elevated locations. In typical lab experiments in boxes, rodents establish a home base location (*32*) and then make increasingly long excursions leading to full exploration of the box (*35*, *74*), while preferring peripheral paths that retain whisker contact with the wall (*34*) (termed ‘Thigmotaxis’). (**A**) Example trajectory of a mouse in the first 2 hours in the maze, showing initial avoidance of tall towers, and increasing time spent on high tower after ∼20-60 min. (**B**) In our indoor landscape mice quickly transitioned to spending time in tall places. Individual plots show average egocentric landscape shape around the mouse, aligned to the head position, in 10 minute bins, for all 13 mice initially exploring configuration A (See methods for details). (**C**) Plot of the same data as in B but as slice through the space from the left to the right of the mouse, individual plots are for different mice. (**D**) Example 3D plot of a mouse in the outdoor arena, with locations of large rocks (∼10 cm) indicated. This mouse not only used rocks as preferred ‘waypoints’, but walked over them rather than around them. (**E**) Example of all the approaches (<30 cm) to one of the rocks. (**F**) Example side view of the approaches to two example rocks, with each approach shaded by its number. Early approaches stick to the base of the rock, and later ones lead to traversing the top of the rock, or to standing and rearing on top. (**G**) Summary statistics from 4 sessions in which rocks were present and 3D tracking was clean enough to quantify height over ground (see Table 3). Dots indicate approaches, with their peak height over ground. Shaded area is a 95% CI of the median. Cumulative traces show meters covered by the 4 mice. Mice start to climb on rocks around 2-4 hrs into the night, after ∼500-1000 meters of exploration.

**Fig. S4.**
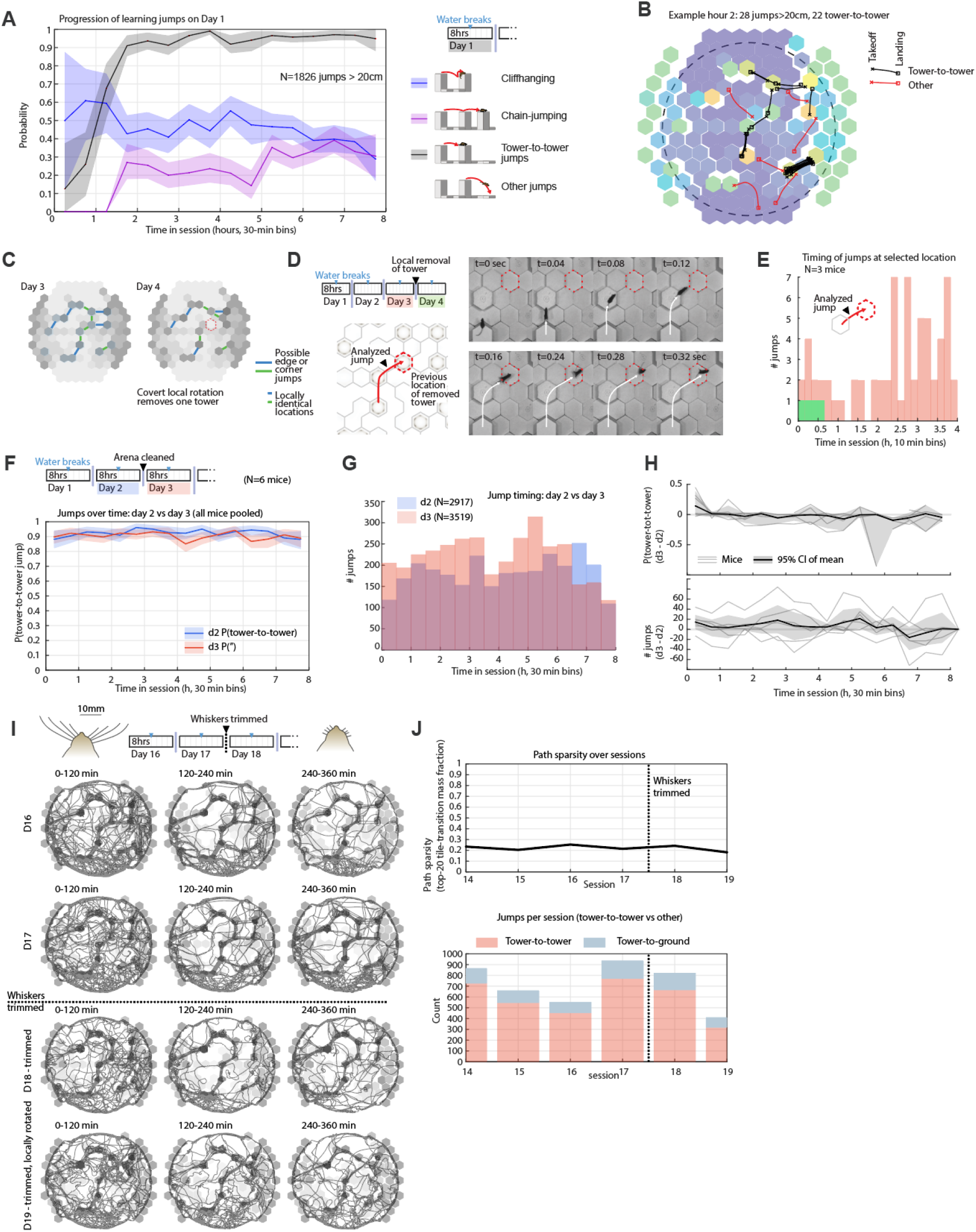
Mice jump across gaps from memory. (**A**) Statistic of jumps during the first day (N=13 mice with initial exploration in landscape A, same analysis as in Fig.1C), showing rapid learning of tower-to-tower jumps (grey), reduction of jumps in which mice hold on to the target ledge and then climb fully onto the target tower, ‘Cliffhanging’ (blue), and emergence of chain-jumping, jumps that follow each other by <1sec (purple). (**B**) example data from one hour of mouse showing 28 jumps, the dotted circle indicates the zone used to exclude jumps at the landscape periphery. (**C**) Schematic illustration of the default landscape ‘A’ (See Fig.S1B,C), with jumps between tall towers indicated by long lines (Blue: edge to edge, green: corner to corner), and short line segments indicating locations that appear identical on the tower itself, but do not lead to landing spots on other towers. On Day 4 we covertly rotated one block of tiles, removing one tower from the previous jump sequence. (**D**) Illustration of jump that is made impossible by rotation of the tiles, and frames from a subsequent failed jump showing that mice still attempt the jump a few times before re-learning that this jump is now impossible. Also see Video 3 for videos of these jumps. (**E**) Quantification of all jumps (tower-to-tower and jumps from towers on the jump direction indicated in D). Mice attempt the jump 1-2 times before completely abandoning this jump at the start of day 4. (**F**) In the main experiments we cleaned the landscape between days 2 and 3 (see Methods). The proportion of tower-to-tower jumps, stayed stable, indicating that jumps were not driven by olfactory cues. (**G**) The number of total jumps per 30min bin similarly stayed stable. (**H**) Per-mouse differences in metrics from F and G. (**I**) We trimmed one mouse’s vibrissae to ∼2mm after 17 days of exploration. Raw position traces show jumps before and after trimming. (**J**) Paths sparsity (Same method as in Fig.2) and Jump count across the whisker trimming, showing no significant reduction in jumping.

**Fig. S5.**
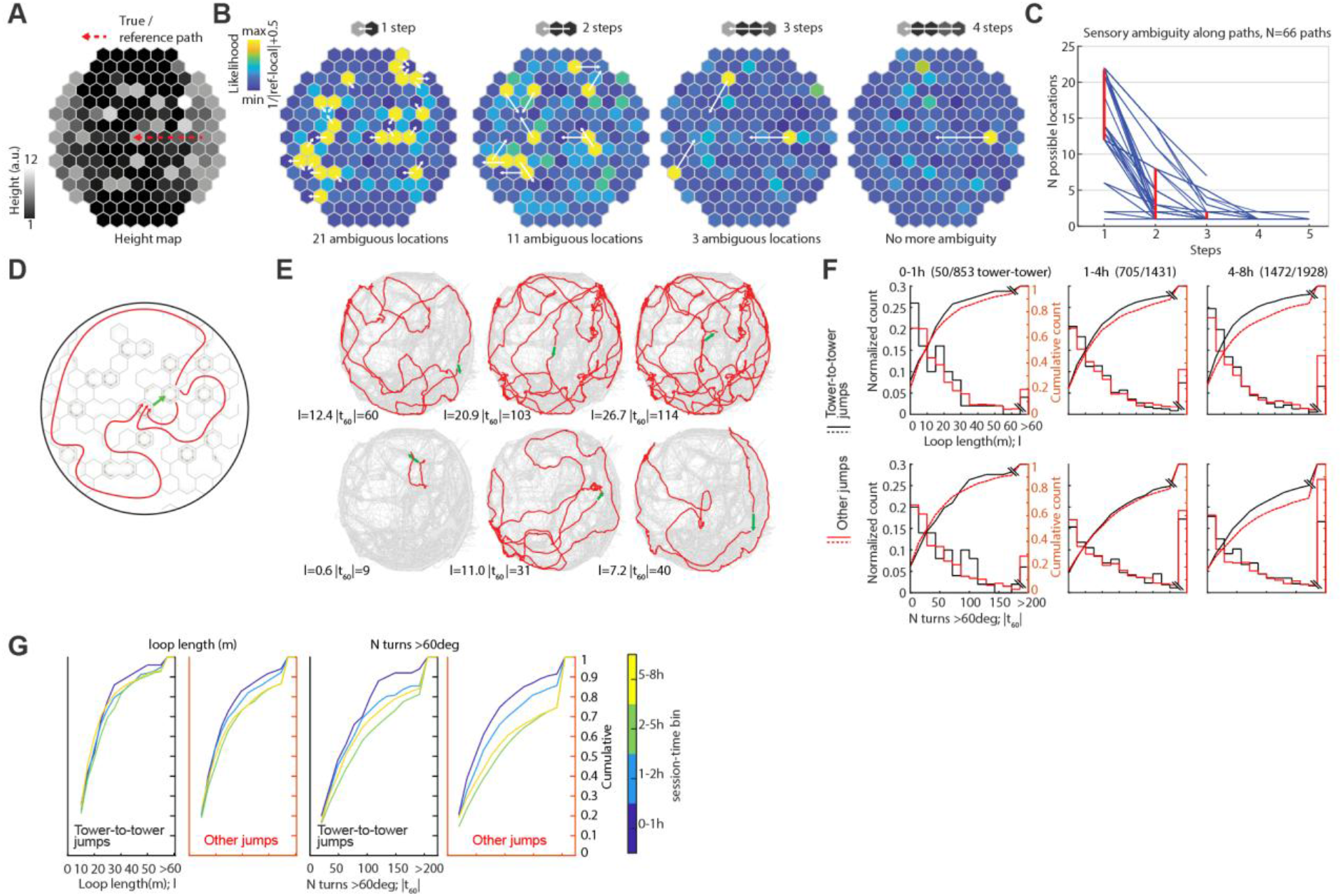
Jump locations are not directly identifiable by their shape, or via path-integration. (**A**) We tested if the shape of the landscape around the jumping-off point was unique enough to identify jump locations. We applied a Bayesian ideal observer model with perfect knowledge of self-motion and of the arena (*53*) backwards in time from jumping points and quantified when the observer converged on a unique location. (**B**) Example run of the ideal observer for one jump location and direction. It takes 4 full tile steps to uniquely identify the location. (**C**) Plot of the N of steps vs. ambiguous locations for 66 possible paths, even with perfect map knowledge, a local ideal Bayesian observer would need at least 3 full tile steps (∼45 cm) of memory to identify the jump locations. (**D**) Illustration of possible path-integration strategy. For any jump (green), mice could first visit the landing spot, then take a short loop back to the jumping-off point and then jump using a memorized landing location. If they cover a long distance since last visiting the landing spot for any given jump, this path-integration strategy becomes more susceptible to small path-integration errors. (**E**) example jumps and their path-integration loops, with their length, and number of >60 degree turns. (**F**) Summary statistics across mice, split by time from the first day, and tower-to-tower vs. other jumps. Histograms and cumulative distributions are plotted. (**G**) Same analysis as F, but only cumulative distributions are shown across time bins.

**Fig. S6.**
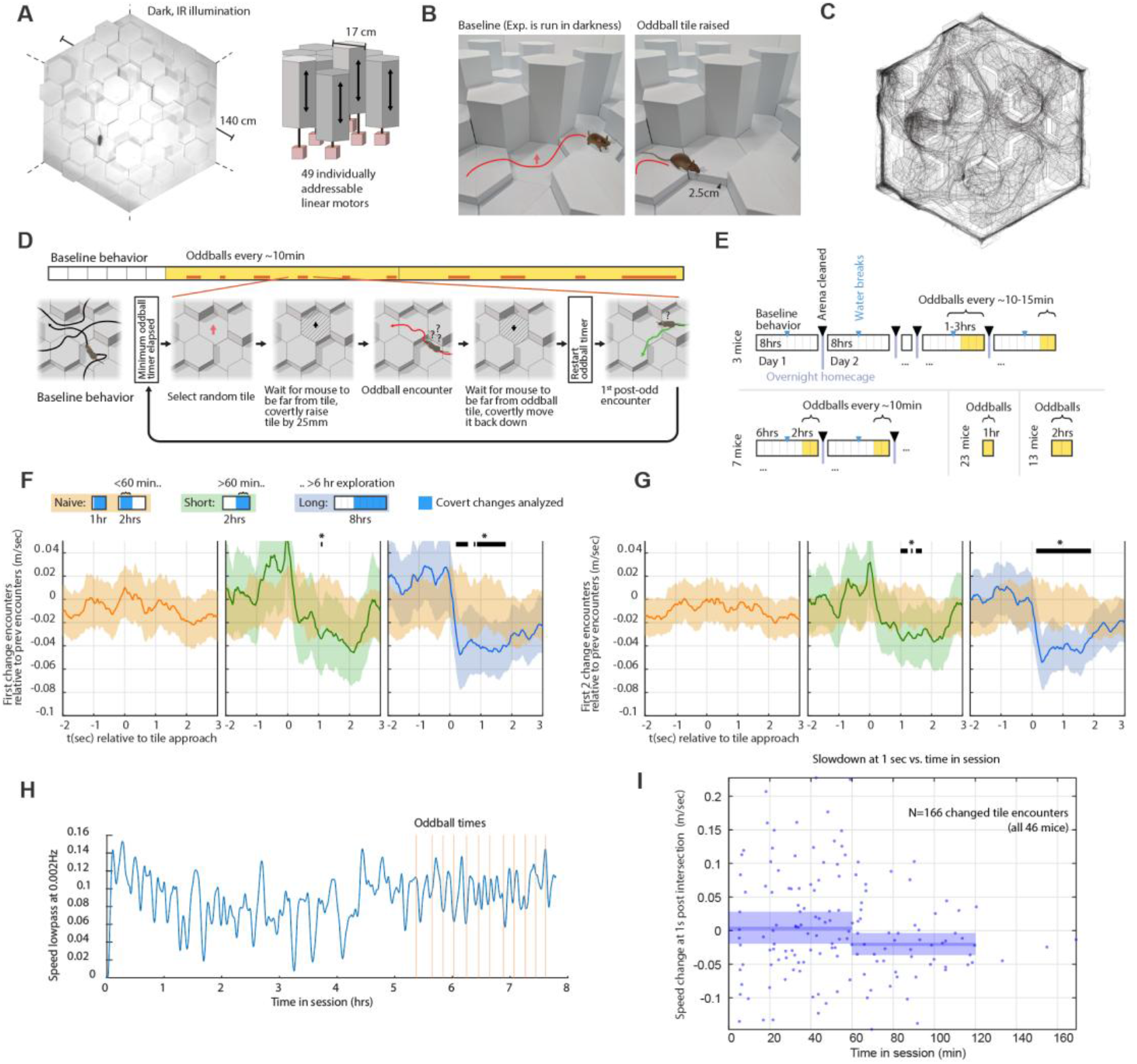
Mice detect small covert changes in a complex landscape. (**A**) Example frame from top-down camera using IR illumination. All experiments were run in darkness. The illustration shows the motor system, 49 of the 52 hexagonal prisms could be moved up or down with individual linear actuators. (**B**) Illustration of a covertly changed ‘oddball’ tile (raised by 25 mm), photos were taken with room lights on, and mouse illustration added to show approximate scale. (**C**) Example tracking data from one session in the reconfigurable landscape. (**D**) Logic of the covert change program (example shows a session with pure baseline behavior, we also ran sessions where oddballs started immediately). See Methods for more details. A tile is randomly selected from a subset of eligible tiles. Then the program keeps checking whether the mouse is sufficiently far away from the chosen tile, before raising it by 25mm. Then the program waits for the mouse to encounter the tile by (center of mass <13cm from tile center), and again waits for the mouse to be far away before lowering the tile to its original position. This procedure ensures that mice don’t ever encounter tiles moving and makes it impossible to localize movement via sound cues. The procedure results in a uneven timing of oddballs because in addition to the timer of e.g. 10 minutes between edits, the wait times for the mouse to be far from the tiles adds delays. The need for mice to be far from the tile before we lower it also results in >1 oddball encounters. (**E**) Overview of the experimental conditions leading to mice either having >6 hrs or 0-1, or 0-2 hrs of exploration time in the maze before covert changes. (**F**) Same analysis as in Fig. 1E, but only the first changed-tile encounter is analyzed (Naïve <60 min: N=106 encounters, 30 sessions, 60-120 min: N=49 encounters, 19 sessions, >6 hrs: N=155 encounters, 16 sessions). (**G**) Same analysis as in Fig. 1E, (Naïve: N=173, 60-20min: N=79, >6 hrs N=263 encounters). (**H**) Example session with mouse speed and oddball times. (**I**) Slowdown relative to baseline at 1 second post-encounter, as function of time in the session.

**Fig. S7.**
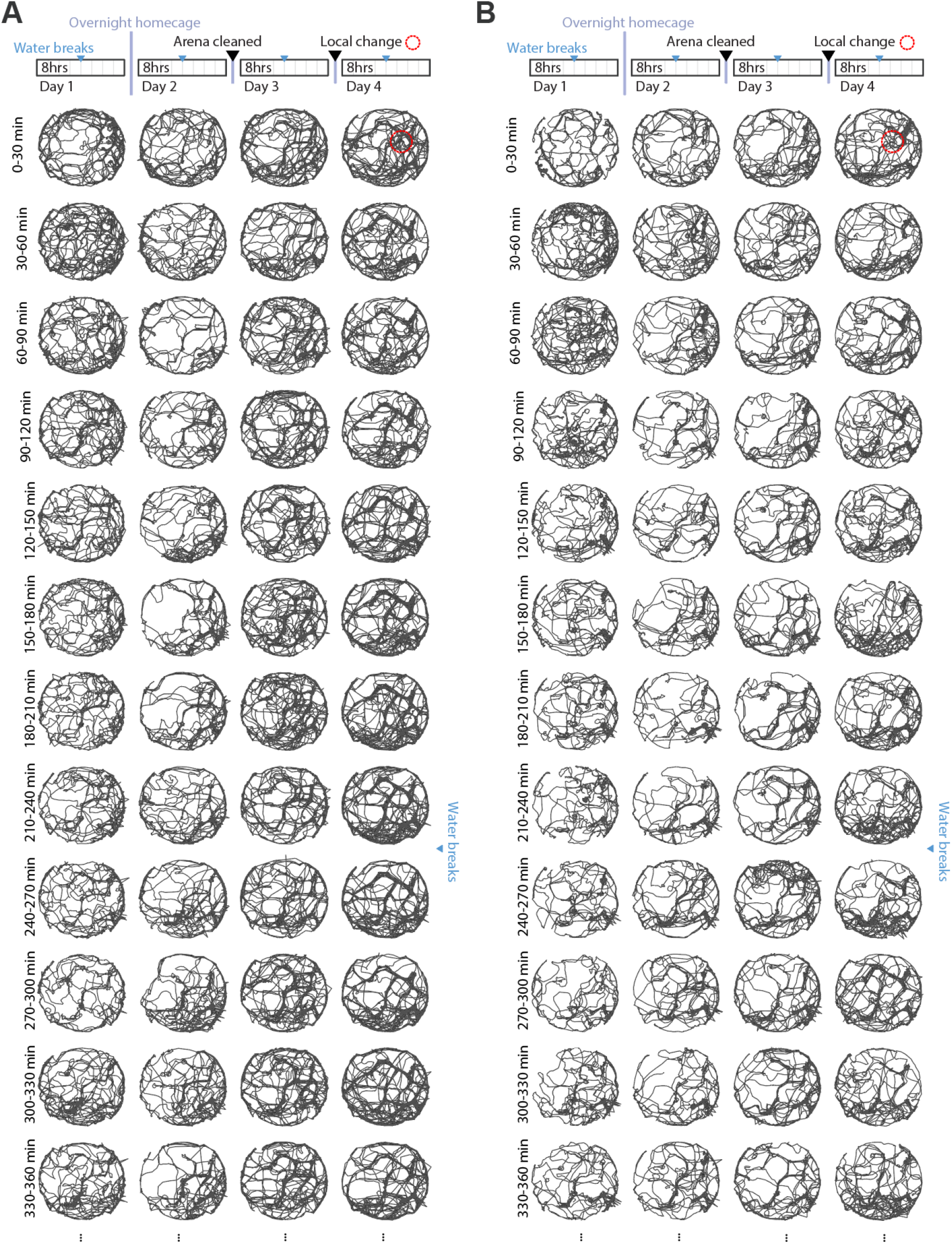
Exploration behavior in large indoor landscape across days. (**A**) Tracking data from the first 6 hours of one mouse across first 3 days in landscape configuration A and day 4 where we covertly rotated one set of 7 tiles. (**B**) Same for another mouse.

**Fig. S8.**
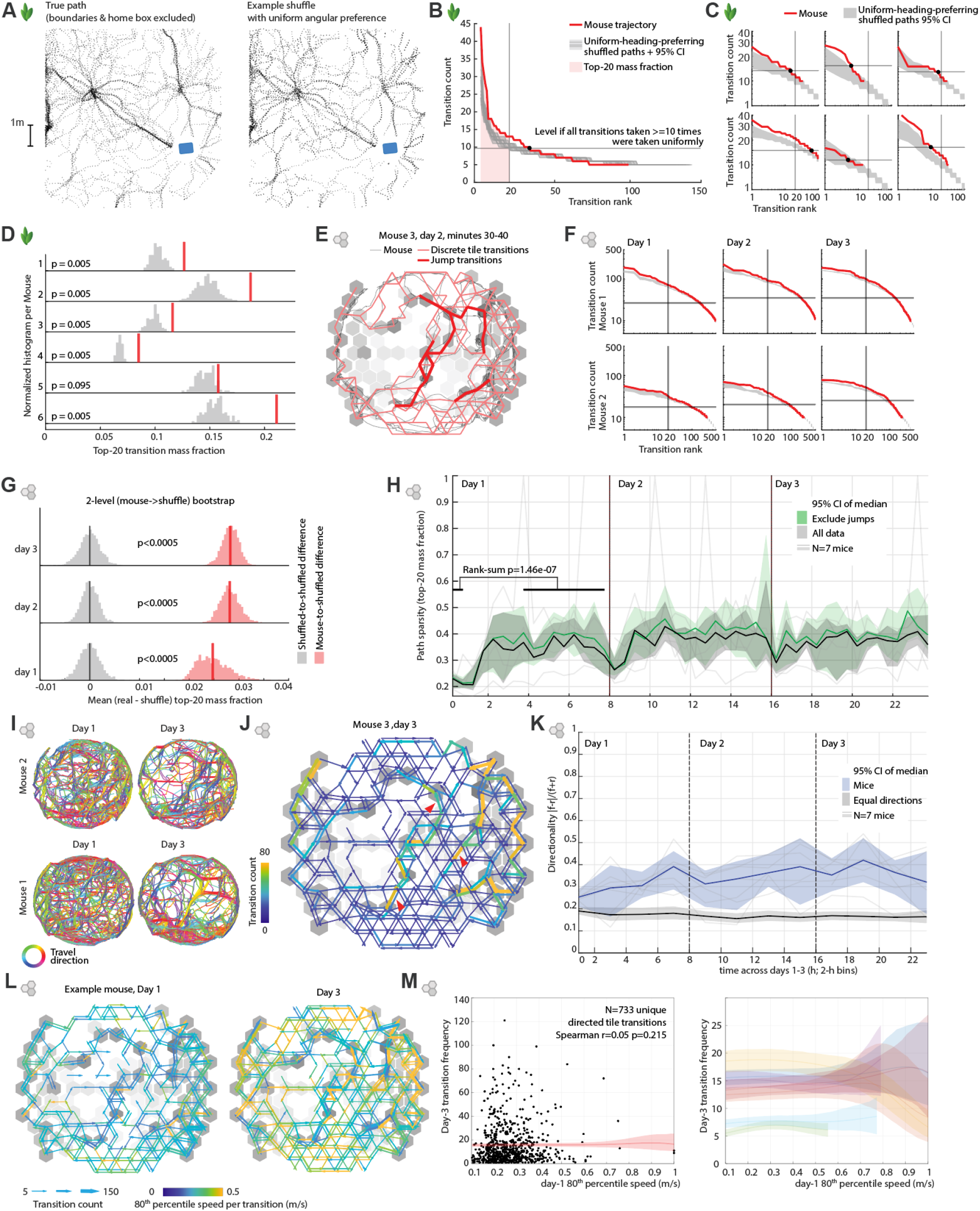
Mice converge on a sparse subset of paths. (**A**) In the outdoor arena we compare the true path to shuffled paths that jump between overlapping paths randomly, while preferring paths that avoid repeating the same heading (see Methods). (**B**) Example statistics for one mouse. Transitions between 50cm cartesian grid bins were counted in a 4-neighborhood maintaining direction between bins to generate a time series of discrete transitions. We then compared the count of these transitions against a flat distribution consisting of all transitions that are taken at least 10 times (horizontal line), and against a 95% CI of the shuffled paths (grey shaded regions). (**C**) same as B but for all 6 mice, axes are log scale for clarity. (**D**) Summary stats for all mice in the outdoor experiments, the distributions are quantified via the fraction taken up by the top 20 transitions for each path, P values are directly from the shuffled distribution. (**E**) For the indoor landscape, we discretized the path into the hexagonal tile layout (see Methods) and annotated each transition as belonging to a jump if it comes within a 1 sec window of a jump (same definition as for analyses of jumps, Fig.1). (**F**) same as C but for indoor experiments in large landscape, and broken up per day. (**G**) Summary statistics for inside experiments, using 20-top transition fraction, comparing within-shuffle to shuffle vs. real differences (N=7 mice, broken out by day), see Fig.2D for summary for outdoor data. (**H**) Top-20 transition fraction vs. time for all data, and excluding jumps. Transitions at landscape boundary are excluded in both conditions. (**I**) Example traces from two mice across days 1 and 3, colored by travel direction, similar to Fig.3C. (**J**) Example of directional transitions from day 3, color and arrow size show directed transition counts. Red arrows point to particularly directed example transitions. (**K**) Summary statistics for emergence of directional path preferences, in 2 hour bins. Blue: For each pair of tiles, directionality is computed as Σ|f−r| / Σ(f+r), with f and r being the count of forward and reverse transitions. Transitions with speed <0.02m/sec and at landscape boundary are excluded and only transitions with f+r>10 are analyzed. Grey: Basic null hypothesis, Binomial(N of forward and backwards, 0.5). All shaded regions are 95% CIs, either via 2 level bootstrap (mouse, transition), or inverse binomial. (**L**) Example days 1 and 3 from one mouse, showing transitions with count >5, thickness indicates count, color indicates the 80^th^ percentile speed. Data from the first hour of each day, and transitions of under 0.1m/sec speed were excluded. (**M**) Scatter plot of all transitions from this mouse, showing lack of relationship between speed on day 1 and subsequent use frequency. Shaded region shows kernel regression estimate 95% CI. Kernel regression 95% CIs for all 7 mice, showing overall lack of relationship.

**Fig. S9.**
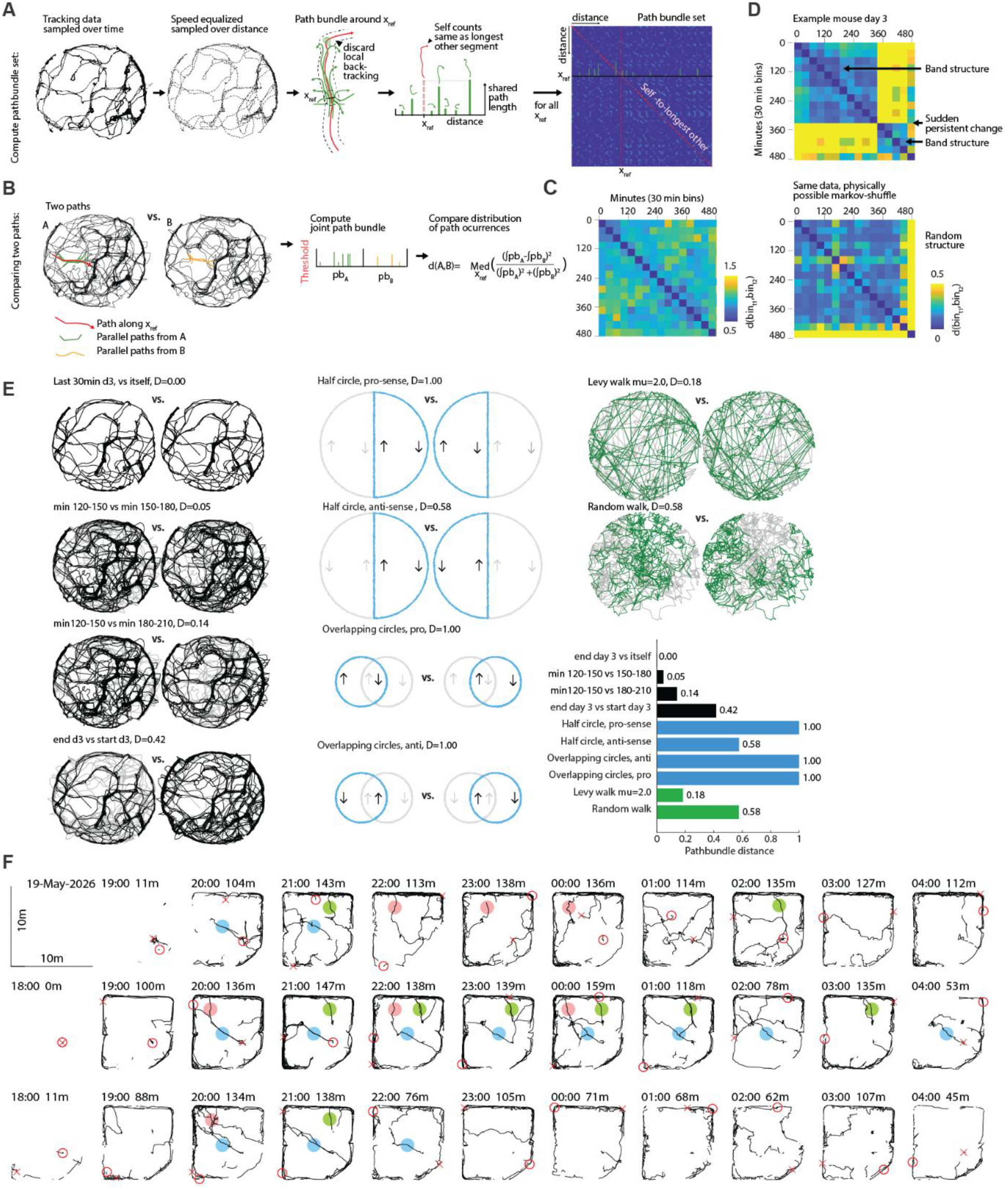
Path bundle method. To quantify the structure of repeated paths, and to compare two paths, e.g. segments of the same mouse at different times, or across mice, we use the ‘Path bundle’ method. (**A**) Computing the path bundle for a path. The path is first resampled so that every step is a fixed distance. We then compute for every point xref in the path the set of other (earlier or later) paths that approach the point and keep track of how far they stay with the current path segment. Points along the overlapping path segment that fold back over itself within 1.5*step-distance are not counted towards this length. The diagonal entry is set to the length of the longest other overlapping path segment. (**B**) We can use the pathbundle to compare the structure of two paths. We concatenate the paths, compute their joint pathbundle, and then compute the median of the normalized squared difference between the parts of the path bundle that belong to the two initial paths, across the entire joint path. See the methods section for a detailed description of the method. (**C**) Example path distance matrix for one day of data in the large indoor landscape evaluated across all 30-minute bins. (**D**) Top: Same as C but for another mouse and day, showing slow drift as well as a change in behavior towards the end of the day. Bottom: the same data but shuffled while maintaining path continuity, (same method as in Fig.1B), removes the temporal structure. (**E**) 10 example pairs of real or synthetic data and their pathbundle distance. All path pairs are 30 minute segments at the same scale as the mouse tracking data. Examples 1-4 show an example mouse tracking dataset, with increasing temporal offsets between the segments, the first pair is a repeated path resulting in a distance of zero. Examples 5-8 show synthetic data with paths that circle the arena and overlap along the mid-line, once pro and once anti-sense, resulting in the overlapping middle segment either registering as identical or not. The two smaller examples show that near-orthogonal crossings are not counted as similar regardless of travel direction, because the threshold cuts off the resulting short overlapping path bundles, resulting in the maximal distance of 1 in both cases. Examples 9 and 10 show two types of random walk, a Levy walk with long straight sections results in a smaller distance than a small fixed-step random walk because the levy walk results in aligned long paths, while the short step-size random walk results in only very short matched segments and is dominated by spatial occupancy. (**F**) Example tracks from the outdoor arena (Fig. 2), in one-hour bins, data from three nights is plotted. A subset of repeated paths is highlighted in different colors to show that repeated paths tend to occur in clusters at the scale of hours.

**Fig. S10.**
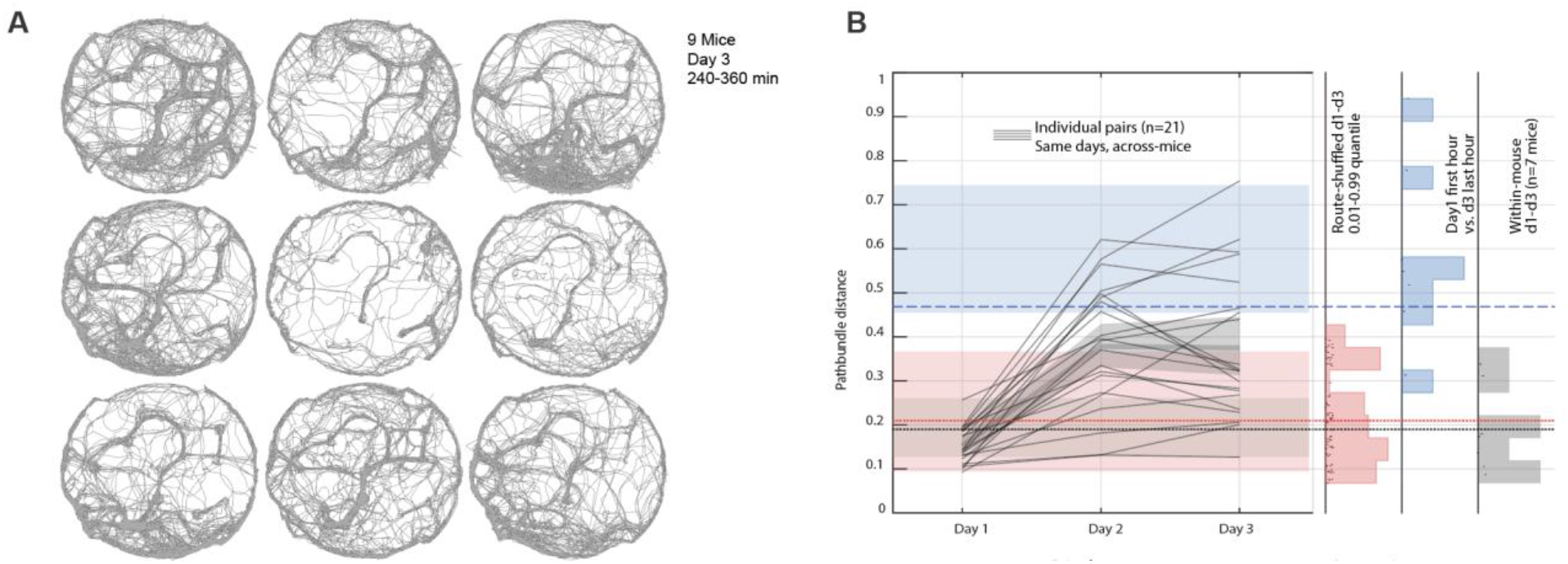
Mice converge on overlapping but diverse paths. (**A**) Example raw paths from 9 mice in landscape ‘A’, showing both individual preferences for specific paths, as well as patterns that are shared between mice. (**B**) Summary statistics (margins of the landscape are excluded from this analysis): Lines show pathbundle distances for the same days across different mice (N=21 pairs). We compute three control conditions. Black: Within-mouse differences across days 1-3 show that individual mice diverge into separate path preferences relative to the 1^st^ day. Blue: The within-mouse difference between hour 1 of day 1 and hour 8 of day 3 shows an estimate of the maximum difference in path vocabulary that each mouse can exhibit. Red: Shuffling the paths (same method as in Fig. 1B) within entire days, generating ∼8 hours of shuffled data, and plotting the 90^th^ percentile of the resulting differences gives an estimate of the upper bound of difference that can be generated within each day by taking the same paths re-sampled at the same frequencies.

**Fig. S11.**
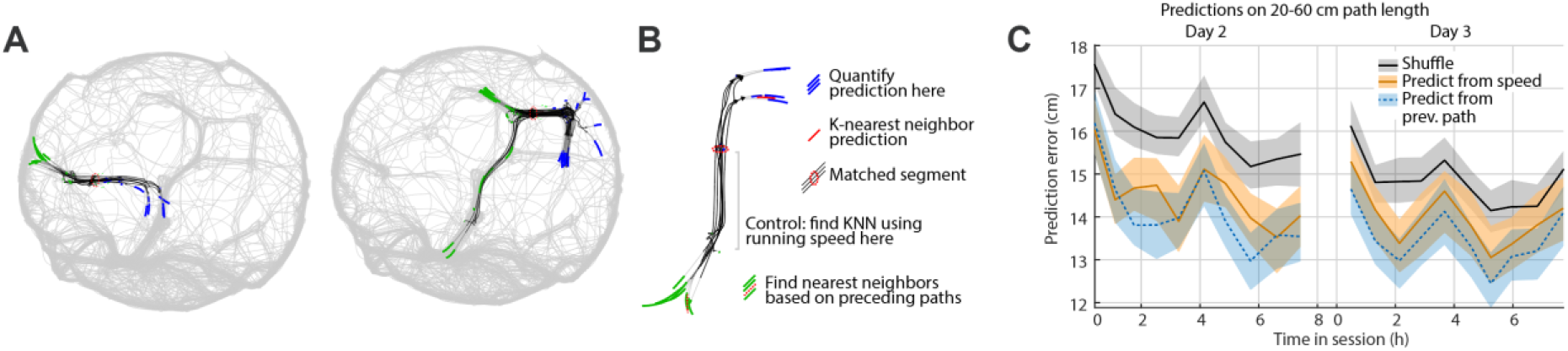
Paths are predictable across matched segments. To test whether paths decisions were driven by an internal state (Fig. 3I, this could be thought of as a memory state, navigation goal, current path preference, etc.), we identified times when the same path segment was repeated at least 6 times within a day, for a length of >20cm (See methods). (**A**) Two example sets of matched paths. (**B**) Example set of matched paths, with legend showing the preceding and following path segments used for the KNN prediction. (**C**) Prediction performance, same analysis as Fig.3I, but split over days 2 and 3, by hour, showing that prediction accuracy increased over time.

**Fig. S12.**
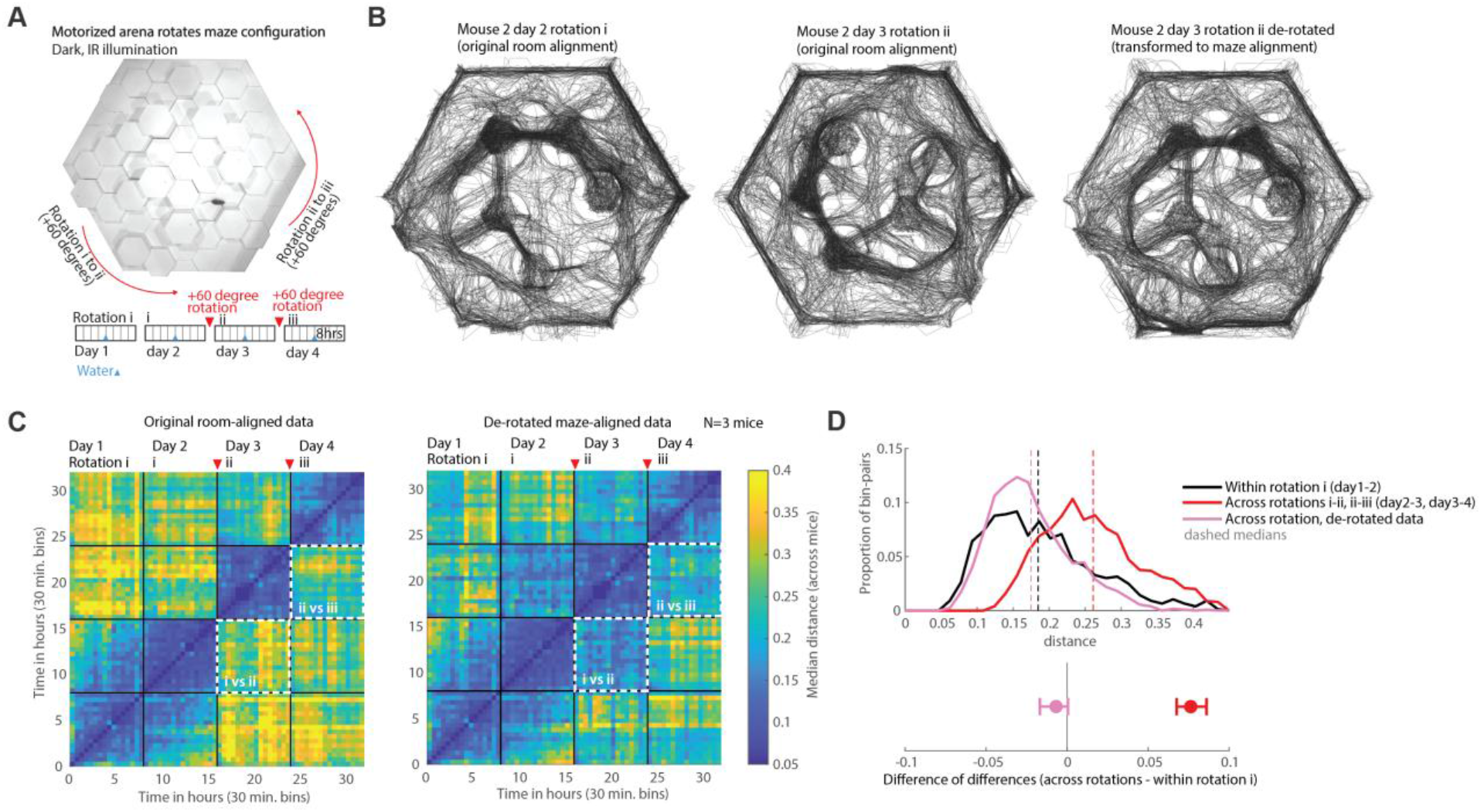
Paths are driven by the shape of the landscape, not by room-centric cues. To test whether the path preference was driven by unavoidable room-centric cues (memory of location where the door to the arena walls is, localized noise from the room air conditioning, air movement), we ran a control cohort of mice (N=3) in the reconfigurable landscape and rotated the shape of the landscape via the motors through three orientations (i/original, then +60 degrees to ii, then another +60 degrees to iii). (**A**) Top-down view of the landscape, and schematic of the experiment. (**B**) Example data from days 2 and 3 of a mouse in original room coordinates, showing a rotated pattern of paths, and the day 3 pattern de-rotated to bring it in alignment with the day 2 pattern. (**C**) Path similarity analysis via path-bundle distance (See Methods, Fig. S9) of the not de-rotated, room-aligned data (left), and the de-rotated data (right). Highlighted areas show alignment of paths structure between days 2 vs. 3, and 3 vs. 4. (**D**) Summary statistics showing rotated (D2 vs. D3 and D3 vs. D4) distances, compared to within-rotation control (Days 1 vs. 2) in red and black. Distances after de-rotation are shown in pink. Bottom: 95% Cis of the median, de-rotating to the shape of the landscape removes all path variance despite the now moving room-centric cues, showing that paths are not significantly affected by room-centric cues.

**Fig. S13.**
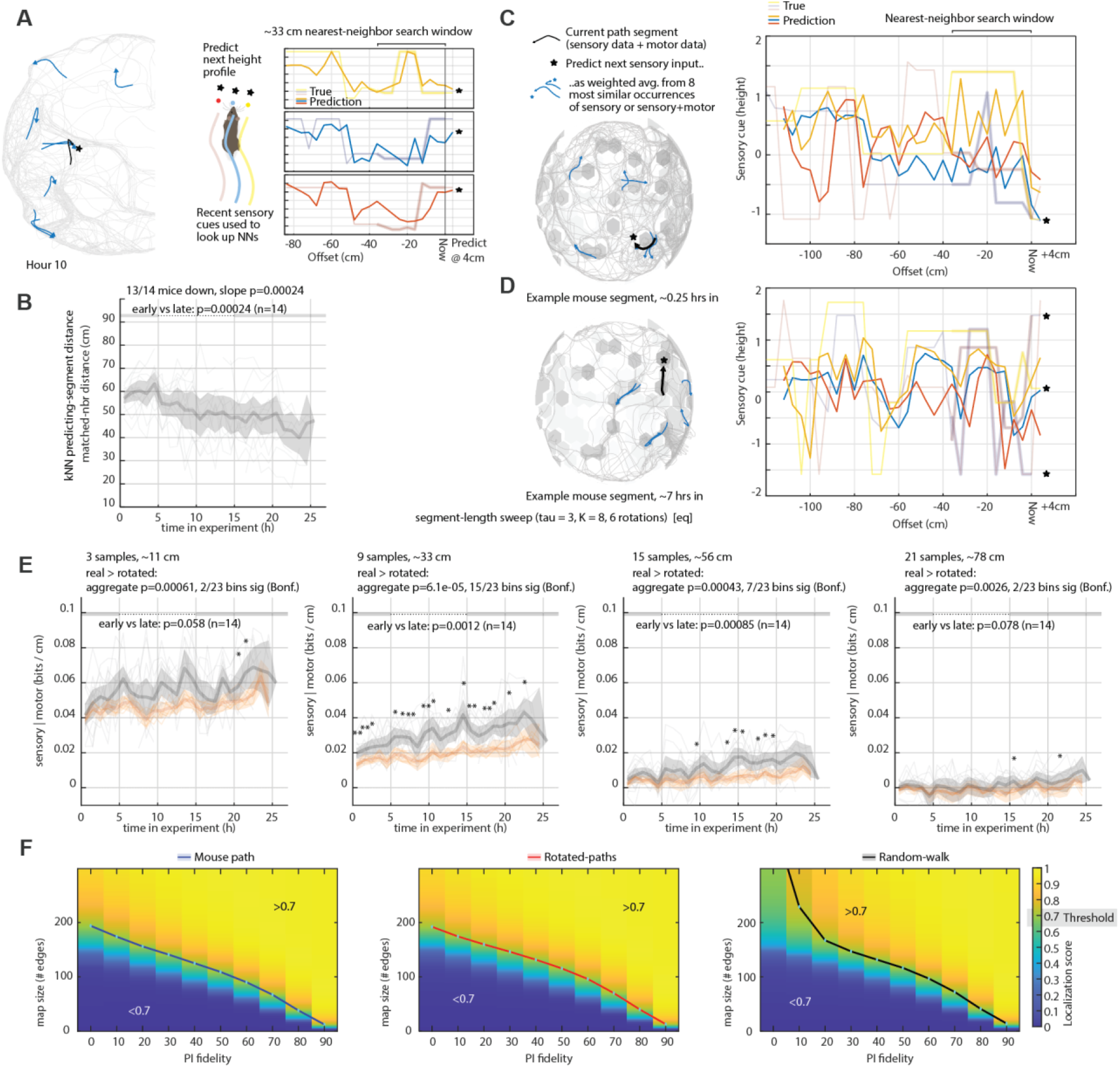
Sensory forecasting and map compressibility. To test whether the sensory input mice encounter becomes more predictable from their own recent experience as they form a map, we used nearest-neighbor (analog) forecasting (*72*) on the egocentric sensory stream. **(A)** For each mouse, the maze height at the mouse’s position and one tile to either side was sampled along the space-equalized path (one sample per ∼4 cm, moving epochs only), giving a three-channel sensory time series; speed and path curvature formed a two-channel motor series. At each point, the current path segment is described by its last 33 cm of sensory (or sensory + motor) data. The eight most similar segments from the same 1-h window of data, excluding temporally adjacent samples, are found (left, blue segments on the maze), and the sensory input 4 cm ahead (star) is predicted as their distance-weighted average. Right panels show the true sensory input (light traces) and the one-step predictions (dark traces) for an example mouse. (**B**) Physical distance between each predicted segment and the eight matched neighbor segments used to predict it (median per 1-h bin; grey, individual mice; black, cohort mean with bootstrap 95% CI). The predicting segments come from increasingly nearby locations over the experiment, indicating that sensory predictability is increasingly supported by revisits of the same routes rather than by similar-looking stretches elsewhere in the maze. (**C**) same as A but other example (**D**) same as C but later in the session. (**E**) Parameter sweep over length of nearest-neighbor segments, from 11-78 cm. The 33 cm chosen based on the approximate length of paths over which paths are predictable (Fig.3I) shows the clearest separation between true and rotated paths, but longer sequences still show the same effect. (**F**) Example heat maps from one mouse showing how the map sizes for a localization score of 0.7 are computed (Fig.4B&C). See Methods and supplementary materials for more information on how the sparse map results are computed.

## Supplementary Text

**Sparsemap documentation.** A complete documentation that expands on Map compressibility analysis in Method regarding why we made certain modeling & optimization choices, and about those earlier attempts that we decide not to include in the final results.

## Supplementary Videos

**Video 1. Initial exploration backtracking behavior.** Sped up trajectory of one mouse over the first ∼10 minutes, showing explore-retreat behavior, and backtracking along outward paths without use of shortcuts.

**Video 2. Mice outdoors climb on rocks.** Example unprocessed thermal videos showing example data of mice exploring the outdoor enclosure and jumping onto rocks. Video is in real time, but edited by selecting periods that show exploration behavior. Some rocks are annotated briefly to aid visibility.

**Video 3. Jumps.** Raw infrared video (experiment was run in darkness) of example jumps from two mice on their first day of exploration, and failed jumps and subsequent behavior after covert removal of one tower. Parts of the video are slowed down.

**Video 4. Exploration behavior in reconfigurable landscape.** Raw infrared video (experiment was run in darkness) of one mouse’s first exposure to the reconfigurable landscape. The video is sped up to show more behavior.

**Video 5. Detection of covert changes.** Raw infrared video (experiment was run in darkness) of a mouse’s first 4 experiences of covert changes to the arena. For each change, we show the selected tile (red arrow), then a baseline pass of the mouse over that tile, then we move the tile once the mouse moves away from the tile (black arrow), then the mouse’s first encounter with the now slightly elevated tile. In the first three instances, the mouse slows down to investigate the tile, in the fourth, it attempts to jump onto the tile unsuccessfully.

**Video 6. Exploration in large indoor landscape.** Raw infrared video (experiment was run in darkness) of a mouse’s first exposure to the large indoor landscape on day 1 and initial exploration on day 2. Parts of the video are sped up to show more exploration behavior.

