## Supplementary material for "Mice navigate complex landscapes by learning a sparse vocabulary of paths": Sparsemap documentation

To run the code, visit our repository: github.com/HermundstadLab/sparsemap

### 1. Glossary

| group | name | math | code | type, shape | value | note |
|---|---|---|---|---|---|---|
| | self position coordinate | $\mathbf{r} = (x, y)$ | `x,y` | `float` | `[0, 2000]` | |
| | normalized coordinate | $x_n, y_n \equiv x \,/\, d_{\text{tile}}, y \,/\, d_{\text{tile}}$ where $d_{\text{tile}}$ is the side-to-side tile width | `xn,yn` | `float` | `[0,13]` | |
| | tile position | $s$ | `s` | `int` | `[0,154]` | |
| | tile trajectory | $\mathbf{s} = (s_0, s_1, \ldots)$ | `s_traj` | `int, list` | `[0,154]` | |
| | heading | $h$ | `h` | `int` | `[0,5]` | |
| | action | $a$ | `a` | `int` | `[0,5]` | |
| | action trajectory | $\mathbf{a} = (a_0, a_1, \ldots)$ | `a_traj` | `int, list` | `[0,5]` | |
| | pose array | $X(s,h)$ | | `bool, (155, 6)` | `{0,1}` | one-hot joint encoding of tile position and heading |
| | belief array | $P(s,h)$ | `pq` | `float, (155, 6)` | `[0,1]` | joint probability distribution of tile position and heading |
| tile walk | heading transition tensor | $H_{\text{none},\, h^t,\, \text{none},\, h^{t+1}}$ | | `bool, (1,6,1,6)` | `{0,1}` | 1) two dummy dimensions that preserves tile position<br>2) note that we use superscript for naming indices in tensor notation rather than stating power. |
| | heading transition operation | | `np.roll` | | | this is an operation equivalent to $H$ |
| | tile transition dictionary | | `sas_dict` | `{(s0, a0): s1, ...}` | `([0,154], [0,5]): [0,154]` | mapping current tile-position-action pair to a target tile position |
| | tile transition tensor | $T_{s^t,\, a^t,\, s^{t+1},\, \text{none}}$ | `T` | `bool, (155, 6, 155, 1)` | `{0,1}` | 1) transition is deterministic<br>2) last dimension is a dummy that preserves the input heading<br>3) note that we use superscript for naming indices in tensor notation rather than stating power. |
| | tile transition tensor with boundary rule | $T_{s^t,\, a^t,\, s^{t+1},\, \text{none}}$ | `T` | `float, (155, 6, 155, 1)` | `{0,1,1/155}` | $T_{s^t, a^t, s^{t+1}, \text{none}} = \begin{cases} 1 & \text{if } a^t = a_{\text{phy}} \\ 1/155 & \text{if } a^t = a_{\text{unphy}} \\ 0 & \text{otherwise} \end{cases}$ |

| | | | | | | |
|---|---|---|---|---|---|---|
| | tile-heading index | $(s, h)$ | `(s,h)` | `int, tuple` | `[0,154]`,<br>`[0,5]` | 1) this 2D index running from (0,0), (0,1), ... to (154,5) marks all edges<br>2) note that a pair of overlapping edges connecting two neighbor tiles are labeled differently given which tile the agent is on. |
| | edge index | $e$ | `e` | `int` | `[0,929]` | this is a 1D index equivalent to its 2D counterpart: $(s, h)$ |
| | edge index mapping | $E(e) = \text{divmod}(e, 6)$<br>$\to (s, h)$ | `e_sh_arr` | `int, (930, 2)` | `col0: [0,154]`,<br>`col1: [0,5]` | |
| map | edge feature (relative height) array | $Z$ | `sh_dz_arr` | `int, (155, 6)` | `[-4,4]` | relative height of two neighbor tiles ranging from -12 to 12 is clipped at magnitude 4. |
| | sparse map | $M$ with a map size $m$ | `map` | `int, list` | `[0,929]` | a list of encoded edges |
| | wall-only map | $M_{\text{wall}}$ | `wallmap` | `int, (90, )` | `[0,929]` | a list of 90 encoded wall edges (each belongs to only one tile) |
| | random map | $M_{\text{rand}}$ | `randmap` | `int, list` | `[0,929]` | a list of randomly selected edges |
| | optimized map | $M_{\text{opt}}$ | `optmap` | `int, list` | `[0,929]` | a list of optimally selected edges |
| | heading diffusion kernel | $D(h, h';\ \kappa)$ | `pi` | `float, (6,6)` | `[0,1]` | this kernel consists of discretized Vonmises distributions with parameter $\kappa$ ranging from 0 (uniform distributions) to 100 (one-hot distributions) |
| | path integrator fidelity | $\kappa$ | `pi_level` | `int` | `[0,100]` | note that $\kappa$ is not the original parameter in Vonmises distribution, but a scaled version to ensure linear progression of standard deviation |
| inference | ambiguous edge mask | $C_{\text{amb}}(s, h;\ \Delta z)$ | `pq_mask_amb` | `bool, (155, 6)` | `{0,1}` | 1) the mask is a function $\Delta z$.<br>2) value 1 encode ambiguous edges with identical $\Delta z$. |
| | unknown edge mask | $C_{\text{unk}}(s, h;\ M)$ | `pq_mask_unk` | `bool, (155, 6)` | `{0,1}` | the mask marks unknown edges not encoded by a given map |
| | total edge mask | $C = C_{\text{amb}} + C_{\text{unk}}$ | `pq_mask` | `bool, (155, 6)` | `{0,1}` | 1) total mask is boolean sum of $C_{\text{amb}}$ and $C_{\text{unk}}$.<br>2) |
| | tile occupancy map | $O(s;\ \mathbf{s})$ | `occ_map` | `float, (155, )` | `[0,1]` | a probability distribution over all tiles given a trajectory |
| | highly occupied tiles | $O^*(s;\ \mathbf{s}) = \begin{cases} 1 & \text{if } O(s) > \text{median}(O) \\ 0 & \text{otherwise} \end{cases}$ | `s_top` | `bool, (155, )` | `{0,1}` | |
| evaluation | score map | $S(s;\ \mathbf{s}, M)$ | `score_map` | `float, (155, )` | `[0,1]` | for a given tile $s$, the tile score $S(s)$ is the average belief value at ground truth position $s$ over entire trajectory |
| | score | $\overline{S}(\mathbf{s}, M)$<br>$\equiv 2\sum_s S(s;\ \mathbf{s}, M) O^*(s;\ \mathbf{s})$ | `score` | `float` | `[0,1]` | the overall score is the average over top half occupied tiles. |
| | compression rate | $CR_x \equiv \frac{m_{\text{baseline}}\left(\overline{S}=\overline{S}^{(\text{target})}\right)}{m_{\text{optimization } x}\left(\overline{S}=\overline{S}^{(\text{target})}\right)}$ | `cr` | `float` | `[1,inf]` | |

| | | | | | | |
|---|---|---|---|---|---|---|
| map optimization | trajectory chunk | $c_i = (s_{il}, s_{il+1},..., s_{(i+1)l-1})$ | traj_chunk | int, list | [0,154] | $i$ is chunk index<br>$l$ is chunk length |
| | optimization schedule | | schedule | {iter: (n_chunks, chunk_id, chunksize), ...} | [0, 929]: ([1,37], [0,36], [500, 20,000]) | |
| | deletion based optimizer | | map_optimizer | | | 1) input: $\mathbf{s}$, $\mathbf{a}$, $D$, schedule<br>2) output: a list of 930 maps, ranging from the full map to empty map<br>3) each map in the list is one edge less than its predecessor. |
| information density | belief diffusion radius | $\sigma(s;\ \mathbf{s}, M)$ s.t.<br>$f(\sigma; S)$<br>$\equiv \sum_{s'} e^{-(\mathbf{r}_s - \mathbf{r}_{s'})^2/(2\sigma^2)} - \frac{1}{S(s; \mathbf{s}, M)} = 0$ | sigma_tile | float, (155,) | [0,inf] | a tile is considered to be a "soft neighbor" of $s$ if it falls within belief diffusion radius |
| | tile locality | $L(s, s'; \sigma) \equiv \frac{e^{-(\mathbf{r}_s - \mathbf{r}_{s'})^2/(2\sigma^2)}}{\sum_{s'} e^{-(\mathbf{r}_s - \mathbf{r}_{s'})^2/(2\sigma^2)}}$ | L_mat | float, (155,155) | [0,1] | $\mathbf{r}_s$ is the normalized coordinate for the center of tile $s$. |
| | masked edge feature | $Z_M(s, h;\ M) =$<br>$\begin{cases} Z(s,h) & \text{if } (s,h) \in M \\ \text{nan} & \text{if } (s,h) \notin M \end{cases}$ | sh_dz_map | int, (155, 6) | [-4,4] or nan | |
| | edge feature distance | $D_Z(s, s';\ M)$<br>$\equiv \min_{\theta} \sum_{\substack{h \text{ s.t.} \\ Z_M(s',h) \\ \neq \text{nan}}} \lvert \text{roll}(Z(s,h), \theta) - Z_M(s', h; M) \rvert$ | d_mat_dz | int, (155,155) | [0,48] | an observed tile $s$ is not distinguishable with $s'$ in map $M$ if their edge features are identical up to a 6-fold rotation:<br>$Z(s) = \text{roll}(Z_M(s'), \theta \in [0,5])$ |
| | tile ambiguity | $A(s, s';\ M) = \begin{cases} 1 & \text{if } D_Z(s, s'; M) = 0 \\ 0 & \text{if } D_Z(s, s'; M) > 0 \end{cases}$ | A_mat | bool, (155,155) | {0,1} | |
| | local tile ambiguity | $A_{\text{local}}(s, s';\ \sigma, M)$<br>$= \frac{L(s,s';\sigma)A(s,s';M)}{\sum_{s'} L(s,s';\sigma)A(s,s';M)}$ | | float, (155,155) | [0,1] | |
| | tile identifiability | $I(s, s';\ M) \equiv 1 - A$ | | bool, (155,155) | {0,1} | |
| | local tile identifiability | $I_{\text{local}}(s, s'; \sigma, M)$<br>$= \frac{L(s,s';\sigma)I(s,s';M)}{\sum_{s'} L(s,s';\sigma)I(s,s';M)}$ | I_mat_local | float, (155,155) | [0,1] | |
| | information gain on observed tile $s$ | $IG(s;\ \sigma, M)$<br>$\equiv D_{KL}(A_{\text{local}} \parallel L)$<br>$= -\log \sum_{s'} L(s, s'; \sigma) A(s, s'; M)$ | IG | float, (155) | [0,inf] | |
| | maximum information gain | $IG_{\text{max}}(s;\ S)$<br>$\equiv D_{KL}(\delta(s, s') \parallel L)$<br>$= -\log S(s; \mathbf{s}, M)$ | IG_max | float, (155) | [0,inf] | |
| | normalized information gain | $IG_{\text{norm}}(s;\ S, \sigma, M) \equiv \frac{IG(s;\sigma,M)}{IG_{\text{max}}(s;S)}$ | IG_norm | float, (155) | [0,inf] | |
| | fractional information gain from identifiable tile $s'$ | $FIG(s, s';\ S, \sigma, M)$<br>$\equiv I_{\text{local}}(s, s'; \sigma, M) \otimes IG_{\text{norm}}(s; S, \sigma, M)$ | FIG_mat | float, (155,155) | | $FIG$ attributes $IG$ by observing $s$ to those tiles $s'$ that are encoded in map $M$. |
| | information map | $IM(s';\ S, \sigma, \mathbf{s}, M)$<br>$\equiv \sum_s FIG(s, s'; S, \sigma, M) \otimes O^*(s; \mathbf{s})$ | | float, (155,) | [0,1] | $IM(s')$ can be interpret as the average information gain upon observing the neighboring tiles of $s'$ that can be attribute to the indentifiability of $s'$. |
| | information density map | $IDM(s';\ S, \sigma, \mathbf{s}, M) \equiv \frac{IM(s';S,\sigma,\mathbf{s},M)}{n_{\text{edges}}(s';M)}$ | infomap | float, (155,) | [0,1] | $IDM(s')$ is simply $IM(s')$ normalized to the number of encoded edges at $s'$ |
| | information density | $ID(S, \sigma, \mathbf{s}, M) \equiv \frac{\sum_{s'} IM(s';S,\sigma,\mathbf{s},M)}{m}$ | info_mean | float | [0,1] | an overall measure about how informative a given map is |

### 2. Task setup

#### 2.1. Tile walk in a hex-grid world

A simulated agent navigates a hex-grid world consisting of 155 hexagonal tiles, $s \in [0, 154]$, which directly resembles the actual arena a mouse navigates in. In this hex-grid world, only up to six discrete actions $a \in [0, 5]$ are allowed at each step to transition from one tile to an neighbor tile. We import mouse behavior into the hex-grid world by converting its trajectory $\mathbf{x} = (x_0, x_1, \dots), \mathbf{y} = (y_0, y_1, \dots)$ to a tile and action sequence $\mathbf{s} = (s_0, s_1, \dots),\ \mathbf{a} = (a_0, a_1, \dots)$ in two steps:

1) Simply assigning each point $(x, y)$ to the enclosing hexagonal tile $s$.
2) Deriving action from tile transition such that $a_t = s_t \rightarrow s_{t+1}$.

In so doing, we make sure that jagged paths are removed from a raw trajectory (see Pathbundle section in Supplementary Materials for details), and a converted trajectory has no tile repetitions AA..., nor a skipping tile AC... where C is not neighbor of A. Schematically, one step tile walk with a head turn $(s_t, h_t) \rightarrow (s_{t+1}, h_{t+1})$ contains three sub-steps:

1) align heading with action $h_t \rightarrow h'_t = a_t$,
2) walk to the target tile $s_t \xrightarrow{a_t} s_{t+1}$, and
3) rotate to the target heading $h'_t = a_t \rightarrow h_{t+1}$.

The above sub-steps can be operationalized as matrix multiplication on a one-hot pose array $X_t(s, h)$ (or in tensor notation $X_{s^t, h^t}$):

1) roll along heading dimension $X_t \xrightarrow{\text{roll } a_t - h_t \text{ along } h} X'_t$,
2) apply tensor product $\sum_{s^t, a^t} T_{s^t,\, a^t,\, s^{t+1},\, \text{none}}\ X'_{s^t, a^t} \rightarrow X''_{s^{t+1}, a^t}$, and
3) roll along heading dimension $X''_t \xrightarrow{\text{roll } h_{t+1} - a_t \text{ along } h} X_{t+1}$.

A transition $s_i \xrightarrow{a} s_j$ is encoded in the tile transition tensor $T$ where $T_{s_i, a, s_j, \text{none}} = 1$. $T$ is a binary tensor with an entry value 1 encodes a physically allowed transition, and 0 otherwise. Note that the last dimension is a dummy which preserves the heading along action $a$. Note that a nonphysical transition out of the wall could happen during a belief diffusion. For such a transition, we implement a lossy boundary rule to erase the spatial information (more discussion in the later section)

#### 2.2. Self-localization under uncertainty

The task for an agent is to self-localize—i.e., to estimate its currently position—in this hex-grid world under three distinct sources of uncertainty. Specifically, an agent is required to maintain a belief distribution about its current position and heading any moment in time, i.e., $p_t(s, h)$. The task performance is then evaluated based on $p_t(s = s_{\text{ground truth}}, h)$ directly. The three sources of uncertainty are as follows:

1) An agent possesses a noisy actuator and odometer, or simply, a noisy path integrator. With such, a belief distribution will start to diffuse outwards from the ground truth tiles over time without any external sensory inputs.
2) The hex-grid world is inherently ambiguous with identical sensory cues scatter across distant locations.
3) An agent possesses incomplete knowledge of the hex-grid world so that unknown locations cannot be ruled out under any observations.

#### 2.3. Noisy path integrator

Path integrator (PI) fidelity is parameterized as $\kappa \in [0, 100]$ which controls the width of discretized vonmises function from a uniform function $\mathrm{vonm}(h; \kappa = 0) = \frac{1}{6}$ to a one-hot function $\mathrm{vonm}(h; \kappa = 100) = \begin{cases} 1 & \text{if } h = 0 \\ 0 & \text{otherwise} \end{cases}$. A heading diffusion kernel $D$ is constructed with shifted vonmises function $D(h, h';\ \kappa) \equiv \mathrm{vonm}(h - h'; \kappa)$, and diffuses the heading dimension of belief distribution $P$ as follow: $P'_t(s, h) = \sum_{h'} D(h, h';\ \kappa) P_t(s, h')$. We assume that only the heading integration is noisy, and there is no uncertainty in distance integration when an agent traverses from tile to tile.

#### 2.4. Ambiguous tactile perception

Since each tile in the hex-grid world has identical shape, the only informative sensory cue comes from the height difference at the interface (edge) between two tiles, i.e., $\Delta z_{i \rightarrow j} = z_j - z_i$. Note that we choose to encode an edge directionally, i.e., $\Delta z_{i \rightarrow j} \neq \Delta z_{j \rightarrow i}$, for a simpler assumption on what an ideal observer can perceive. Each observation is inherently ambiguous because it does not completely give away the self location, for there are only 9 height differences $\Delta z \in [-4, 4]$, but 930 edges in total. Note that the there are 12 different height levels (each level is 2.5cm higher than the last) in the actual arena, but we clip height differences at level 4 (about 1 body length), i.e., $\Delta z_{i \to j} \equiv \mathrm{clip}(z_j - z_i,\ -4,\ 4)$, to simulate what mouse physically allowed to perceive. The 930 edges are indexed $e \in [0, 929]$ or $(s, h) \in [0, 154] \otimes [0, 5]$, and then stored in a $155 \times 6$ edge feature array: $Z(s, h) = \Delta z_{s \xrightarrow{a=h} s'}$. To convert between 1d and 2d indexes: $(s, h) \equiv \mathrm{divmod}(e, 6)$. Given an observed edge feature $\Delta z$, the ambiguous edges across entire arena is stored in a binary mask:$C_{\text{amb}}(s, h;\ \Delta z) = \begin{cases} 1 & \text{if } Z(s,h) = \Delta z \\ 0 & \text{otherwise} \end{cases}$.

#### 2.5. Limited knowledge of the environment

The limited knowledge an agent possesses about the hex-grid world is quantified as a *sparse map,* $M$, that encodes a fraction of total 930 edges. A sparse map $M = \{e_0, e_2, ...\}$ can range from an empty map: $M_{\text{empty}} = \emptyset$ to a full map $M_{\text{full}} = \{0, 1, ...929\}$. Given a spare map $M$, a binary mask, $C_{\text{unk}}(s, h;\ M) = \begin{cases} 1 & \text{if an edge } (s,h) \notin M \\ 0 & \text{otherwise} \end{cases}$, can be constructed to mark the unknown edges. The more unknown edges there are, the less likely an agent can localize itself under an arbitrary observation, for any unknown edges are inherently ambiguous by default to any observations.

### 3. Task evaluation

#### 3.1. Ideal observer

We assume an agent to be an ideal observer when inferring its own location. Note that the reason for using an ideal observer is not to model the mouse brain, but to compute a performance upper bound given all the environmental and self-motion cues an agent can gather. The three sources of uncertainty discussed above defines the agent's two-step internal belief update, i.e., diffusion step and collapse step, which we will unpack next.

#### 3.2. Diffusion of belief distribution

In the diffusion step, the uncertainty of self position increases due to the self motion is inherently noisy. As discussed in the last section, we assume a noise model for moving in the hex-grid world where only the heading variable $h$ is noisy whereas the traversing distance is deterministic at every step. To diffuse the joint belief distribution of tile position and heading by one step, i.e., $P_t(s, h) \xrightarrow{\text{diffuse}} \tilde{P}_{t+1}(s, h)$, we first apply diffusion kernel along the heading dimension: $\sum_{h'} D(h, h';\ \kappa) P_t(s, h') \to P'_t(s, h)$. Next, we apply a transformation, identical to the three-step operation for tile walk discussed above, on $\tilde{P}_t(s, h)$:

1) roll along heading dimension towards the upcoming action $P'_t \xrightarrow{\text{roll } a_t - h_t \text{ along } h} P''_t$,

2) apply tensor product for transitioning to the next tile $\sum_{s^t, a^t} T_{s^t, a^t, s^{t+1}, \text{none}}\ P''_{s^t, a^t} \to P'''_{s^{t+1}, a^t}$, and

3) roll along heading dimension $P'''_t \xrightarrow{\text{roll } h_{t+1} - a_t \text{ along } h} \tilde{P}_{t+1}$.

Below, we show two examples of belief diffusion while an agent is walking straight without any knowledge about the hex-grid world:

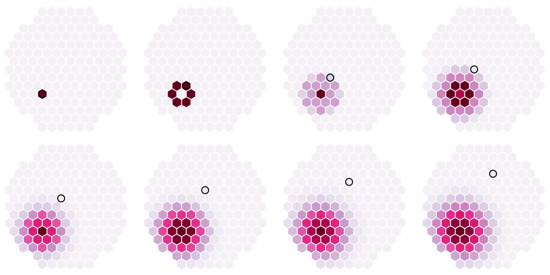

**Figure 1:** low PI fidelity: $\kappa = 0$

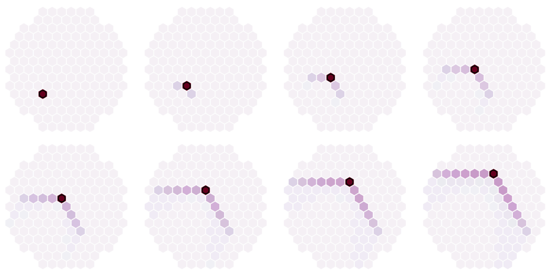

**Figure 2:** high PI fidelity: $\kappa = 80$

#### 3.3. Collapse of belief distribution

In the collapse step, $\tilde{P}_{t+1}(s,h) \xrightarrow{\text{collapse}} P_{t+1}(s,h)$, the uncertainty of self position either decreases or stays the same depending on the current sensory cues $Z(s_{t+1},h)$ and the agent's knowledge $M$ about the arena. Upon arriving a tile $s_{t+1}$, we assume the agent observe all 6 edges of the tile before going to the next one. The belief update under such a series of observations follows standard Bayes rule: $P_{t+1}(s,h) = \dfrac{C_{\text{obs'd at } s_{t+1}}(s,h)\ \tilde{P}_{t+1}(s,h)}{\sum_{s,h} C_{\text{obs'd at } s_{t+1}}(s,h)\ \tilde{P}_{t+1}(s,h)}$ where the total mask:

$$C_{\text{obs'd at } s_{t+1}} = \prod_{h=0}^{5} C_{\text{unk}}(s,h;\ M) + C_{\text{amb}}(s,h;\ Z(s_{t+1},h)).$$

#### 3.4. Belief distribution propagation

To sum up, one step belief distribution propagation from $s_t$ to $s_{t+1}$ is simply the concatenation of the diffuse and collapse steps:

$$P_t(s,h) \xrightarrow{\text{diffuse}} \tilde{P}_{t+1}(s,h) \xrightarrow{\text{collapse}} P_{t+1}(s,h)$$

#### 3.5. Lossy boundary rule

To generate a tile walk trajectory, we either convert a mouse trajectory, or sample one from a self-avoiding random walker (discussed below). In either case, there are no unphysical transitions out of the arena at a tile next to the boundary; i.e., at a wall tile, $s_{\text{wall}} \xrightarrow{a_{\text{unphy}}} s \notin [0,154]$ where $a_{\text{unphy}}$ never occurs. However, to propagate a belief distribution, one faces this issue of unphysical transitions in which some belief value could leak out of arena after a diffusion step causing $\sum_{s,h} \tilde{P}_t(s,h) < 1$, and thus require an additional renormalization step. Consequently, this belief renormalization step effectively provides the agent sensory cues about where the wall is by ruling out those vanishing belief values at certain wall tiles. Among various solutions to this issue, we choose to implement a lossy boundary rule that effectively erase any position information upon a unphysical transition: $T_{s^t,a^t,s^{t+1},\text{none}} = \begin{cases} 1 & \text{if } a^t = a_{\text{phy}} \\ 1\,/\,155 & \text{if } a^t = a_{\text{unphy}} \\ 0 & \text{otherwise} \end{cases}$ which redistributing the out-of-boundary belief values back to all tiles uniformly.

#### 3.6. Localization score

We define a score map $S$ in which a tile score for $s$ is the mean of the marginal belief value when the agent is physically on it:

$$S(s;\mathbf{s},M) \equiv \frac{1}{\sum_t w_t} \sum_t \left( w_t \sum_{h^t} P_{s^t,h^t} \right), \text{ where } w_t = \begin{cases} 1 & \text{if } s^t = s \\ 0 & \text{otherwise} \end{cases}.$$

Note that for a certain tile $s_{\text{unoccupied}}$ that an agent never visit, $S(s_{\text{unoccupied}})$ is not defined. These holes in a score map make the overall evaluation tricky. Ideally, one should not include those holes in the final score for the agent does not even utilize those associate sensory cues to navigate the arena. However, consider the following comparison between two agents:

- Agent #1 covering all 155 tiles uniformly, it would need a large map to reach a high performance overall, whereas
- Agent #2 covering only a small fraction of arena only needs a much smaller map to ensure high performance on those occupied tiles.

To balance these extremes we define the localization score $\overline{S}$ as the mean of top half (77 tiles) of highly occupied tiles only:

$$\overline{S} = 2 \sum_s S(s;\mathbf{s},M) O^*(s;\mathbf{s}),$$

where $O_b(s)$ is binarized occupancy map $O(s)$ s.t. $O^*(s;\mathbf{s}) = \begin{cases} 1 & \text{if } O(s) > \text{median}(O) \\ 0 & \text{otherwise} \end{cases}$. This definition punish a highly sparse trajectory because there will be many low score ones among the chosen 77 tiles.

### 4. Sparse map optimization

#### 4.1. Defining the optimization problem

To optimize a sparse map, we first fix the following three parameters:

1) a tile walk trajectory $\mathbf{s} = (s_0, s_1, ..., s_t, ...)$ and $\mathbf{a} = (a_0, a_1, ..., a_t, ...)$ with $t_{max} \sim 20,000$.

2) a path integrator fidelity $\kappa \in [0, 100]$

3) a fixed sparsity, or equivalently, a map size $m \in [0, 930]$

The optimization problem can then defined as

$$M^* = \underset{M|m}{\operatorname{argmax}} \ \overline{S}(\mathbf{s}, \mathbf{a}, \kappa, M),$$

where $\overline{S}$ is the localization score defined in the last section. Given the vast space of possible map with a moderate map size: $n_{M|m} = C_m^{930}$, it is not feasible to evaluate them all to find the optimal. Below, we introduce a greedy algorithm to greatly narrow the search space while acquiring a performance much greater than a randomly sampled map, i.e., $\overline{S}\big(\mathbf{s}, \mathbf{a}, \kappa, M^*\big) \gg \overline{S}(\mathbf{s}, \mathbf{a}, \kappa, M_{\text{rand}})$.

#### 4.2. Iteratively deleting the least important edge

The greedy optimizer starts from the full map $M_{\text{full}}$ and progressively deletes one least important remaining edge at each iteration. When there is a tie in score among multiple candidates, we select the least visited edge to delete. As a result, one acquires a series of optimized sparse maps $\Big(M_{\text{full}}, M^*_{m=929}, ..., M^*_{m=1}, M_{\text{empty}}\Big)$ with every map size. Even with such a reduction of search space, it is still infeasible to run with our compute budgets, we thereby adopt three implementations to speed up the optimizer without significantly compromising its performance.

#### 4.3. Three implementations to speed up the optimization

##### A. Normalizing just once for computing posterior

As discussed in the collapse step above, the agent could observe up to six edges at each tile, and thereby up to six sequential computation for updating posterior. It is much faster to compute the total mask as the product of individual masks, and treat the original six observations as a singular computation for updating posterior.

##### B. Rolling $T$ instead of rolling $P$ for head turns

Since we diffuse heading belief the same amount at each step (independent of the actual head turning) during a tile walk, the head turn $h^t = a_t \rightarrow h^{t+1} = a_{t+1}$ bares no consequences in how belief distribution actually propagates. Knowing this, one can pre-compute up to just six heading updates and pack them in the tile transition tensor instead of computing ~20,000 actual heading updates: $T_{\text{old}} \rightarrow T_{\text{new}} = T_{h\in[0,5]}$.

##### C. Evaluating on a small chunk of the full trajectory with an annealing-like scheduler

To further speed up the optimization, we split the full trajectory into $n_c = L/l_c$ chunks where $L$ is the trajectory length and $l_c$ is the chunk length. At iteration $i$, the $j$-th chunk $c_j = (s_{jl_c}, s_{jl_c+1}, ..., s_{(j+1)l_c-1})$ is evaluated, where $j = i \bmod n_c$. This speeds up the overall optimization roughly $n_c \sim 20$ fold. To ensure such a sped-up optimization still resembles the original one when evaluating on the full trajectory, we require $n_c \ll 930$ (total number of iterations) so that each part of the full trajectories has plenty of coverage.

Next, we progressively increases $l_c$ over the course of optimization via a scheduler: $(c_j,\ l_c,\ n_c)_{\text{iter } i}$ in which the computation cost is kept the same: $l_{c\ |\ \text{iter } i} \times (930 - i) \approx$ const. The scheduler is design to use shorter chucks in the beginning because the optimizer can afford to be a bit sloppy then. In earlier iterations when the map size is big and the performance is high, many candidate deletions yield the same result, doing a through evaluation could be wasteful. However, at later iterations, when the performance undergoes a rapid collapse, it is essential to select the next edge to delete very carefully by evaluating a much longer chunk.

#### 4.4. Pre-filtering candidate edges basing on tile occupancy

There is a caveat after implementing the chunk evaluation due to the low arena coverage in the earlier iterations. Given that a short chunk is about 1/30 of the full trajectory, there could be many more tiles that are not visited within each chuck. Consequently, there would be no performance changes by deleting the edges associate with those unoccupied tiles. However, these premature deletions could greatly affect the performance when evaluating on the full trajectory. To avoid this issue, we remove those unvisited edges from the candidate list at each iteration. Note that for those edges that are never visited on the full trajectory, we always keep them on the candidate list at every iteration. The final candidates at each iteration is summarized in the diagram below:

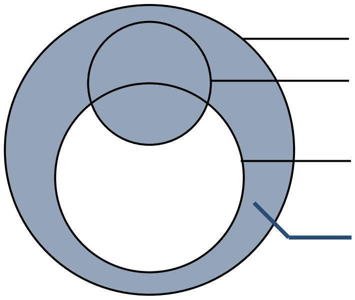

**Drawing 1: Compute candidate edges at iteration $i$**

### 5. Numerical experiments

For the map optimization introduced above, one finished job consists of 931 optimized maps, from a full map to an empty map, each is the same as the last but one edge less. In this study, we run three large job batches with each job being labeled with multiple parameters as discussed next.

#### 5.1. Parameter space

A job is parameterized with 1) map type, 2) trajectory type, 4) trajectory id, 3) path integrator (PI) fidelity. We organize the jobs into three batches based on map type:

| map_type | note |
|---|---|
| optmap | optimized map |
| randmap | random map |
| wallmap | fixed wall-only map with 90 edges encoded |

#### 5.2. Job batch for optimized maps

Within this batch, a job is labeled with the remaining parameters:

```
job = ((traj_type, traj_id, traj_subid), pi_level)
```

where each parameter can take on multiple values:

| traj_type | traj_id | traj_subid | note |
|---|---|---|---|
| mouse | mouse_id ∈ [0,6] | none | |
| mouse_rotated | mouse_id ∈ [0,6] | rot_id ∈ [1,5] | $\theta_{rot} = \texttt{rot\_id} \times 60°$ |
| random_walk | traj_seed ∈ [0,9] | none | |

&

$\texttt{pi\_level} \in \{0,10,20,...90\}$

These parameter combinations amount to *520 jobs.*

#### 5.3. Job batch for random maps

For random maps, an additional `map_seed` parameter is added:

```
job = ((traj_type, traj_id, traj_subid), pi_level, map_seed)
```

where each parameter can take on multiple values:

| traj_type | traj_id | traj_subid | map_seed | note |
|---|---|---|---|---|
| mouse | mouse_id ∈ [0,7] | none | [0,9] | 10 random maps. Each map seed yield a shuffled list of 930 edges. A map with map size $m$ is a trimmed sub-list up to the $m$-th entry. |
| random_walk | traj_seed ∈ [0,9] | none | [0,9] | sharing the same set of 10 random maps above. |

&

pi_level ∈ {0,10,20,...90}

These parameter combinations amount to *1700 jobs.*

#### 5.4. Job batch for wall-only maps

Unlike the job batches above, there is only one wall-only map. We thereby replace the variable within a job from map size to PI fidelity: pi_level ∈ {80,81,...,90}. The parameter to specify a job is then

```
job = (traj_type, traj_id, traj_subid)
```

with each job parameter take on the values same as the batch `'optmap'`:

| traj_type | traj_id | traj_subid |
|---|---|---|
| mouse | mouse_id ∈ [0,7] | none |
| mouse_rotated | mouse_id ∈ [0,7] | rot_id ∈ [1,5] |
| random_walk | traj_seed ∈ [0,9] | none |

These parameter combinations amount to *52 jobs*.

### 6. Results

In this section, we show the complete results of the above numerical experiments:

1) the raw performance curves as a function of map size,
2) the heat maps of performance landscape,
3) the tradeoff curves of a constant target score on performance landscape, and
4) the compression rates as a ratio of performance gain from a random map.

#### 6.1. Optimized maps achieve higher localization score than random maps

In the figure below, each performance curve as a function of map size ∈ [0,930] is one complete map optimization given a set of fixed parameters:

1) map type ∈ {optimized map, random map}
2) trajectory type ∈ {mouse, rotated mouse, random walk}
3) path integrator (PI) fidelity
4) trajectory id ∈ {mouse id, seed id for random walk, seed id for random map}

These curves are grouped by the first three parameters. The multiple trajectory ids within a group are treated as the source of performance variation. From these results, one can see that

1) the optimized maps (labeled *optmap*) consistently perform better than random maps (labeled *randmap*)
2) the maps optimized on a mouse trajectory (labeled *optmap, mouse*) have smaller performance variation than any other groups.

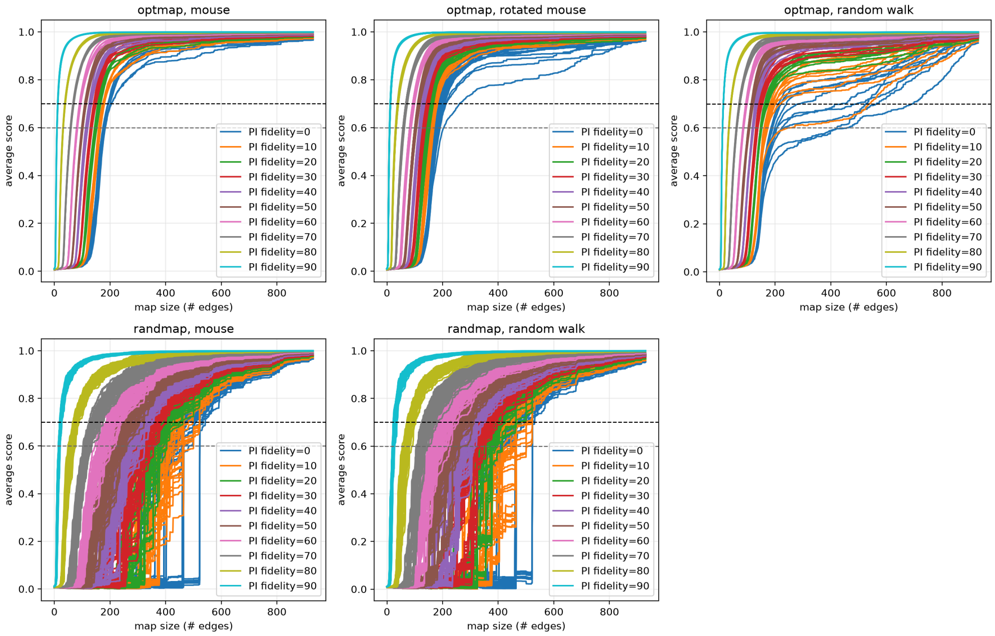

**Figure 3:** Raw performance curves as a function of map size

#### 6.2. Optimized maps are 2x more efficient than random maps

Next, we plot the performance landscape (as a function of map size and PI fidelity) by averaging over the trajectory ids within a group defined above. For each performance landscape (heat map), we plot a tradeoff curve via collecting the map sizes $m^{\text{(target)}}$ that achieve the same localization score closest to a fixed target score: i.e., $m^{\text{(target)}}$ s.t. $\overline{S}\left(m^{\text{(target)}}; \mathbf{s}, M\right) = \overline{S}^{\text{(target)}}$. The variations of $m^{\text{(target)}}$ directly come from the performance variations due to multiple trajectory ids within a group; in the figure below, we plot 95% confidence intervals (CI) as error bars on the tradeoff curve for each heat map. All tradeoff curves exhibit a similar decreasing trend in map size while increasing PI fidelity. And it's directly apparent that the tradeoff curves for optimized maps are shallower (indicating better efficiency) than that of random maps. To compare tradeoff curves among optimized maps, we plot the compression rate as "percentage increase in map size" in the main text which shows that

> *the maps optimized on a mouse trajectory achieve significantly higher efficiency than that on a rotated mouse trajectory or a random walk trajectory*.

In the last panel of the figure below, we instead plot compression rates $CR$ that are normalized to the tradeoff curves of random maps:

$$CR_x \equiv \frac{\overline{m}^{\text{(target)}}_{\text{baseline}}}{\overline{m}^{\text{(target)}}_x},$$

and the error bar calculated with uncertainty propagation:

$$\Delta CR_x = \overline{CR}_x \times \left( \frac{\Delta m^{\text{(target)}}_x}{\overline{m}^{\text{(target)}}_x} + \frac{\Delta m^{\text{(target)}}_{\text{baseline}}}{\overline{m}^{\text{(target)}}_{\text{baseline}}} \right)$$

With a target score $\overline{S}^{\text{(target)}} = 0.7$, one reach similar conclusions in the main text:

1) *the optimized maps are roughly 2x more efficient (with a higher compression rate) than random maps;* and
2) *the maps optimized on a mouse trajectory are significantly more efficient, across a wide range of PI fidelity, than that on a rotated mouse trajectory or on a random walk trajectory.*

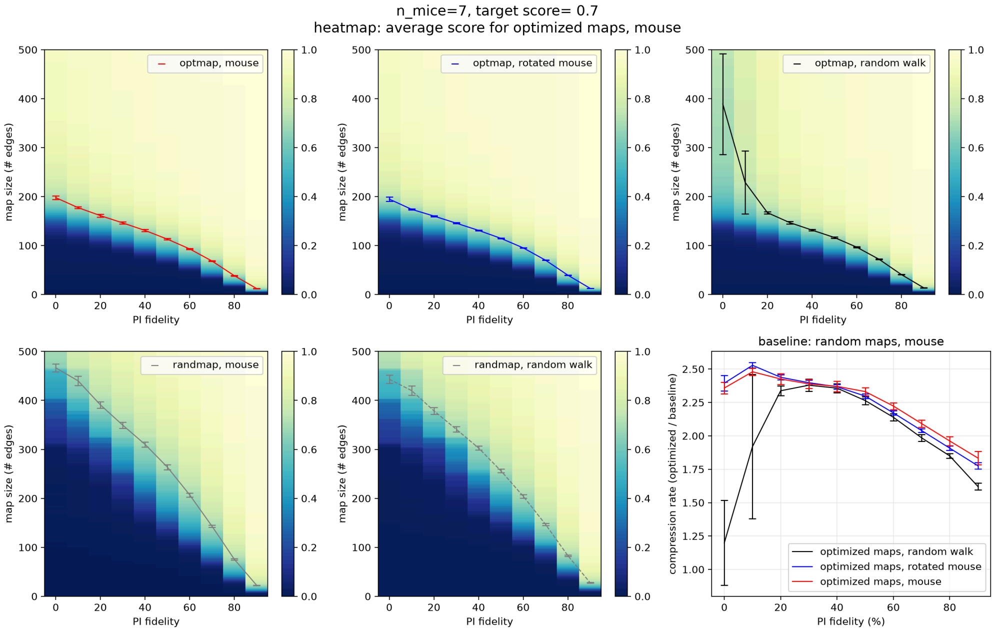

**Figure 4:** Heat maps of performance landscape. Last panel: compression rate as normalized tradeoff curves of a constant target score on performance landscape

### 7. Information density correlates with localization score

The main results from our numerical experiments (above and in the main text) show that the maps optimized on mouse trajectories are better than the other baseline maps in performing the location task, i.e.,

$$\overline{S}_{\text{optmap, mouse}} \gtrsim \overline{S}_{\text{optmap, rot mouse}} \gtrsim \overline{S}_{\text{optmap, random walk}} > \overline{S}_{\text{randmap}}$$

given the same map size $m$. To explain why optimized mouse maps perform better, we hypothesize that they achieve higher *information density* overall by selectively encoding more locally informative edges.

#### 7.1. Information density defined as regional indentifiability

Intuitively, information density should capture identifiability of an agent's self location given

1) its knowledge of the environment (sparse map),
2) prior belief (a position distribution), and
3) immediate observation (edge features of the current tile).

Since we focus on the good performance regime, we assume the prior belief is highly localized at ground truth location, i.e., belief diffusion radius $\sigma(s) \sim 1\,/\,S(s)$ where $S(s)$ is the location score at tile $s$. And thus information density should be local, and could be readily interpreted as how much observed features $\{Z(s,h)\ \forall h\}$ help to discriminate tile $s$ among the neighbor tiles within radius $\sigma$.

##### A. Locality

Formally, we define tile locality as a gaussian envelope:

$$L(s, s'; \sigma) \equiv \frac{e^{-|\mathbf{r}_s - \mathbf{r}_{s'}|^2/(2\sigma(s)^2)}}{\sum_{s'} e^{-|\mathbf{r}_s - \mathbf{r}_{s'}|^2/(2\sigma(s)^2)}},$$

where $|\mathbf{r}_s - \mathbf{r}_{s'}|$ is the distance (normalized to tile width) between tile $s$ and $s'$, and $\sigma(s)$ can be computed by numerically solving the following equation which ensures the peak of gaussian at ground truth tile $s$ to match the localization score $L(s, s' = s; \sigma(s)) = S(s)$:

$$f(\sigma; S) \equiv \sum_{s'} e^{-(\mathbf{r}_s - \mathbf{r}_{s'})^2/(2\sigma^2)} - \frac{1}{S(s; \mathbf{s}, M)} = 0$$

**B. Ambiguity**

To quantify how informative a given observation on tile $s$ against the agent's knowledge of the arena—sparse map $M$, we define an auxiliary function—i.e., local tile ambiguity between two tiles $(s, s')$ is a binary similarity measure $A$ weighted by locality $L$:

$$A_{\text{local}}(s, s'; \sigma, M) \equiv \frac{L(s, s'; \sigma)A(s, s'; M)}{\sum_{s'} L(s, s'; \sigma)A(s, s'; M)},$$

where the binary tile ambiguity is defined as

$$A(s, s'; M) \equiv \begin{cases} 1 & \text{if } D_Z(s, s'; M) = 0 \\ 0 & \text{if } D_Z(s, s'; M) > 0 \end{cases}$$

with the feature distance $D_z$ defined as Hamming distance between an observed feature vector $Z(s, h)$ at tile $s$ and a masked or "remembered" feature vector $Z_M(s', h) = \begin{cases} Z(s', h) & \text{if } (s', h) \in M \\ \text{nan} & \text{if } (s', h) \notin M \end{cases}$ at tile $s'$ based on agent's sparse map. The distance is computed as the minimum of the six possible rotations

$$D_Z(s, s'; M) \equiv \min_{\theta} \sum_{\substack{h \text{ s.t.} \\ Z_M(s', h) \neq \text{nan}}} |\text{roll}(Z(s, h), \theta) - Z_M(s', h)|$$

With this definition, even if two tiles are inherently unambiguous—i.e., $Z(s, h)$ is a dissimilar vector from $Z(s', h)$, two tile can still be ambiguous if the agent does not have the knowledge to tell them apart; e.g., for an empty map $M_\emptyset$, $Z_M(s', h) = \text{nan}\ \forall h$, and $D_Z(s, s'; M_\emptyset) = 0$.

Note that local tile ambiguity can be interpreted as a belief posterior after observing tile $s$ if the prior distribution is tile locality function $L$; that is,

$$P_{\text{before}}(s') = L(s, s'; \sigma) \xrightarrow{\text{upon observing } s} P_{\text{after}}(s') = A_{\text{local}}(s, s'; \sigma, M)$$

**C. Identifiability**

Local tile identifiability is simply a complementary measure to local tile ambiguity:

$$I_{\text{local}}(s, s'; \sigma, M) \equiv \frac{L(s, s'; \sigma)I(s, s'; M)}{\sum_{s'} L(s, s'; \sigma)I(s, s'; M)},$$

where $I(s, s'; M) \equiv 1 - A(s, s'; M)$. This additional measure is needed to attribute information gain to the encoded tiles in a sparse map as we will discuss next.

**D. Fractional information gain**

In Bayesian inference, the information gain after an observation can be quantified as the distance between the prior and posterior belief distributions. We thus choose KL-divergence to quantify the information gain after an observation at tile $s$:

$$\begin{aligned} IG(s; \sigma, M) &\equiv D_{KL}(P_{\text{after}} \parallel P_{\text{before}}) \\ &= D_{KL}(A_{\text{local}} \parallel L) \\ &= -\log \sum_{s'} L(s, s'; \sigma)A(s, s'; M) \end{aligned}$$

To derive the last equality, we use the fact that $A_{\text{local}}$ is simply a masked version of $L$. To better intuit this quantity, we normalize it to the maximum gain—which happens when the agent is fully localized upon observing $s$, $L(s, s'; \sigma) \xrightarrow{\text{upon observing } s} \delta(s, s')$:

$$IG_{\text{norm}}(s; S, \sigma, M) \equiv \frac{IG(s; \sigma, M)}{IG_{\text{max}}(s; S)} \in [0, 1]$$

where $IG_{\text{max}}(s; S) = D_{KL}(\delta(s, s') \parallel L) = -\log S(s; \mathbf{s}, M)$.

Next, we define fractional information gain $FIG(s, s')$ to attribute $IG_{\text{norm}}(s)$ at $s$ to the neighbor tiles $s'$ using local tile identifiability $I_{\text{local}}(s, s')$:

$$FIG(s, s'; S, \sigma, M) \equiv I_{\text{local}}(s, s'; \sigma, M) \otimes IG_{\text{norm}}(s; S, \sigma, M)$$

**E. Information density**

With fractional information gain in hand, it is easy to define an information density map $IDM(s')$ by summing over all observed tiles $s$ with nonzero contribution to the encoded tile $s'$:

$$IDM(s'; S, \sigma, \mathbf{s}, M) \equiv \frac{\sum_s FIG(s, s'; S, \sigma, M) \otimes O^*(s; \mathbf{s})}{n_{\text{edges}}(s'; M)}$$

where $n_{\text{edges}}(s'; M)$ is number of encoded edges on tile $s'$ according to the sparse map $M$, and $O^*(s; \mathbf{s}) = \begin{cases} 1 & \text{if } O(s; \mathbf{s}) > \text{median}(O) \\ 0 & \text{otherwise} \end{cases}$ is a binary weight to specify the highly occupied tiles under the agent's trajectory $\mathbf{s}$. The main idea is to only consider observations that are frequently made by the agent. The overall information density $ID$ can then computed as the average over all encoded edges:

$$ID(S, \sigma, \mathbf{s}, M) \equiv \frac{\sum_{s'} IM(s'; S, \sigma, \mathbf{s}, M)}{m}$$

where $m$ is the total number of edges encoded in map $M$. This definition of information density depends on both the trajectory $\mathbf{s}$ and map $M$, which might not be ideal for our purpose—i.e.,

*To explain that an optimized map achieving higher performance for it encodes highly informative edges among neighbors*.

One could argue, in an extreme scenario, that information density depends solely on the trajectory $\mathbf{s}$ and has nothing to do with the map —i.e., $ID(\mathbf{s})$ instead of $ID(\mathbf{s}, M)$.That is, the higher localization score $\overline{S}(M_{\text{opt}}) > \overline{S}(M_{\text{rand}})$ resulting from a better map $M_{\text{opt}}$ cannot be explained by a measure that is independent of the map. Ideally, one would like to have the opposite scenario—i.e., $ID(M)$ to remove the confound $\mathbf{s}$. In the discussion below, we show how the information density depends mostly on the map as long as the performance is high, or in other words, the belief distribution is highly localized near the ground truth tile at all time.

### 7.2. A simple interpretation of information density

We argue that the information density defined above depends mostly on the agent's map instead of detail trajectory as long as its performance is high. The intuition behind this assessment is the following:

*The high performance ensures that the information gain upon an observation at tile $s$ mostly depends only on its nearest neighbors $s' \in \left\{ s_{\text{nbr}} \text{ s.t. } s \xrightarrow{a} s_{\text{nbr}} \ \forall a \right\}$ which is independent of the detail trajectory $\mathbf{s}$.*

For a gaussian belief distribution, the diffusion radius $\sigma$ (standard deviation) is already less than one tile width for a mid-level score $\overline{S} \approx 0.4$, and thus it's safe to say that nearly all contribution to the information gain depends on the nearest neighbor tiles alone for our target score $\overline{S}^{\text{(target)}} = 0.7$. Note that this independence of detail trajectory does not hold for low performance regime where the belief distribution cannot be approximated as a localized gaussian envelope and the diffusion radius $\sigma$ could change from time to time as well. Below, we re-derive a simplified information density by taking a limit $\overline{S} \to 1$:

$$\begin{aligned} IG_{\text{norm}}(s;S,\sigma,M) &\equiv \frac{IG(s;\sigma,M)}{IG_{\text{max}}(s;S)} \\ &= \frac{-\log \sum_{s'} L(s,s';\sigma)A(s,s';M)}{-\log S(s;\mathbf{s},M)} \\ &\xrightarrow{S=1-\epsilon} \frac{\log 1 - n_{\text{identifiable}} \times \frac{\epsilon}{6}}{\log 1-\epsilon} \\ &\approx \frac{n_{\text{identifiable}} \times \frac{\epsilon}{6}}{\epsilon} = \frac{n_{\text{identifiable}}(s;M)}{6} \end{aligned}$$

$$\Longrightarrow FIG(s,s';S,\sigma,M) = \begin{cases} 1\,/\,6 & \text{for identifiable tile } s' \\ 0 & \text{otherwise} \end{cases}$$

where $n_{\text{identifiable}}$ is number of identifiable tiles among the six nearest neighbor tiles of $s$. The information density map can then be simplified as

$$\begin{aligned} IDM(s';M) &= \frac{\sum_{\text{identifiable } s\in\text{nbrs of } s'} O^*(s;\mathbf{s}) \times \frac{1}{6}}{n_{\text{edges}}(s';M)} \\ &= \frac{n_{\text{identifiable \& freq-visited nbrs}}(s';M)}{6\ n_{\text{edges}}(s';M)} \end{aligned}$$

And the overall information density:

$$ID(M) = \frac{\sum_{\text{encoded } s'} n_{\text{identifiable \& freq-visited nbrs}}(s';M)}{6\ m},$$

which is independent of the detail trajectory $\mathbf{s}$, but only depends on the map $M$ and the coarse-grained occupancy $O^*(s;\mathbf{s})$. This simplified version of information density at an encoded tile $s'$ can be intuitively interpreted as *the fraction of identifiable and frequently observed tiles $s$ out of the six nearest neighbors of $s'$* as illustrated below:

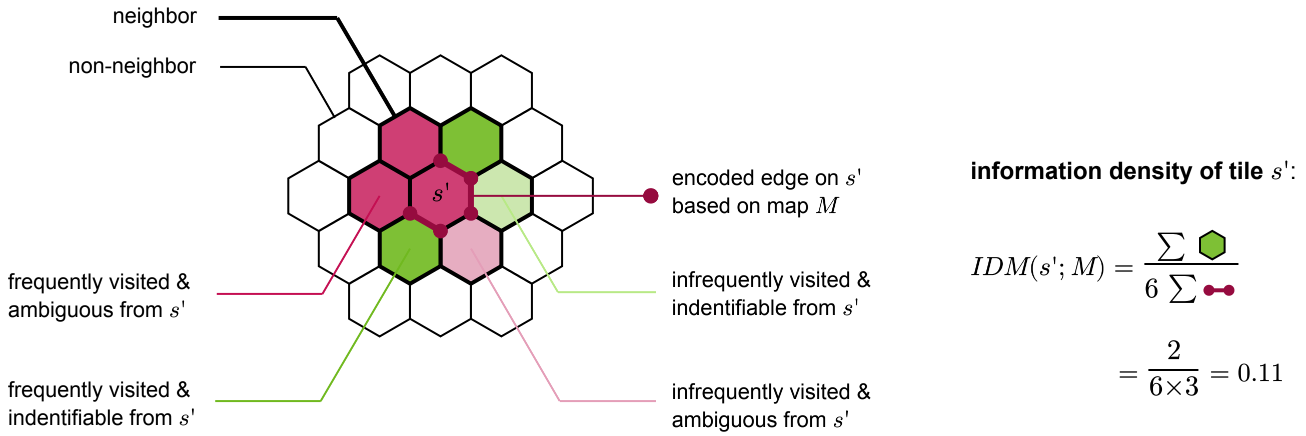

**Drawing 2:** Compute simplified information density of tile $s'$ under the nearest neighbor limit.

#### 7.3. Optimized maps encode more informative edges than random maps

In this subsection, we show that both the original and simplified versions of information density defined above yield qualitatively similar results. We compute them across different map optimizations, trajectories, map sizes, and path-integration (PI) fidelities. The goal is to explain, with information density, the two main findings about the localization performance in the main text:

1) Optimized maps perform better than random maps: $\overline{S}_{\text{optmap}} > \overline{S}_{\text{randmap}}$.

2) A map optimized on a mouse trajectory is more efficient than that optimized on a random walk trajectory: $\overline{S}_{\text{optmap, mouse}} \gtrsim \overline{S}_{\text{optmap, random walk}}$.

### A. Original version of information density

We organize the results into the similar heat maps shown in the main text, but with the information density presented instead of the localization score. The five heat maps are labeled with map type (*optmap or randmap*) and trajectory type (*mouse, rotated mouse, or random walk*). Each pixel in a heat map is a specific combination of map size and PI fidelity.

To highlight the high performance regime, we select one map size at one PI fidelity that yield $\overline{S}_{\text{optmap, mouse}} \approx \overline{S}_{\text{target}} = 0.7$. Note that these map sizes are selected from the maps optimized on the mouse trajectories:

| PI fidelity (%) | selected map size | PI fidelity (%) | selected map size |
|---|---|---|---|
| 0 | 198 | 50 | 113 |
| 10 | 177 | 60 | 93 |
| 20 | 161 | 70 | 68 |
| 30 | 146 | 80 | 39 |
| 40 | 131 | 90 | 12 |

To have a fair comparison, we fix these map sizes across different heat maps shown in the figure below as the same write curves in the heat map. One can see that the three heat maps of optimized maps (top row) are qualitatively the same, and very different from that of the random maps (bottom rows). In the last summary plot, at the selected map sizes, one can see that the information density are significantly different between optimized maps and random maps. This clear our first goal to explain a better (first-order) localization performance with a higher information density. However, for the subtler (second-order) performance difference among different optimized maps, the information density is an insufficient explainer: i.e., $ID_{\text{optmap, mouse}}$ is overlapping with $ID_{\text{optmap, random walk}}$ across most of the PI fidelities.

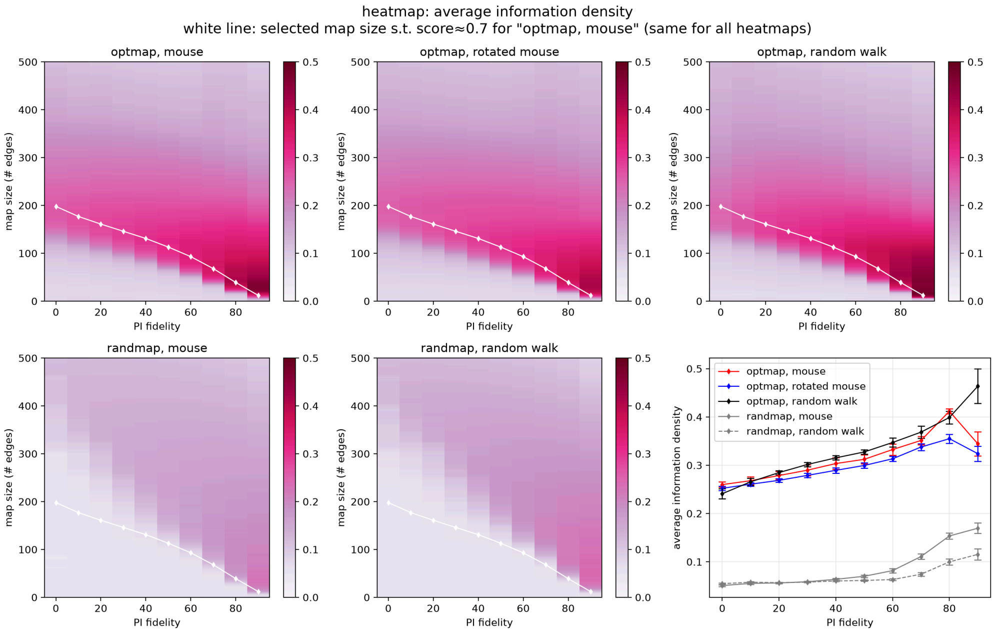

**Figure 5:** Heat maps of information landscape; white curves: selected map size so that $\overline{S}_{\text{optmap, mouse}} \approx 0.7$. Last panel: average information density at selected map size for each PI fidelity

To see how optimized maps encode informative edges, we plot information density maps for PI fidelity=70% and map size=68 edges in the figure below. One can see that 1) the information density (colors) on most tiles of an optimized map is larger than that of a random map, and 2) an optimized map encodes just one edge per tile (small circle size) at such a high sparsity unlike a random maps (encodes upto 3 edges per tile) which shows that the map optimizer allocates limited resource more effectively.

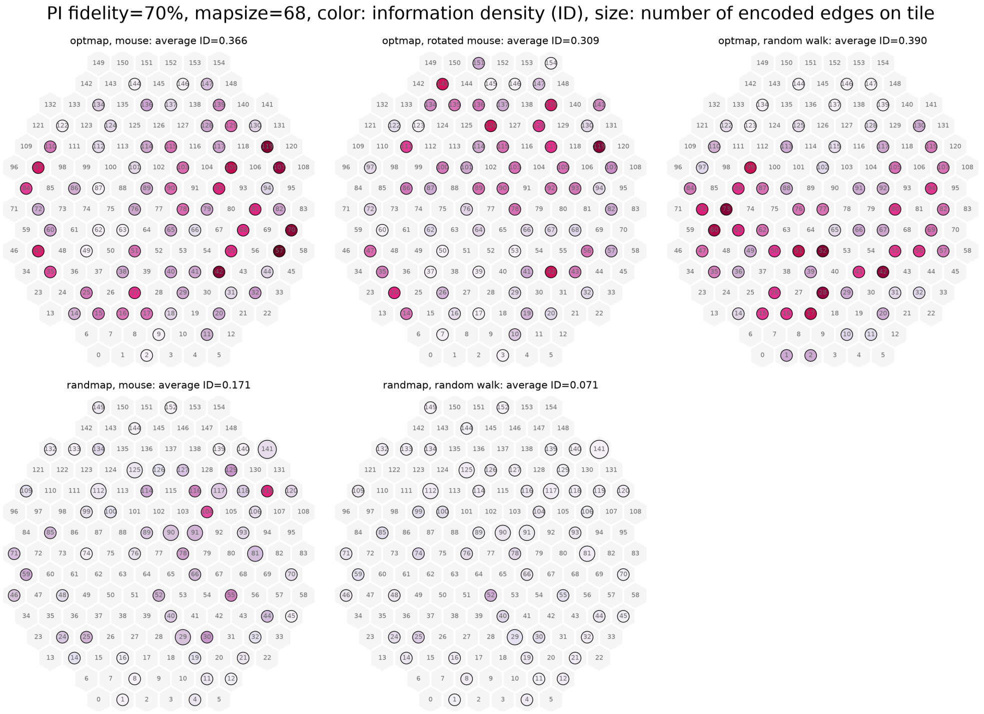

**Figure 6:** Information density maps at PI fidelity=70% and map size=68 edges

#### B. Simplified version of information density

As discussed earlier, the original information density is defined in such a way that could implicitly depend solely on the detailed trajectory $\mathbf{s}$. To resolve this potential confound—i.e., the better performance due to an optimized map cannot be explained by a measure that is independent of the map $M$, we derive a simplified version of information density by taking the high-performance limit: $\overline{S} \to 1$ so that $IDM(s'; M) = \frac{n_{\text{identifiable \& freq-visited nbrs}}(s';M)}{6\ n_{\text{edges}}(s';M)}$ depends only on the map.

The figures below are in identical format to the figures in the last section except now the new version of information density is plotted. From the summary plot at bottom right panel, one can see that the trends of these curves are qualitatively the same comparing to the original version; that is, 1) the optimized maps have higher information density than the random maps: $ID_{\text{optmap}} > ID_{\text{randmap}}$, and 2) within optimized maps $ID_{\text{optmap, mouse}}$ is overlapping with $ID_{\text{optmap, random walk}}$ across most of the PI fidelities.

Comparing the heat maps below to the origional version, one can see that the main difference show up in the area below the white curves where the high-performance limit fails. This difference is most apparent for the random maps because their performance is much lower on the white curves than the optimized maps—e.g., $\overline{S}_{\text{optmap, mouse}} \approx 0.7$ whereas $\overline{S}_{\text{randmap, mouse}} \approx 0.2$. As a result, their information density maps in the figure below also show the most differences.

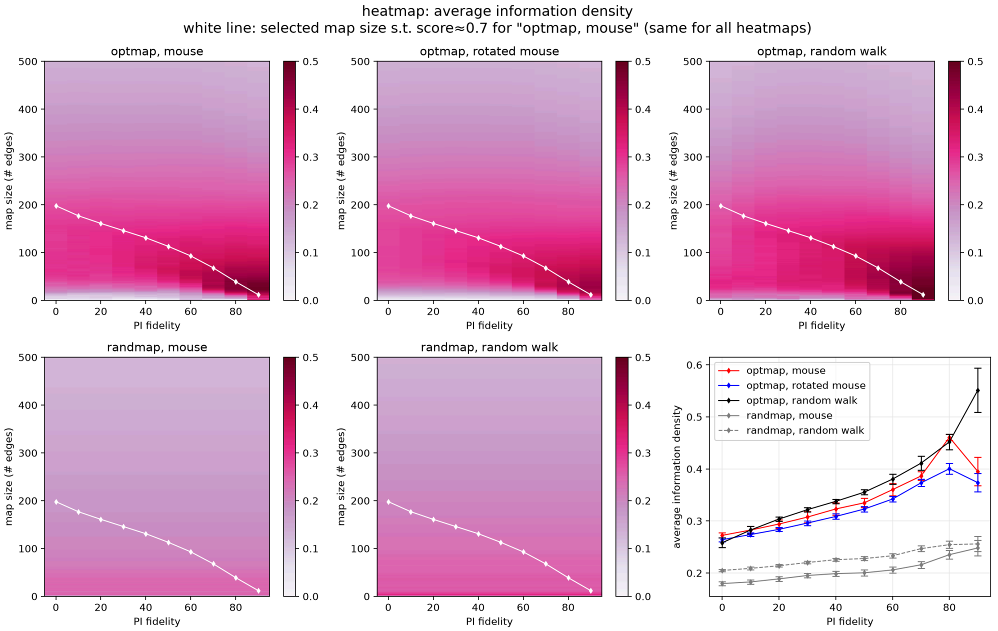

**Figure 7:** Heat maps of simplified information landscape. All parameters are the same as Figure 5.

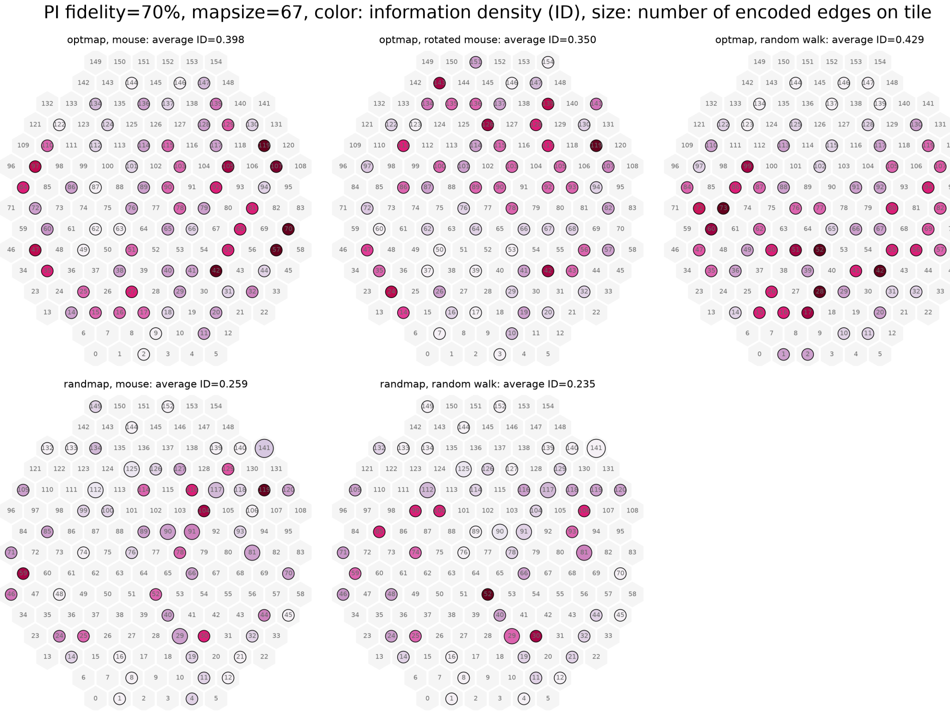

**Figure 8:** Simplified information density maps. All parameters are the same as Figure 6.

#### 7.4. Information density is insufficient to explain how an optimized map is better than the others

As discussed above, the subtler performance difference among different optimized maps cannot be explained by the difference in their information density. So how can a map optimized on the mouse trajectory performs better still? We argue that this small but significant improvement is a second-order effect that requires a measure that depends on the detailed trajectory $\mathbf{s}$ explicitly. More specifically, a desirable measure could depend on the specific shape of the belief distribution right before an observation; if so, the specific path an agent take leading to the observation will contribute to the information gain. To sum up,

1) the current definition of information density is insensitive to the detailed trajectory, and

2) it does not explain the performance improvement of a map optimized on mouse trajectory over random trajectory

Based on these findings, we think that an alternative information-theoretic measure with a higher, yet bounded, order of dependence on the detailed trajectory could capture the performance improvement. And this is worth exploring in future work.

### 8. Other numerical schemes we do not implement and why

#### 8.1. Iteratively adding the most important edge

Another obvious choice among greedy optimizers is to iteratively adding an edge starting from the empty map. It could arguably to reach a better performance in certain situations, e.g., optimizing a map with very high sparsity with a high-fidelity path integrator. The reason we do not adopt this scheme is because it is incompatible with the chuck scheduler (discussed above) for speeding up optimization. Since the optimizer starts with the empty map where carefully adding an edge could greatly impact the performance, it is best to evaluate on the full trajectory in the beginning which happens to be the iteration with the longest candidate list for finding an edge to add. For that, the chunk scheduler can hardly speed things up.

#### 8.2. Pre-filtering candidate edges basing on belief instead of physical occupancy

When there is a tie in score among multiple candidate edges, we select the least physically visited edges to delete. It can be argued that an unvisited edge could still be important for belief distribution could still occupy the associated tile, and it would require encoding that "belief-occupied" edge in a map for ruling out this physically-unoccupied tile. Our preliminary results (not shown) show that this optimizer's performance is highly unstable, sensitive to hyperparameter tuning (e.g., random seed, numerical precision, etc.), and generally worse than our current optimization scheme. The precise reason on why such a unreliable performance requires more research beyond the scope of this paper. But we speculate that using the belief occupancy also suffers from the premature deletion issue we discussed earlier, where a physically unoccupied tile during a chunk could have a nonvanishingly small belief occupancy and thus being selected to delete.

#### 8.3. Simpler diffusion of belief distribution without heading

Although we adopt the minimalistic assumption about the path integrator—i.e., only heading integration is noisy, the derived belief diffusion ends up involve both heading and position variable. Consequently, it is less straightforward to implement and could be less intuitive in some cases (see example figures in the section Diffusion of belief distribution).

A simpler belief diffusion can be implemented to only propagate position variable: $P_t(s) \xrightarrow{\text{diffuse}} P_{t+1}(s)$ with a modified tile transition tensor from the original deterministic $T$ to a probabilistic one $\tilde{T} \equiv \alpha T + (1-\alpha) T_{\text{ring}}$ where $T_{\text{ring}}$ transition a tile $s_t$ to a ring of neighbors of the original destination $s_{t+1}$—i.e., $s_t \xrightarrow{a_t} [s_{t+1,\,0}, s_{t+1,\,1}, \ldots, s_{t+1,\,5}]$ as illustrated below

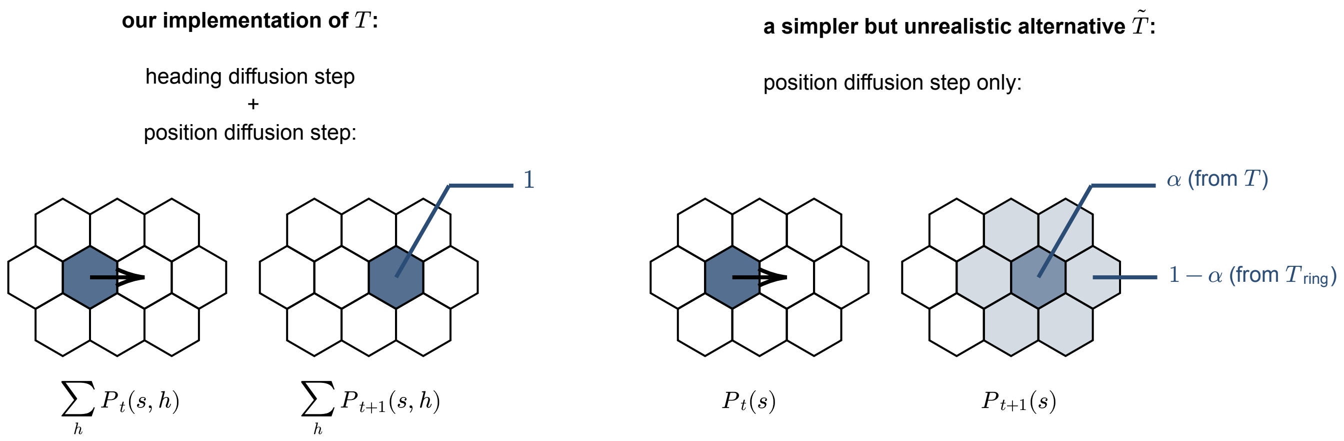

**Drawing 3:** Two different styles of noisy path integrator

This alternative version of belief diffusion step is equivalent to a uniform outward diffusion plus a ballistic movement along the agent's action. This seemingly simple implementation nonetheless makes very involved assumptions about noise model. It would require precise diffusion mechanisms for both heading ($\tilde{H}$) and distance integration ($\tilde{T}$). Even worse, it requires to fine tune these two supposedly independent mechanisms together, e.g., $\tilde{H} \mid \boldsymbol{\theta}_{H,T}$ instead of $\tilde{H} \mid \boldsymbol{\theta}_{H}$, in order to create such a uniform omni-directional diffusion while the entire distribution zigzagging across the whole arena without changing shape. Given that this alternative implementation is unlikely to resemble any real world situations, we favor our current implementation for its minimalistic assumption.

#### 8.4. Lossless boundary rule and alternative minimal map

As discussed above, the reason we implement a lossy boundary rule—where an out-of-boundary belief value is redistributed uniformly across all tiles equivalent to an information erasure—is because the alternative rule will give away unwanted information (or phantom information) about the boundary locations even when the agent has an empty map. Having discussed so, however, the lossy boundary comes with its own drawbacks. For the information erasure could be too extreme provided that those incorrect belief values would have stayed near the ground truth tile rather than suddenly spreading across the whole arena at the encounter of boundary. It's therefore worth discussing the alternative lossless boundary rule.

One way to embrace the unwanted boundary information is to store all the boundary tiles in a map from the start. One can thus use a boundary-only map to replace the original empty map as the default minimal map. In so doing, one focuses on the performance gain against this new baseline. The figure below, with a wall-only map, shows how much higher the performance is on a mouse trajectory than that on a random walk trajectory for high path integrator fidelity:

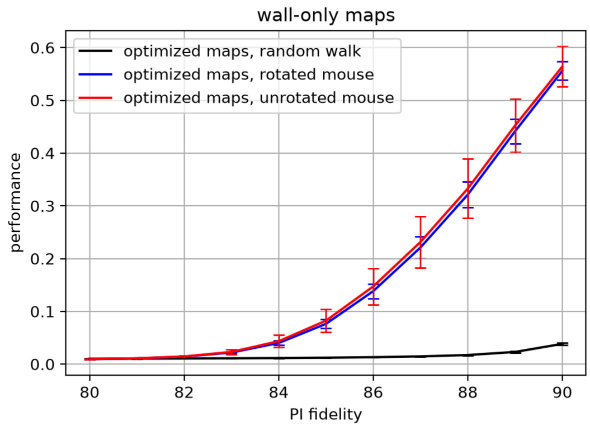

**Figure 9:** Mouse trajectories have unfair advantage against random walk due to disproportionate amount of wall following behavior. To study efficient map beyond simply encoding wall edges, we thus implement lossy boundary condition as discussed in section above.

From the above result, the reason for a mouse trajectory to have such high performance is because a much higher proportion of wall following behavior which causes a lot of phantom information gain. This is undesirable for our goal is to understand whether mice could use informative landmarks (not just the wall) to self localize. Yet with this minimal wall-only map, the performance is pretty much dominated simply by the amount of wall-following in a given trajectory. And this could make the subsequent statistical claims less straightforward, and the associated analysis much more involved.
